# A Wnt-specific astacin proteinase controls head formation in *Hydra*

**DOI:** 10.1101/2020.08.13.247569

**Authors:** Berenice Ziegler, Irene Yiallouros, Benjamin Trageser, Sumit Kumar, Moritz Mercker, Svenja Kling, Maike Fath, Uwe Warnken, Martina Schnölzer, Thomas W. Holstein, Markus Hartl, Anna Marciniak-Czochra, Jörg Stetefeld, Walter Stöcker, Suat Özbek

## Abstract

The *Hydra* head organizer acts as a signaling center that initiates and maintains the primary body axis in steady state polyps and during budding or regeneration. Wnt/beta-Catenin signaling functions as a primary cue controlling this process, but how Wnt ligand activity is locally restricted at the protein level is poorly understood.

Here we report the identification of an astacin family proteinase as a Wnt processing factor. *Hydra* astacin-7 (HAS-7) is expressed from gland cells as an apical-distal gradient in the body column, peaking close beneath the tentacle zone. *HAS-7* siRNA knockdown abrogates HyWnt3 proteolysis in the head tissue and induces a robust double axis phenotype, which is rescued by simultaneous *HyWnt3* knockdown. Accordingly, double axes are also observed in conditions of increased Wnt levels as in transgenic actin::HyWnt3 and HyDkk1/2/4 siRNA treated animals. *HyWnt3*-induced double axes in *Xenopus* embryos could be rescued by co-injection of *HAS-7* mRNA. Mathematical modelling combined with experimental promotor analysis indicate an indirect regulation of *HAS-7* by beta-Catenin, expanding the classical Turing-type activator-inhibitor model.

Our data suggest a negative regulatory function of Wnt processing astacin proteinases in the global patterning of the oral-aboral axis in *Hydra*.

## Introduction

Wnt ligands secreted from a local source at the posterior end of the embryo are conserved molecular cues for the patterning of the primary body axis in bilaterians and non-bilaterians [1–3]. The role of Wnt/beta-Catenin signaling in the axial patterning of cnidarians has been extensively studied in the freshwater polyp *Hydra* [4–7], which has a single oral-aboral body axis. The head is separated from the gastric region by a ring of tentacles and runs out at the upper part into a cone-shaped mouth region, called the hypostome. At its apical tip the hypostome contains the head organizer [8] comprising a small cluster of ecto- and endodermal cells that continuously express *HyWnt3* in steady state polyps (Fig. 1a) [9]. *HyWnt3* is upregulated early during head regeneration and has been shown to initiate a cascade of Wnt signaling events directing axial patterning [7]. While the spatially restricted HyWnt3 ligand production is controlled at the transcriptional level by repressive elements in the *HyWnt3* promotor region [9, 10], it is poorly understood how Wnt activity is regulated at protein level in the extracellular space. In *Hydra*, only a member of the Dkk1/2/4 family of secreted Wnt inhibitors has so far been shown to function as a Wnt antagonist by creating a Wnt-suppressed region in the body column [11]. Recently, we have shown that the matricellular protein Thrombospondin (HmTSP) is expressed directly from or in close vicinity of *HyWnt3* expressing cells of the hypostome and exerts a negative regulatory function on organizer formation [12]. It is unclear, though, whether HmTSP interacts directly with Wnt ligands or modulates Wnt inactivity by influencing receptor mobility or turnover.

**Fig. 1.**
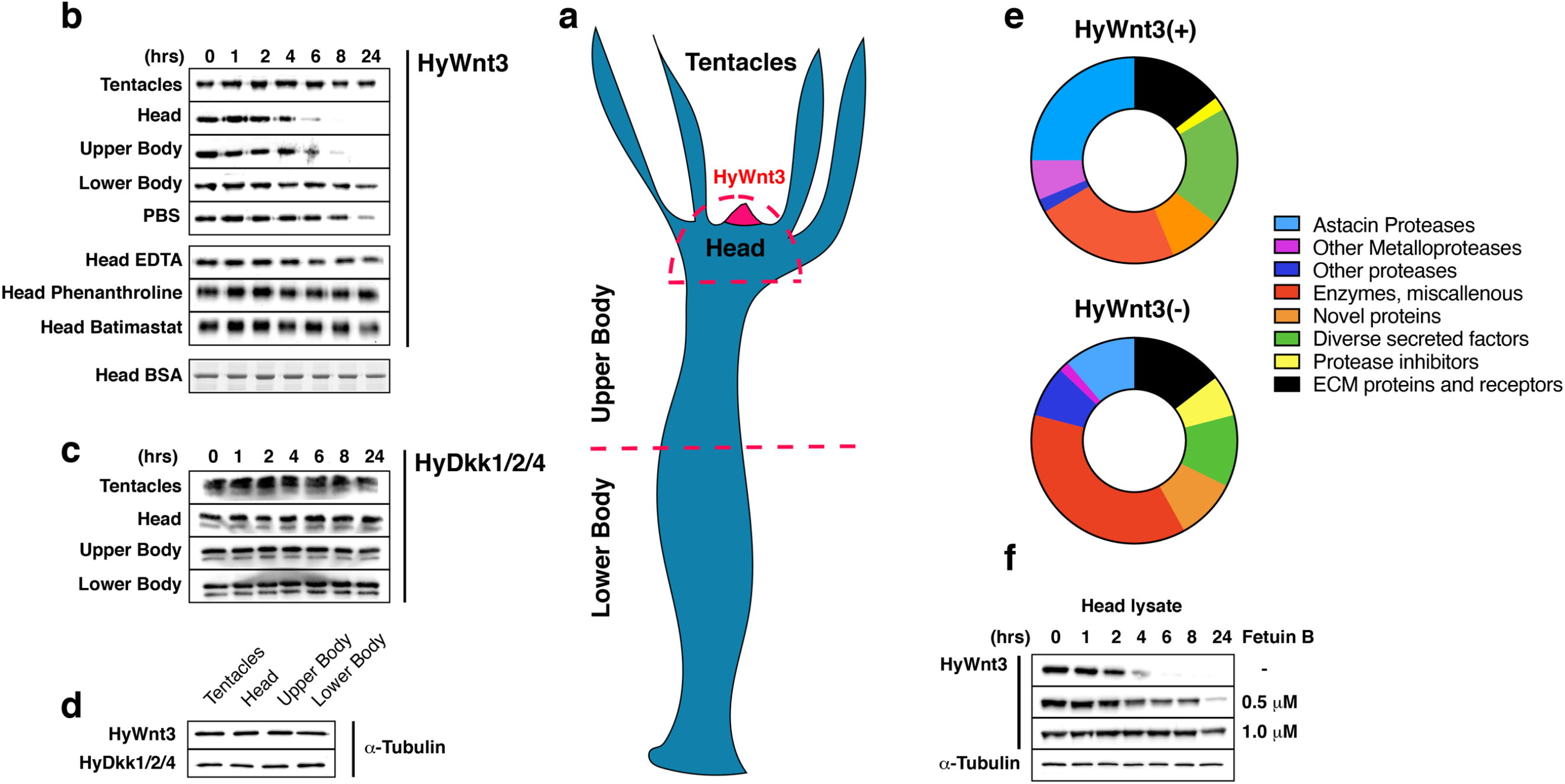
Screen for HyWnt3 proteolytic activity in *Hydra* tissue lysates. (a) Schematic representation of body plan. Body parts used for lysates in b-d are indicated. The hypostomal organizer, which harbors HyWnt3 expressing cells is marked in red. (b) Cleavage of recombinant HyWnt3-His, monitored by Western blotting with anti-His antibody, reached half-maximal levels after ~4 hrs incubation in head lysate and after ~6 hrs in upper body lysate. No cleavage was observed during incubation in tentacle and lower body lysates, while incubation in the PBS control showed unspecific cleavage at 24 hrs. No unspecific proteolysis on 1 μg BSA was detectable in HL over the time period of 24 hrs as detected by SDS-PAGE and Coomassie staining. HyWnt3-His cleavage activity in head lysate was completely blocked by the addition of broad zinc metalloproteinase inhibitors EDTA and Phenanthroline or the matrix metalloproteinase inhibitor Batimastat. (c) No cleavage was observed for the recombinant Wnt antagonist HyDkk1/2/4-His in the respective body tissue lysates during a 24 hrs incubation time. Mark that the double band appearance is an SDS-PAGE artifact. (d) Tissue lysates from different body parts of adult hydra polyps as indicated in the scheme were adjusted in total protein concentration by tubulin Western blotting. (e) Distribution of protein classes in *Hydra* head lysate secretome identified in HyWnt3(+) and HyWnt3(−) fractions as indicated. The full dataset is given in Additional file 4: Table S1a-b. (f) HyWnt3-His processing is inhibited by recombinant mouse Fetuin-B protein in a dose-dependent manner as indicated.

Morphogen activity during embryogenesis can also be restricted by proteinases that process secreted ligands. A prominent example is the zinc metalloproteinase BMP1 and its splice variant mammalian Tolloid (i.e. Xolloid in *Xenopus*), which specifically cleaves Chordin and thus promotes local BMP signaling at the ventral side of the vertebrate embryo [13]. A similar case for morphogen inactivation has been proposed for TIKI1, a highly conserved metalloproteinase expressed in the *Xenopus* organizer and shown to antagonize Wnt function by cleaving eight amino-terminal residues of Wnt3a [14].

In *Hydra*, functional studies on astacin metalloproteinases have indicated important roles in processes of morphogenesis and regeneration [15–17]. Yan et al. have shown that the metalloproteinase HMP1 is localized to the head pole and an anti-HMP1 antibody can effectively block head regeneration [17]. HMP2, a *Hydra* astacin proteinase of the meprin family, formed an opposing gradient to HMP1, showing highest expression at the basal pole of the animal [16]. Although different mechanistic pathways as the proteolytic activation of morphogens or regulatory peptides have been discussed in these studies, no detailed molecular mechanisms comparable to those for Tolloid or TIKI1 have been described so far for any cnidarian metalloproteinase.

Here, we identify a member of the astacin proteinase family in *Hydra* with Wnt3 processing activity. *Hydra* Astacin-7 (HAS-7) is expressed in an increasing gradient towards the tentacle base of the polyp, forming a ring-like zone between head and body column that shows upregulated expression for several other members of the astacin family. siRNA knockdown of *HAS-7* eliminates the HyWnt3 proteolytic activity of the head tissue leading to a robust double axis phenotype with a fully developed head structure. In addition, *HAS-7* mRNA injection into *Xenopus* embryos rescues double axes induced by *HyWnt3* mRNA. Our combined experimental data and mathematical models demonstrate a direct mechanistic link between astacin proteinases and Wnt-regulated pattern formation in *Hydra* by restricting Wnt ligand activity to the head region via specific proteolysis.

## Results and Discussion

### Identification of HyWnt3 proteolytic activity in *Hydra* head lysate

To identify factors restricting Wnt activity in the extracellular space we first examined the protein half-life of recombinant HyWnt3-His in tissue lysates generated from different body parts of *Hydra* (Fig. 1a). For this, lysates of the head region, tentacles, upper and lower body trunk were prepared and their soluble fractions were adjusted in total protein concentration to 4 mg/ml. ~10 ng of purified recombinant HyWnt3-His protein was incubated with equal amounts of each tissue lysate and the reaction was stopped after different time points. Detection by Western blotting located the highest proteolytic activity for HyWnt3-His in the head (half-life ~ 4 h) and, to a lesser extent, upper body lysates (half-life ~ 6 hrs) (Fig. 1b, d). While unspecific proteolysis of HyWnt3-His was evident after 24 hrs incubation in PBS, it stayed remarkably stable in lysates of tentacles and the lower body part. Incubation of 1 *μ*g BSA in head lysate (HL) did not show unspecific proteolysis over the given time period of 24 hrs (Fig. 1b). In HL samples supplemented with broad spectrum metalloproteinase inhibitors like EDTA and 1,10-Phenanthroline or the specific matrix metalloproteinase inhibitor Batimastat, HyWnt3-His processing was completely blocked in the given time frame, indicating that metalloproteinases could be responsible for the observed activity. A parallel experiment performed with recombinantly expressed HyDkk1/2/4-His protein, a major Wnt antagonist [11], showed no specific proteolytic activity for this factor in the respective lysates (Fig. 1c, d).

To isolate candidate factors involved in HyWnt3 processing we next used a proteomic approach. A pool of HL from 200 polyps was fractionated by cation exchange chromatography (Additional file 1: Fig. S1a) and peak fractions were re-examined for their HyWnt3-His processing activity applying a 6 hrs incubation time (Additional file 1: Fig. S1b). A fragment encompassing the two N-terminal cadherin domains of *Hydra* cadherin [18] was used as a control substrate to monitor general matrix metalloproteinase activity. We observed complete HyWnt3-His cleavage using fractions 1-5, while *Hydra* cadherin was degraded partially by fractions 2 and 3. To exclude a high background of possibly unspecific proteinases in fractions 1-3 we pooled fractions 4-5 (HyWnt3(+)) and 6-7 (HyWnt3(−)) for further analysis and performed orbitrap mass spectrometry analysis after in solution digestion of the respective pooled samples. When we filtered the obtained protein hits for unique sequences of proteins having a signal peptide for secretion and at least two peptide hits, astacin family proteinases constituted the largest group in the HyWnt3(+) secretome while miscellaneous enzymes dominated in the HyWnt3(−) fraction (Fig. 1e, Additional file 4: Table S1a-b, Additional file 5: Table S2). Of the 12 astacin sequences detected in the HyWnt3(+) fraction, 4 were also present in the HyWnt3(−) fraction, although with lower protein scores. The HyWnt3(−) secretome additionally contained an increased number of proteinases belonging to diverse families (Additional file 4: Table S1b, Additional file 5: Table S2). We concluded from these results that metalloproteinases, in particular astacin-type proteinases, are likely candidates for the observed HyWnt3-His processing activity. To confirm this notion, we tested the proteolytic activity of HL on HyWnt3 in the presence of recombinant mammalian Fetuin-B, which was recently shown to function as a highly specific physiological inhibitor of astacin-type proteinases like ovastacin [19]. As shown in Fig. 1f, murine Fetuin-B blocked HyWnt3-His processing by HL in a dose-dependent manner.

### Characterization of the HyWnt3(+) astacin secretome

The HL HyWnt3(+) secretome contained 12 unique astacin sequences (hence called *Hydra* Astacins, HAS) with HAS-1 and HAS-7 showing the highest protein scores in the orbitrap mass spectrometry analysis (Additional file 4: Table S1a, Additional file 5: Table S2). The alignment of the pro- and catalytic domains with known astacin proteinase amino acid sequences demonstrated a high conservation of critical sequence motifs as the aspartate switch residue, methionine turn, and zinc binding motif (Fig. 2a). The domain structure of astacins comprises a signal peptide and a variable pro-domain segment, which is cleaved to activate the central ~200-residue catalytic domain (Fig. 2a-b). Typical for cnidarian astacins is the possession of C-terminal ShKT (*Stichodactyla* toxin) domains [22]. The majority of the astacins detected in our analysis comprises 1-2 ShKT domains, but several lack a C-terminal segment (Fig. 2b). HAS-11 is exceptional in possessing six ShKT domains in a tandem repeat. None of the astacin sequences was predicted to possess a transmembrane domain. To be able to correlate the observed HyWnt3-specific proteolytic activity in the head and upper body tissues with expression patterns of the respective genes, we performed *in situ* hybridization experiments in whole mounts (WISH). Conventional WISH using mRNA antisense probes and sense controls was performed for the genes encoding the two highest-scoring protein hits, *HAS-1* and *HAS-7*, as well as for the previously described *HMP1* [16]. For the remaining astacin genes of the Wnt3(+) fraction, synthetic locked nucleic acid (LNA) probes were used that have an increased target specificity for highly similar gene sequences. ISH probe and LNA sequences are summarized in Additional file 6: Table S3. The majority of astacin genes showed an endodermal expression pattern in the upper half of the body trunk, often as a gradient that increased towards the head with a sharp boundary beneath the tentacle formation zone (*HAS-2*, *HAS-7*, *HMP1*, Fig. 2 c-d, f), reminiscent of the SEC-1 positive gland cell subpopulation described by Schmidt & David [23]. Several astacins (*HAS-5*, *HAS-9-11*) were expressed in a narrow collar demarcating the body region from the head (Fig. 2g-j). Four of the examined genes did not yield a reproducible staining pattern by WISH. *HAS-1* was exceptional in exhibiting a distinct expression in large gland cells in the endoderm that were homogenously distributed along the entire body column (Fig. 2e). This pattern is reminiscent of the Wnt antagonist *HyDkk1/2/4*, which is expressed by a subpopulation of zymogen gland cells [11]. By tracing the astacin genes of the HyWnt3(+) fraction in the *Hydra* single cell transcriptome database provided by the Celina Juliano lab (https://portals.broadinstitute.org/single_cell/study/stem-cell-differentiation-trajectories-in-hydra-resolved-at-single-cell-resolution) [24] we were able to identify cell-specific expression signatures for each sequence (Additional file 2: Fig. S2a-b). Scatter plots showing the expression profiles along cell types and differentiation trajectories clearly identify i-cell-derived gland cells distributed mostly along the transition zone between head and upper body (zymogen to mucous gland cell transition) as the source of all examined astacins, except *HAS-1*, which was restricted to zymogen gland cells of the body column. A close-up view of the epithelial layers in the upper gastric region of a *HAS-7* WISH sample confirmed that the mRNA signal is not present in the endodermal epithelial cells closely aligning the central line of extracellular matrix, called mesoglea (Additional file 2: Fig. S2c). Gland cells in *Hydra* are interspersed singly between the endodermal epithelial cells without direct contact to the mesoglea [23]. In summary, the candidate astacin proteinases identified in our analysis are a product of endodermal i-cell derived gland cells with an overlapping peak expression beneath the head and tentacle region. We focused our further analysis on HAS-7 due to its high protein score in the mass spectrometry analysis and the graded expression pattern reflecting the diminishing HyWnt3-His proteolysis from head to lower body (Fig. 1a). Also, six of the astacins identified in the HyWnt3(+) HL fraction were also contained in the HyWnt3(−) fraction (HAS-1-3, HAS-6, HAS-9, HAS-11), indicating that they are unlikely candidates for acting as HyWnt3-specific proteinases. Interestingly, none of the astacins identified in both fractions belongs to the group showing a graded apical-to-basal expression pattern.

**Fig. 2.**
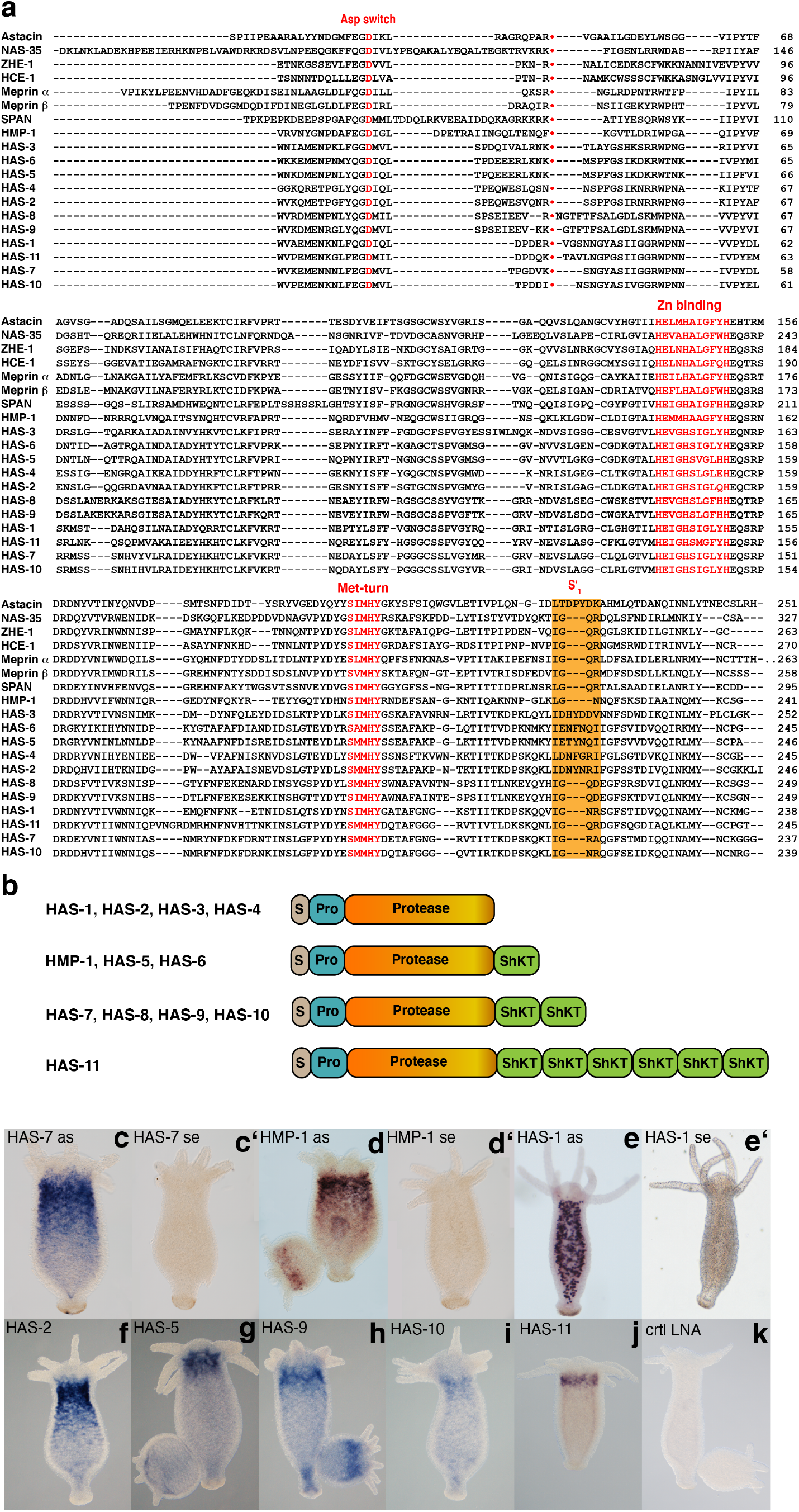
Sequence features and expression patterns of HyWnt3(+) astacin genes. (a) Multiple sequence alignment of pro-domain and catalytic domain sequences of astacins identified in this study. For comparison, astacin sequences from diverse species outside the cnidarian phylum were included. Gene ID numbers are as follows: Astacin *A. astacus* (P07584), NAS-35 *C. elegans* (P98060), ZHE-1 *Danio rerio* (Q1LW01), HCE-1 *O. latipes* (P31580), Meprin *α H. sapiens* (Q16819), Meprin *β H. sapiens* (Q16820), SPAN *S. purpuratus* (P98068), HMP1 (NP_001296695.1), HAS-3 (XP_002166229.3), HAS-6 (XP_002157397.2), HAS-5 (XP_002164800.1), HAS-4 (XP_002162738.1), HAS-2 (XP_002162822.1), HAS-8 (XP_002153855.1), HAS-9 (XP_002161766.1), HAS-1 (XP_012565441.1), HAS-11(XP_012561076.1), HAS-7 (XP_012560086.1), HAS-10 (XP_002159980.2). In red: the aspartate switch residue in the pro-peptide, the zinc binding motif and the Met-turn. Orange background: residues forming the S1 ′ sub-site. **•** denotes the activation site. (b) Domain structures of astacins detected in HyWnt3(+) head lysate fractions. S, signal peptide; Pro, pro-domain; ShKT, *Stichodactyla* toxin domain. (c-k) WISH experiments using antisense and sense oligonucleotide probes (c-e’) or LNAs (f-k) show a collar-like expression pattern marking a transition zone between head and body column for the majority of astacin genes. Representative of 10 hydras examined.

### *HAS-7* knockdown induces double axis formation

We next asked whether *HAS-7* depletion perturbs normal axis formation as a consequence of diminished HyWnt3 processing. siRNA gene expression knockdown by electroporation of adult polyps has recently been established as a robust method in *Hydra* [12, 25, 26]. We designed siRNAs targeting conserved motifs in the HAS-7 pro-domain (siRNA1), catalytic domain (siRNA2) and a less conserved region in the C-terminal ShKT domain (siRNA3) (sequences are given in Additional file 7: Table S4). To be able to monitor the electroporation efficiency in the endoderm we used an endodermal actin::GFP/ecdodermal actin::RFP transgenic strain (reverse watermelon) and applied siGFP in combination with target gene siRNAs [12, 25]. Although the knockdown phenotype for the strongly expressed GFP transgene is largely half-sided, we have evidence from previous studies that for native target genes the effect can be systemic [26]. To monitor the gene knockdown at protein level, we raised a polyclonal antibody against a unique epitope in the HAS-7 primary sequence located between the catalytic domain and the N-terminal ShKT domain (APPTAGPTISPT). A Western blot analysis of the different body part lysates used for the initial proteolytic assay showed a broad band migrating at ~40 kDa, which was strongest in the head tissue lysate (Fig. 3a). Upper and lower body tissues showed diminished signals reflecting the decreasing HyWnt3-His proteolytic activity observed in the respective body parts (Fig. 1b). The calculated molecular mass of full-length HAS-7 is 37.9 kDa, the mature proteinase lacking signal and pro-peptide sequences is predicted to have 33.3 kDa. Upper and lower body lysates showed additional bands at 55 and 70 kDa, which might be the result of a cross-reaction with a related epitope sequence, or represent a dimeric form of mature HAS-7 as described previously for meprins [27]. A recombinant histidine-tagged version of HAS-7 protein expressed in the insect cell line High Five could be detected by the specific Has-7 antibody (Fig. 3b, Additional file 3: Fig. S3). The apparent molecular mass of recombinant HAS-7 was slightly higher compared to the main band in the HL reflecting the additional amino acids of the tag sequence (1.7 kDa). siRNA knockdown with a combination of siRNAs 1/2 led to a moderate reduction of the 40 kDa band intensity as detected in whole body lysates (Fig. 3c). The combination of siRNAs 2/3 completely reduced band signals in this region and at the 70 kDa position. The 55 kDa band showed equal intensity in all conditions suggesting that it likely represents an unspecific protein. When we monitored HyWnt3-His half-life in head lysates of HAS-7 siRNAs 2/3 versus siGFP-treated animals, HyWnt3-His processing in head lysates after *HAS-7* knockdown was significantly impaired (Fig. 3d-e), suggesting that HAS-7 proteinase is responsible for the observed HyWnt3 cleavage in *Hydra*. The siRNA knockdown was additionally confirmed at the transcriptional level by quantitative RT-PCR analysis of head tissue from siGFP and siHAS-7 treated animals (Fig. 3f).

**Fig. 3.**
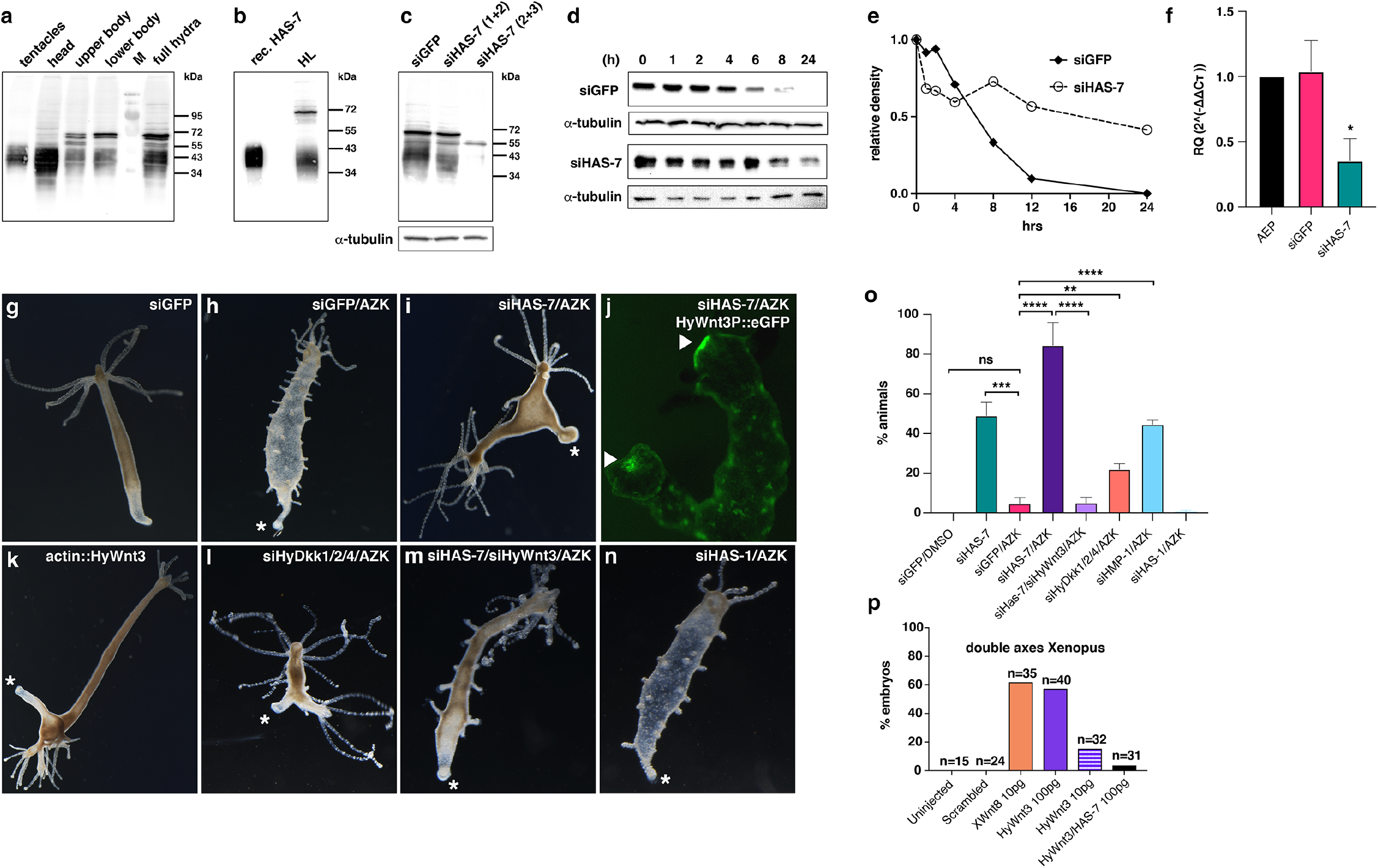
Functional analysis of *HAS-7* knockdown. (a) Detection of HAS-7 protein in body lysate samples using a polyclonal HAS-7 antibody. M = protein marker. (b) Recombinant HAS-7 protein expressed in High Five cells compared to native protein in head lysate as detected by HAS-7-specific antibody. Note that that the slightly higher apparent molecular mass is due to the introduced histidine tag of recombinant HAS-7. (c) Knockdown effect of different siRNA combinations on HAS-7 protein levels was assayed by anti-HAS-7 Western blot analysis of complete hydras treated as indicated. Tubulin was used as loading control of the respective hydra lysates. The distinct band at 70kDa in a-c likely represents a dimer of processed HAS-7. (d) HyWnt3-His proteolysis is impaired in head lysate of animals electroporated with siHAS-7 (2+3) as compared to siGFP control animals. Head lysates were generated 6 days after electroporation. Tubulin was used as loading control of the respective head lysates applied for each time point. (e) Relative intensities of the Western blot bands in d. (f) Quantitative real-time PCR analysis of HAS-7 expression in head tissues confirms the decreased expression in siHAS-7 treated animals compared to siGFP treated and untreated (steady-state AEP animals) controls. Results represent mean +/− S.D. from 3 independent experiments, analyzed by t tests. *p < 0.05. (g) siGFP control electroporation without AZK treatment shows normal morphology. (h) siGFP control animal showing ectopic tentacle formation after AZK treatment. (i) Double axis phenotype in hydras treated with AZK after HAS-7 (2+3)/GFP siRNA electroporation. Asterisk denotes the peduncle region. (j) Both heads in HyWnt3P::HyWnt3 transgenic animal treated as in h exhibit hypostomal HyWnt3 expression (arrow). Smaller spots along the body column represent ectopic organizers that usually give rise to ectopic tentacles as in h. (k) Double axis phenotype in actin::HyWnt3 transgenic hydra. (l) Double axis phenotype in hydras after HyDkk1/2/4/GFP siRNA electroporation and AZK treatment. (m) Rescue of double axis phenotype in animals treated with AZK after electroporation with a combination of HAS-7 (2+3) and HyWnt3/GFP siRNAs. (n) No double axes were observed in hydras treated with AZK after electroporation with HAS-1/GFP siRNA. (o) Ratios of double axis phenotypes in hydras after electroporation with siGFP or combinations of siGFP with siRNAs as indicated. In animals without subsequent AZK treatment double axes were counted 6 days after electroporation. In animals treated additionally with AZK, incubation was started 6 days after electroporation and the numbers of double axes in each group were counted 5 days after AZK removal. Animals (n) in each group were: siGFP/DMSO = 192, siGFP/AZK = 230, siHas-7/siGFP = 186, siHAS-7 /siGFP/AZK = 248, siHAS-7/siHyWnt3/siGFP/AZK = 203, siHyDkk1/2/4/siGFP/AZK = 290, siHMP-1/siGFP/AZK = 204, siHAS-1/siGFP/AZK = 150. Results from at least three independent experiments are shown. Each column represents the total percentage of one group, bars indicate the mean ± S.E.M. ****P value < 0.0001, ***P value < 0.0005, **P value < 0.001. ns = not significant. The data were analyzed using an unpaired parametric T-test with Welch’s correction followed by pairwise multiple comparisons of each group with the other groups. (p) Coinjection of HAS-7 mRNA inhibited HyWnt3 mRNA induced ectopic axis in *Xenopus laevis* axial duplication assay. *Xenopus* Wnt8 (XWnt8) mRNA injection served as positive control. Injected mRNA doses were as indicated. n indicates the number of embryos analyzed for each experimental condition.

In steady-state hydras, *HAS-7* mRNA depletion led to the induction of ectopic axis with a fully developed head structure in about 50% of the animals (Fig. 3i, o). To confirm that this is due to an increased Wnt signaling level we challenged electroporated hydras by treatment with AZK, a glycogen synthase kinase-3β inhibitor that leads to a systemic increase of beta-Catenin activity. This treatment normally induces ectopic tentacles along the gastric region 2-3 days after AZK removal emanating from small spot-like organizers that transiently express *HyWnt3* (Fig. 3h) [4]. siHAS-7 electroporated animals that additionally received an AZK pulse showed fewer ectopic tentacles, but formed double axes at a constant rate of 80-90% as compared to siGFP electroporated polyps, indicating that this phenotype dominates under conditions of increased basal levels of beta-Catenin activity (Fig. 3 h-i, o). A transgenic reporter line expressing eGFP under control of the *HyWnt3* promotor (HyWnt3P::eGFP) [9] confirmed that the ectopic head induced by *HAS-7* siRNA knockdown harbors a functional head organizer with hypostomal *HyWnt3* expression (Fig. 3j). Interestingly, a double and often multiple axis morphology was also observed in a transgenic strain that expresses *HyWnt3* under control of the actin promoter in all endodermal epithelial cells (Fig. 3k). Double axes were also obtained when animals were electroporated with siRNAs targeting the Wnt antagonist *HyDkk1/2/4* (Fig. 3l), although animals in this group regularly showed in addition multiple ectopic tentacles along the body column. In summary, depletion by siRNA knockdown efficiently reduced HAS-7 protein levels as well as HyWnt3-His proteolytic activity of the respective HL and resulted in an almost exclusive double axis phenotype that in AZK-treated animals. This is likely a consequence of unphysiologically increased HyWnt3 activity as similar phenotypes were observed by transgenic *HyWnt3* overexpression or *HyDkk1/2/4* depletion. This assumption was tested by combining *HyWnt3* and *HAS-7* siRNAs under AZK conditions, which resulted in a complete reversal of the double axis phenotype, although ectopic tentacles were still formed due to the increased beta-Catenin activity (Fig. 3m, o). Interestingly, *HMP-1* siRNA knockdown combined with AZK treatment also resulted in ectopic axis induction, but at a substantially lower level than for *HAS-7* (Fig. 3o), indicating some redundancy among astacins with similar expression patterns. No secondary axes were observed in animals electroporated with *HyDkk* and *HMP-1* siRNAs without subsequent AZK treatment. siRNA knockdown of *HAS-1*, which does not exhibit a graded apical-distal expression pattern, did not result in double axis formation after AZK treatment (n, o). To further confirm that HAS-7 is responsible for HyWnt3 inactivation *in vivo* we made use of the standard axial duplication assay in *Xenopus laevis*. Injection of 10 pg *XWnt8* mRNA in each blastomere of four-cell embryos induced secondary axis formation in more than 60% of the embryos compared to controls (Fig. 3p). Although *HyWnt3* injection induced secondary axes as well, higher mRNA doses (100 pg) were necessary to reach a comparable level to *XWnt8*. Co-injection of 100 pg *HAS-7* mRNA inhibited *HyWnt3*-induced secondary axis formation, indicating that *HAS-7* is sufficient to antagonize *HyWnt3* function in a heterologous setting.

### beta-Catenin dependency of *HAS-7* expression

The *HAS-7* expression zone, similar to the one of *HyDkk1/2/4* [11], is distinctly excluded from the region of Wnt expression in the hypostome area. This questions a direct beta-Catenin/TCF-dependent regulation of *HAS-7* as previously demonstrated for *HmTSP* that is expressed by *HyWnt3*-positive cells of the organizer [12]. When *HAS-7* gene expression was monitored by WISH following Alsterpaullone (ALP) treatment, we observed a global increase within the gastric region after 24 hrs, resulting in a loss of the original apical-distal expression gradient (Fig. 4a-b). 48 hrs after ALP wash, a line of increased *HAS-7* expression was evident close to regions of ectopic tentacle formation (Fig. 4c). In contrast to global activation by ALP, beta-catenin inhibition by iCRT14 treatment [28] did not alter the *HAS-7* expression pattern (Fig. 4e), arguing against a direct regulation by beta-Catenin/TCF. In transgenic actin::HyWnt3 *Hydra*, *HAS-7* was strongly upregulated throughout the gastric region (Fig. 4f), confirming that the observed increase of *HAS-7* expression induced by ALP is likely a consequence of global beta-Catenin activation. Interestingly, the head region was free of *HAS-7* mRNA in actin::HyWnt3 transgenic animals.

**Fig. 4.**
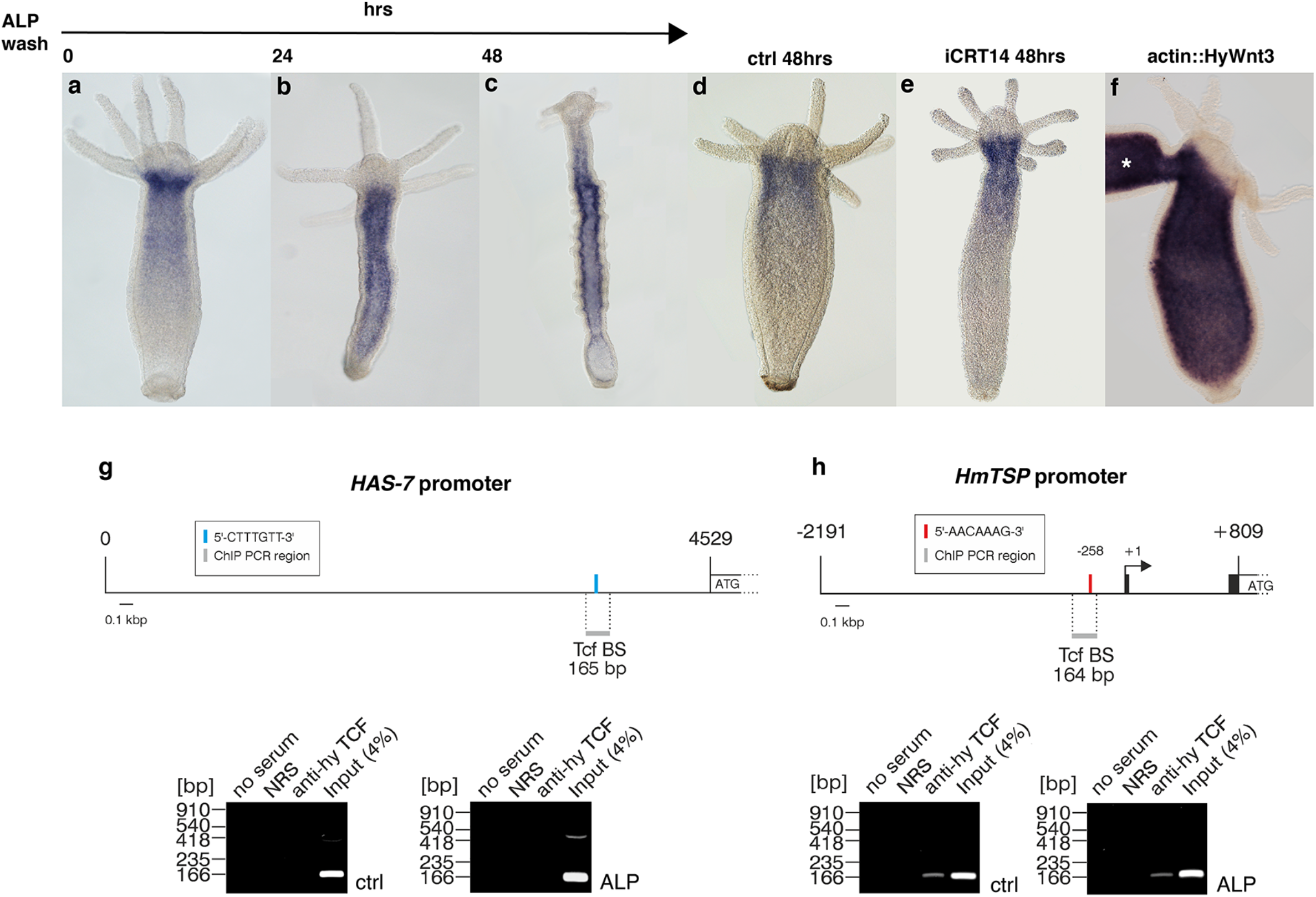
β-Catenin dependent expression of *HAS-7*. (a-c) ISH analysis of *HAS-7* expression after ALP treatment shows a global upregulation after 24 hrs and a shift towards the developing ectopic organizers along the body column after 48 hrs as compared to DMSO-treated controls (d). At 0h after ALP wash (a) no change of the *HAS-7* expression pattern compared to untreated controls (compare d and Fig. 2a) was evident. (e) Inhibition of beta-Catenin activity by iCRT14 treatment does not reduce normal HAS-7 expression levels. (f) *HAS-7* expression is globally upregulated in the gastric region of transgenic actin::HyWnt3 animals. The asterisk marks a secondary axis. Representatives of 10 hydras examined. (g-h) No detectable binding of *Hydra* TCF to the *HAS-7* promoter. (g) ChIP analysis of the *Hydra magnipapillata HAS-7* promoter. Upper site: Topography of the *HAS-7* 5’-untranslated region (nt 1 to 4529). The ATG indicates the translation start site. The position of a canonical TCF binding motif (5’-CTTTGTT-3’) is indicated by a blue bar. The localization of the 165-bp DNA segment flanked by the specific ChIP primer pair is visualized with a grey bar. Lower site: ChIP analysis of the *Hydra HAS-7* promoter region using chromatin from untreated whole hydra animals (ctrl), and from animals treated with ALP. A polyclonal antibody directed against *Hydra* TCF was used for precipitation, followed by PCR amplification of the indicated fragment from the *HAS-7* regulatory region. Reactions with normal rabbit serum (NRS) or total chromatin (Input) were used as controls. PCR products were resolved by agarose gel electrophoresis, and visualized by ethidium bromide staining. (h) ChIP analysis of the *HmTSP* promoter performed under the same conditions as in (g) and used as a positive control. Upper site: Topography of the 3,000-bp *HmTSP* promoter (nt −2191 to +809). Black boxes depict the first two exons of the *HmTSP* gene. The arrow indicates the transcription start site of the *HmTSP* mRNA, and ATG the translation start site. The position of the tested canonical TCF binding motif (5’-AACAAAG-3’) is indicated by a red bar. The localization and size of a 164-bp DNA segment flanked by specific the specific ChIP primer pair is visualized with a grey bar. Lower site: ChIP analysis as described under (g).

Upon examination of the *HAS-7* promoter region for regulatory elements, we identified a single putative TCF binding element with the conserved sequence motif 5’CTTTGTT3’ (Fig. 4g), similar to those experimentally confirmed in the *HyWnt3* [9] and *HmTSP* [12] promoters (CTTTGWW, W=A or T). To gain evidence whether *HAS-7* is directly regulated by beta-Catenin/TCF, we performed a chromatin immunoprecipitation (ChIP) assay using a *Hydra* TCF-specific antiserum as described previously [12, 29]. Chromatin was taken from control polyps and ALP-treated animals. No PCR product was obtained with primers flanking the identified TCF binding element in the *HAS-7* promoter in control and ALP conditions (Fig. 4g). In contrast, amplification of the TCF binding element in the *HmTSP* promoter yielded the expected PCR product from both samples (Fig. 4h). This suggests that TCF is not bound to the putative DNA binding region in the *HAS-7* promoter both in steady-state polyps and under conditions of global increase in nuclear beta-Catenin. We therefore presume that *HAS-7* is an indirect downstream target of beta-Catenin/TCF, which is positively regulated as part of the global Wnt signaling response, but is controlled by additional factors that down-regulate its expression in the vicinity of Wnt producing cells.

### HAS-7 function ensures single organizer dominance in an extended Turing model

To investigate the role of HAS-7 during axis and head pattern formation in *Hydra*, we built a mathematical model describing HAS-7-HyWnt3 spatio-temporal dynamics as an extension of the classical activator-inhibitor models of Gierer and Meinhardt [30–32]. We used the framework of a hypothetical two-component activator-inhibitor concept based on current experimental evidence of HyWnt3 and beta-catenin/TCF dynamics (see Additional file 8: Supporting Information S1 for details of the model and its biological justification) and employ our experimental findings concerning HAS-7. In particular, we extend the previous models by explicitly distinguishing between beta-catenin/TCF and HyWnt3 driven pattern formation. In addition, we verify how HAS-7 modulates formation of the body axes or the head in perturbed conditions (such as beta-catenin/TCF upregulation).

A scheme of the HAS-7-Wnt3 model is depicted in Fig. 5a. Based on our findings we postulate a negative feedback loop between HyWnt3 and HAS-7 (see Supporting Information for more details), representing one of the most frequent network motifs directing body-axis fine-scale segmentation in animals [33]. Additionally, we assume that *HAS-7* expression is placed downstream of the organizer/HyWnt3, and thus it is only indirectly positively affected by beta-catenin/TCF. In particular, the HyWnt3-HAS-7 interaction model comprises of two subsystems for organizer and axis formation, which act on different spatial scales. These include the small-scale HyWnt3-driven organizer pattern formation vs. beta-catenin/TCF signaling controlling large-scale body axis formation, as well as an additional subsystem directing tentacle formation (yellow blocks in Fig. 5a). The latter does not influence HAS-7-Wnt3 interactions. However, as it is affected by the body-axis and head system [30, 31], including it in the model makes possible additional model verification using, among other, data obtained under AZK treatment.

**Fig. 5.**
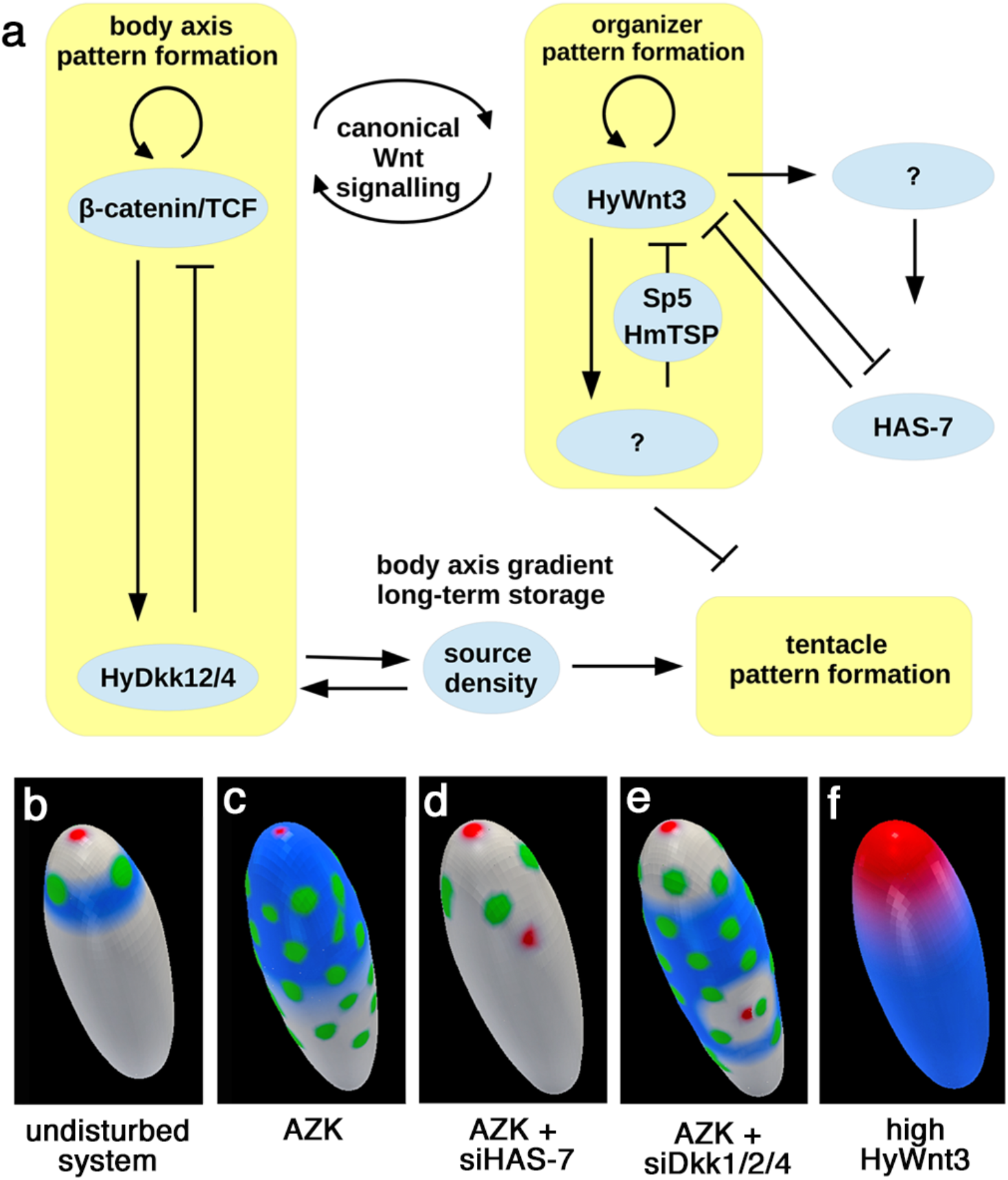
Mathematical model of HAS-7 function. (a) schematic representation of the model. Yellow blocks depict self-containing pattern formation systems consisting of an activator and an inhibitor. (b)-(f) finite element simulations of different experimental scenarios. Blue color corresponds to high levels of HAS-7 expression, red color to HyWnt3 expression, and green color indicates tentacle formation. The scale of these expression intensities is similar across all simulation plots.

As depicted in Fig. 5b, simulating the unperturbed system leads to the formation of a regular ring of tentacles below the hypostome. Furthermore, a single *HyWnt3* expression spot appears at the tip of the hypostome and *HAS-7* expression shows a distinct ring-like pattern. Similar patterns are observed in HAS-7 siRNA knockdown simulations without AZK treatment as modelling parameters were adjusted to preclude secondary axis formation in the unstimulated state (see SI file S1 for further details). Simulated AZK treatment assuming higher initial values of the source density [31] leads to multiple ectopic tentacles and elevated HAS-7 expression in the entire body column, but no additional axes (Fig. 5c). Combining the AZK treatment with *HAS-7* depletion (Fig. 5d) indeed leads to loss of ectopic tentacles and the development of a secondary axis, whereas AZK treatment combined with *HyDkk1*/2/4 depletion leads to formation of a secondary organizer including the formation of ectopic tentacles (Fig. 5e). Finally, *HyWnt3* overexpression (Fig. 5f) leads to a strong and body-wide expression of *HAS-7*. The only distinct difference between simulations and experimental observations is that a simulated *HyWnt3* overexpression does not lead to a multiple axis phenotype but to a broadened *Wnt3* expression domain at the head (Fig. 5f). The reason for this is that we describe each pattern formation system by the activator-inhibitor model, and an inherent mathematical property of this class of models does not allow changes in the activator production to cause an increase in the number of resulting heads (Marciniak-Czochra lab, in preparation). Thus, the applied activator-inhibitor model may be seen as an approximation of *de novo* pattern formation mechanism that is still unknown. In this context it is worth mentioning that *de novo* patterning may be based on not only chemical processes, but equally likely from mechanochemical, cellular and bio-electrical processes as well [34–36].

In summary, the proposed model qualitatively reproduces several of our experimental observations, including those which we did not even consider in the modelling process, such as the absence of ectopic tentacles in *HAS-7*-depleted animals treated with AZK. Analyzing the impact of model assumptions on the simulation results in the context of experimental data, we conclude that the function of HAS-7 is, most likely, to prevent the formation of multiple *HyWnt3* spots. In particular, the dominance of a single organizer is ensured by HAS-7-driven degradation of the canonical Wnt ligands. We assume that this task is especially important when beta-catenin/TCF levels are increased, which may occur in *Hydra* under natural conditions, for example in interaction with the environment [37], or as a response to injury [29].

### Substrate specificity of the HAS-7 proteinase

Altogether seven Wnt genes are expressed in the adult hypostome of *Hydra* (HyWnt1, −3, −7, −9/10a, −9/10c, −11, and −16) [7]. The size of their respective expression domains differs, with Wnt1, Wnt3, Wnt9/10a, c and Wnt11 showing a more confined spot-like expression reaching from the tip to the apical half of the hypostome, while Wnt7 and Wnt16 exhibit broader expression domains that extend to the tentacle bases. Wnt9/10b is not expressed in the adult polyp and Wnt5/8 are tentacle-specific [38]. Wnt2 is distinct in exhibiting a transitory expression at the site of bud initiation [7]. During head regeneration, the hypostomal Wnts are expressed in a temporally graded fashion with Wnt3, Wnt9/10c and Wnt11 being at the top of the cascade. Recombinant HyWnt3-His has been shown to induce head-forming capacity in tissue of the gastric region [7], indicating a primary role in initiating head formation. Our data indicate a redundant expression of several astacin proteinases in the sub-tentacle region with possibly similar substrate specificity. We have no evidence whether all head-specific Wnts are degraded by HAS-7 and closely related astacins, but considering that HyWnt3 acts at the top of the signaling cascade, it might serve as the primary substrate to inactivate the generation of a secondary organizer. When we screened the HyWnt3 amino acid sequence for putative cleavage sites for astacins, which were recently shown to have a strong preference for aspartate in the P1’ position (first amino acid C-terminal of the cleavage site) and proline in P2’ or P3’ in substrate peptides [20], only D187 followed by Pro188 fulfilled this criterion in HyWnt3. To evaluate if this is supported by the structural properties of the two proteins we created a model of the HyWnt3:HAS-7 complex using X-ray crystal structures of Wnt8 [39], crayfish astacin [40, 41] and zebrafish hatching enzyme (ZHE-1) [42] as templates (Fig. 6a-b). The N-terminal pro-domain, which confers protection of the active site is cleaved to transition the inactive zymogen into a catalytically potent astacin [22, 43, 44]. As a consequence, the alpha-helical N-terminal domain of HyWnt3 is able to approach the active site cleft of HAS-7 opposite to the lipid-mediated Frizzled receptor binding site [39] (Fig. 6a). The KDP motif of the putative cleavage site in HyWnt3 is positioned where the pro-domain cleavage site would reside. It is embedded in the deep active-site cleft, which like a horizontal patch spans the entire catalytic domain (Fig. 6b). The catalytic zinc ion resides at the bottom of the active site cleft and shows a slightly distorted octahedral coordinating sphere complexed with the imidazole moieties of His132, His136 and His142, respectively. HyWnt3 binds to HAS-7 in an extended conformation and is anchored to the proteinase cleft in an antiparallel manner. Astacins are the only extracellular proteinases with a primary specificity for negatively charged residues (preferably aspartate) in the position C-terminally of the cleaved peptide bond [20] in the protein target and the elongated KKRK*DPRKIM motif would allow for most efficient cleavage. The negatively charged aspartate side chain at the P1’ position is presumably bound by the positively charged arginine (R217) in the S1’ substrate binding site of HAS-7. In addition, a proline in close proximity in P2’ as given in the HyWnt3 sequence is frequently observed in astacin substrates [20, 45]. Mutation of the D187-P188 motif in HyWnt3 to alanine led to highly decreased secretion levels of the respective protein, indicative of protein instability or impaired folding in the secretory pathway, which precluded a meaningful cleavage assay.

**Fig. 6.**
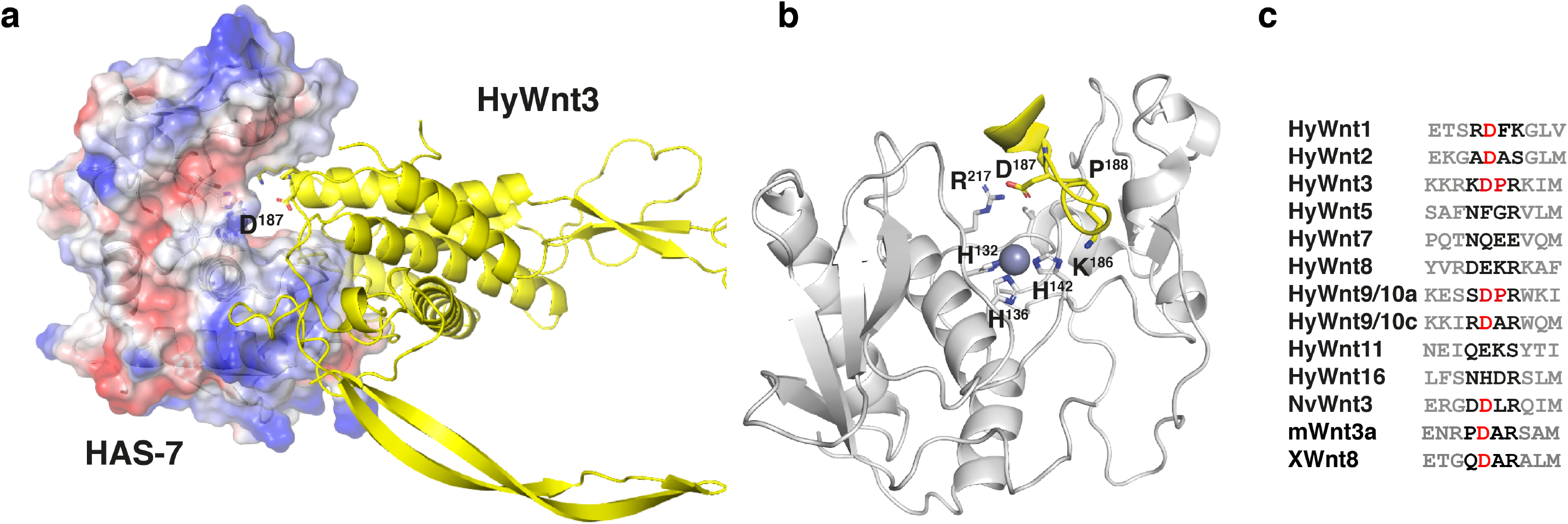
Structural model of the putative HyWnt3:HAS-7 complex. (a) Overview of HAS-7 (surface potential) complexed with HyWnt3 (yellow cartoon presentation). The active site Zn-ion of the astacin is shown as pink sphere. (b) Detailed view inside the active site pocket, shown in standard presentation with the Zn-ion located at the bottom of the cleft. (c) Alignment of amino acid sequences in different *Hydra* Wnt proteins encompassing the putative DP cleavage motif in HyWnt3.

Upon inspection of all *Hydra* Wnt sequences we found the DP motif at this position to be fully conserved only in HyWnt9/10a (Fig. 6c). Interestingly, non-canonical Wnts HyWnt5 and HyWnt8 [38] that are confined to the tentacle region and Wnts that have a broader expression domain in the head, like HyWnt7 and HyWnt16, were less conserved in this region and even lacked the aspartate residue. It is an attractive hypothesis that these Wnts act in confining the expression of HAS-7 and related astacins to the sub-tentacle region.

## Conclusions

Metalloproteinases have a broad spectrum of activities and can be involved in extracellular matrix remodeling but also in processing of signaling molecules in a variety of developmental events. Here, we show that a member of the cnidarian astacin proteinase family acts as a negative feedback regulator of Wnt3 activity by direct morphogen processing, thereby providing a limitation of high canonical Wnt activity to the hypostomal organizer. Given the complex expression pattern of Wnt ligands in *Hydra’s* head organizer, it will be interesting to unravel possible fine-tuned substrate specificities of the seemingly redundant astacin proteinase presence insulating the head region.

## Methods

### Animals

*Hydra vulgaris* was used for all experiments. For siRNA experiments, a transgenic *Hydra vulgaris* strain expressing endodermal GFP and ectodermal RFP (reverse water melon) provided by Robert Steele’s laboratory was used as described before [25]. The transgenic reporter line expressing eGFP under control of the Hydra Wnt3 promotor was described by Nakamura et al. [9]. The transgenic vector used to generate the strain expressing HyWnt3 under control of the actin promotor was produced by cloning the full-length HyWnt3 sequence between the actin promotor and actin 3’ flanking sequences of the pBSSA-AR vector using NcoI and XbaI sites [9]. Generation of the transgenic actin::HyWnt3 line was performed by microinjection as described before [9]. All animals were maintained in artificial *Hydra* medium (HM, 1 mM CaCl_2_, 0.1 mM MgCl_2_, 0.1 mM KCl,1 mM NaH_2_CO_3_, pH 6.8) at 18 °C in polystyrene dishes (Carl Roth) and fed two to three times per week with freshly hatched *Artemia salina* nauplii, unless indicated otherwise. Media was renewed 3–4 hrs after feeding and again the following day. Animals were starved for 24 hrs prior to experiments, unless indicated otherwise.

### Wnt proteolysis assays using tissue lysates

*Hydra* Wnt3 cDNA lacking the native leader peptide were subcloned into the pCEP-Pu mammalian expression vector, which introduces a BM-40 signal sequence and a C-terminal histidine tag as described previously [12]. Recombinant HyWnt3 was expressed using transiently or stably transfected HEK293T cells. For this, cells were seeded in full DMEM Medium (Gibco) in 6-well plates and transfected at 70% confluency using 1 μg of the respective vector construct together with 5 μl transfection reagent (TransIT^®^-LT1 Reagent, Mirus Bio) in 200 μl serum-free DMEM medium after 20 min incubation time. The cells were maintained in a humidified 5% CO_2_ atmosphere at 37 °C. After 48 hrs the medium was harvested, followed by Ni-NTA sepharose (Qiagen) batch purification. For 1 ml conditioned medium, 40 μl Ni-NTA beads were added and incubated for 1 hr at 4 °C under constant rotation. After a 2 min centrifugation step at 500 rpm the beads were washed two times with PBS. Elution was performed with 250 mM Imidazol for 15 min at RT. The beads were centrifuged at 500 rpm and the supernatant containing the histidine-tagged Wnt protein was instantly frozen in liquid nitrogen and stored at −80 °C. The eluted *Hydra* Wnt3 was verified by Western blotting using 10% SDS-PAGE gels. The proteins were transferred to PVDF membranes by wet blotting. Membranes were blocked for 1 hr at RT in PBS containing 5% BSA and 0.2% Tween-20 (PBST), incubated with mouse Penta His antibody (Qiagen) at 1:1000 in 1% BSA at 4 °C overnight, washed 3 x 5 min with PBST and incubated with anti-mouse horseradish peroxidase-conjugated antibody at 1:10000 in PBST containing 5% BSA for 1 hr at RT. The membrane was washed 2 x 5 min with PBST and 2 x 5 min with PBS and blots were developed using a peroxidase substrate for enhanced chemiluminescence.

To generate *Hydra* body part lysates, 100 hydras were cut into four parts using a scalpel. Tentacles, head without tentacles, upper and lower gastric region were immediately transferred on ice, sonicated in 250 μl ice cold PBS, centrifugated at 10000 rpm for 1 min at 4 °C, snap frozen and kept in aliquots at −80 °C. Body part lysate concentrations were adjusted to 4 mg/ml protein concentration using a nanodrop photometer and verified by Western blotting with alpha-tubulin antibody (Sigma-Aldrich). For the proteolysis assay, ~10 ng of purified HyWnt3-His was incubated with 15 μg of the respective tissue lysates in a total volume of 12 μl in PBS for 0, 1, 2, 4, 6, 8 and 24 hrs at RT. Peak fractions from the ion exchange chromatography were incubated for 6 hrs. For inhibitor treatments, 1, 10-Phenanthroline (Sigma-Aldrich), Batimastat (Sigma-Aldrich) or EDTA (AppliChem) were added at 200 μM, respectively. Fetuin-B was diluted in PBS and incubated for the given time periods as indicated in Fig. 1f) The incubation was stopped by adding SDS-PAGE sample buffer (10% glycerol, 50mM Tris-HCl pH 6.8, 0.02% bromophenol blue, 2% SDS, 1% 2-mercaptoethanol) and heating at 96 °C for 5 min. HyWnt3-His band intensities were documented by immunoblotting as described above. Unspecific proteolysis in HL was monitored adding 15 μg of the lysate to 1 μg of BSA (Roth) in a total volume of 12 μl in PBS and incubation at RT. The reaction was stopped by adding sample buffer and heat denaturation at 96 °C for 5 min after the same time periods as for HyWnt3-His. Samples were analyzed using 10% SDS-PAGE gels and Coomassie staining.

Full-length recombinant HyDkk1/2/4-His protein was expressed in *E. coli* BL21 (DE3) cells from a pET15b vector (Novagen) and purified under native conditions from the supernatant of cell pellets lysed by several freeze/thaw steps in PBS. The cleared supernatant was filtered and purified using Ni-NTA agarose beads (Qiagen). The eluted protein solution was dialyzed against PBS and purity was checked by SDS-PAGE. For the proteolysis assay, 10 ng of purified HyDkk/1/2/4-His was incubated with 15 μg of the respective tissue lysates in a total volume of 12 μl in PBS for 0, 1, 2, 4, 6, 8 and 24 hrs at RT. HyDkk1/2/4-His band intensities were documented by immunoblotting as described above.

The sequence for the *Hydra* Cadherin CAD1-2 domain (15-239) was amplified by PCR using a large 5’ cDNA fragment of *Hydra* Cadherin as template. The PCR product was cloned into the pET19B vector (Novagen) using NdeI and BamHI sites and the recombinant protein was expressed in *E. coli* BL21 (DE3) cells. Inclusion bodies were solubilized in PBS, 8 M urea and bound to Ni-NTA agarose beads. Refolding was performed by changing to PBS buffer prior to elution with PBS, 250mM imidazole, 0.4 M L-arginine (AppliChem). The eluted protein solution was dialyzed against PBS and purity was checked by SDS-PAGE. 10 ng of the purified protein was used as substrate with 5 μl of either the HyWnt3(+) and HyWnt3(−) HL fraction. Detection of the *Hydra* cadherin fragment by immunoblotting was performed as described above.

The polyclonal HAS-7 specific antibody was raised against a peptide corresponding to a sequence between the catalytic and ShKT domains (CKGGGNPPTGPPTAPP). For Western blotting, a PVDF membrane was blocked in 5% skimmed milk powder in PBS, 0.2% Tween-20, for 1 hr at RT. The primary antibody was applied at 1:1000 in 1% skimmed milk powder in PBST and membranes were incubated overnight at 4 °C. A peroxidase-conjugated anti-rabbit antibody (Jackson ImmunoResearch Laboratories, Inc.) was applied in blocking solution at 1:10000 for 1 hr at RT. Quantification of Western blot band intensities was performed using ImageStudioLite software.

### Ion exchange chromatography and mass spectrometry analysis

A pool of head lysate was generated by decapitating 400 animals closely beneath the head region and removing tentacles using a scalpel. Head tissue pieces were resuspended in ice cold 20 mM Tris-HCl, pH 7.5, and lysed on ice by passing at least 10 times through a 20-gauge needle. The lysate was centrifuged for 15 min (14000 rpm) at 4 °C and the supernatant was applied to anion exchange chromatography using a MonoQ HR5/5 column (GE healthcare). The eluent buffer contained 1 M NaCl in 20 mM Tris-HCl, pH 7.5. Peak fractions indicated in Additional file 1: Fig. S1 were screened for HyWnt3-His proteolytic activity as described above and frozen at −20 °C until further use. Pooled HyWnt3(+) and HyWnt3(−) fractions were subjected to mass spectrometry analysis after in solution tryptic digestion. Peptide separation was achieved using a nano Acquity UPLC system (Waters). Protein mass spectrometry analyses were performed as previously described [46]. In short: The nano UPLC system was coupled online to an LTQ OrbitrapXL mass spectrometer (Thermo Fisher Scientific). Data dependent acquisition with Xcalibur 2.0.6 (Thermo Fisher Scientific) was performed by one FTMS scan with a resolution of 60000 and a range from 370 to 2000 m/z in parallel with six MS/MS scans of the most intense precursor ions in the ion trap. The mgf files processed from the MS raw files were used for protein database searches with the MASCOT search engine (Matrix Science, version 2.2) against NCBI GenBank Proteins (version of October 28, 2015). Domain composition of GenBank protein accessions was analyzed against CDD (NCBI) and InterProScan 5 (EMBL-EBI).

### Expression and purification of recombinant HAS-7

Expression of HAS-7 occurred in insect cells using the Bac-to-Bac system (Invitrogen, Thermo Fisher Scientific). A synthetic fragment of the HAS-7 full-length cDNA (Biomatik) was subcloned into the pCEP-Pu mammalian expression vector, which introduces a C-terminal histidine tag and the fragment containing the C-terminal His_(6)_ - tag was inserted into the donor plasmid pFASTBac1 to generate bacmids in *E. coli* DH10Bac cells for transfection of insect cells according to the manufacturer’s manual. Amplification of recombinant baculoviruses occurred in *Spodoptera frugiperda* 9 cells (Sf9 CCLV-RIE 203, Friedrich–Löffler Institute, Greifswald, Germany) growing as adherent monolayer in Grace’s insect medium supplemented with 100 U/ml penicillin and 100 μg/ml streptomycin. For expression of HAS-7-His_(6)_ *Trichoplusia ni* cells (BTI-TN-5B1-4/High Five-cells CCLV-RIE 305, Friedrich–Löffler Institute, Greifswald, Germany) were cultured as suspension in Express Five SFM containing 100 U/ml penicillin, 100 μg/ml streptomycin and 16.4 mM L-Glutamine. Infected High Five cells were incubated for 72 hrs at 27 °C in Fernbach flasks (shaking incubator Multitron, INFORS). Following centrifugation, proteins were precipitated at 10 °C by step wise addition of solid ammonium sulfate to the supernatant resulting in a 60% ammonium sulfate saturation. After further stirring over night at 10 °C and centrifugation (9000 x g, 90 min, 10 °C) the harvested pellet was resuspended in 1/10 of the expression volume in 50 mM Tris-HCl pH 7.6, 300 mM NaCl, 20 mM imidazole and dialyzed against the same buffer. The cleared supernatant gained after centrifugation (8000 x g, 10 min, 4 °C) was applicated onto a Ni-NTA column (Qiagen). Following several washing steps with 50 mM Tris-HCl pH 7.6, 300 mM NaCl containing increasing imidazole concentrations (20 mM, 50 mM and 100 mM), the protein was finally eluted in buffer containing 250 mM imidazole. The elution fractions were pooled, dialyzed against PBS (140 mM NaCl, 2.7 mM KCl, 10 mM Na_2_HPO_4_ 2H_2_O, 1.8 mM KH_2_PO_4_, pH 7.4) and concentrated (Millipore Amicon Ultra, 3 K). SDS-PAGE and transfer to PVDF was performed as described previously [47]. The membrane was blocked in 3% BSA in TBS for 2 hrs, incubated for 1 hr with Penta-His antibody at 1:2000 and further incubated for 1 hr with secondary antibody (goat anti mouse POX 1:7500 in 7.5% skimmed milk powder in TBS). After each antibody treatment three washing steps were inserted (2 x TBST, 1x TBS). The Clarity Western ECL Substrate (Biorad) was used for detection.

### *In situ* hybridization

Customized LNA digoxygenin-labeled RNA probes were designed and produced by Qiagen corresponding to the antisense strands of the respective astacin cDNAs (sequences are given in Additional file 6: Table S3). The whole-mount ISH procedure was performed as described previously [48]. For hybridization, the LNA probe was added to a final concentration of 1 μM in fresh hybridization solution (1:1 mixture of deionized formamide and buffer containing 5x SSC (750 mM NaCl, 75 mM sodium citrate), 0.2 mg/ml yeast tRNA, 2% of 50x Denhardt’s solution, 0.1 mg/ml heparin, 0.1% Tween-20 and 0.1% CHAPS) and hybridized for ~60 hrs at 55 °C. Digoxygenin-labelled RNA probes for *HAS-1*, *HAS-7* and *HMP-1* corresponding to the sense and antisense strands were prepared using an RNA labelling *in vitro* transcription kit (Roche). ISH probe sequences covered the predicted full-length mRNA sequence for each gene. The further procedure was as described previously [12]. Samples were finally mounted in 90% glycerol in PBS or in Mowiol 4–88 (Carl Roth) and images were acquired with a Nikon Digital Sight DS-U1 camera mounted on Nikon Eclipse 80i and imaging software NIS Elements (3.10, SP3, Hotfix, Build645). Further image processing was performed with Adobe Photoshop CS6 and Fiji.

### Alsterpaullone and iCRT14 treatment for ISH analysis

80 budless hydras, which were fed the day before, were incubated in 5 μM ALP (Sigma-Aldrich) in DMSO in 100 ml HM. The animals were kept in the dark at 18 °C for 24 hrs. After incubation, the HM was changed every day. Samples were taken for ISH at 24 hrs, 48 hrs and 72 hrs after ALP treatment. Hydras incubated for 24 hrs in DMSO and fixed after 72 hrs served as control. For the β-catenin inhibitor experiment, 30 budless hydras, which were fed the day before, were collected in 10 ml petri dishes filled with HM. iCRT14 (Sigma-Aldrich) in DMSO was added to the petri dishes at a final concentration of 10 μM. The petri dishes were incubated in the dark at 18 °C. After 48 hrs samples were taken for WISH.

### Electroporation with siRNAs

siRNAs (HPLC grade) (siGFP, scrambled siGFP, siHAS-7, siHMP1, siHyDkk1/2/4, siHyWnt3, see Additional file 7: Table S4 for sequences) were purchased from Qiagen. Electroporation of siRNA was performed as described recently [12] using 3 μM of siGFP (1 μM siGFP and 2 μM scrambled siGFP) or a combination of siGFP and target siRNAs (1 μM each). Immediately after the pulse, 500 μl restoration medium consisting of 80% HM and 20% hyperosmotic dissociation medium (6 mM CaCl_2_, 1.2 mM MgSO_4_, 3.6 mM KCl, 12.5 mM TES, 6 mM sodium pyruvate, 6 mM sodium citrate, 6 mM glucose and 50 mg/l rifampicin, 100 mg/l streptomycin, 50 mg/l kanamycin, pH 6.9) was added to the cuvette, the animals were transferred to Petri dishes containing restoration medium and allowed to recover for one day. Viable polyps were transferred to new dishes containing HM and maintained under standard culture conditions. For AZK treatment, animals were incubated at 6 days post-electroporation either with 0.1% DMSO or 50 nM AZK in HM for 16 hrs. Thereafter, they were rinsed in HM several times and cultured under standard conditions. 5 days after AZK treatment, the animals were anesthetized in 1 mM Linalool (Sigma-Aldrich) [49] using a Nikon SMZ25 stereomicroscope equipped with Nikon DS-Ri2 high-definition color camera. Note that AZK instead of ALP was used in siRNA experiments to be able to reduce the total inhibitor concentration applied on the electroporated animals. 50 nM AZK induces the same grade of ectopic tentacles as 5 μM ALP when applied in the standard assay published previously [38].

### Chromatin immunoprecipitation

Chromatin immunoprecipitation analysis was carried out as described recently by using sheared extracts from formaldehyde-treated *Hydra* animals, which have been treated without or with 5 μM ALP (48hrs) and an antiserum directed against a recombinant *Hydra* TCF protein [12, 29]. PCR of precipitated DNA was done using specific primers flanking the potential TCF binding sites in the 5’-regulatory regions of the *Hydra HAS-7* or the *TSP* gene (Fig. 4g-h). PCR primer sequences: TCF binding motif in *HAS-7* (5’-GCTGTTATCTGTCCGCTTTC-3’/5’-CCATATAGAGGCCACACACC-3’), and the proximal TCF binding motif in *TSP* (5’-TTGAAGGCATTTAACAACTTGC-3’/5’-TGCCCAAATGTAAAGTTCTGTG-3’).

### Real-time Quantitative PCR

The RNA isolation was performed as described previously [46]. 60 hydra heads per condition were isolated either from steady state AEP polyps or siRNA treated transgenic reverse water melon animals by removing the tentacles and gastric region using a scalpel. The heads were immediately transferred into 200 μl TRIzol and stored at −20 °C. For siRNA (siGFP or siHAS-7) treated hydras, head samples were isolated six days after electroporation. cDNA synthesis was performed using the SensiFAST cDNA Synthesis Kit (Bioline) according to manufacturer’s instructions. RT-qPCR was carried out with a StepOnePlus ™ instrument (Applied Biosystems, Thermo Fisher Scientific) using the SensiFAST SYBR-Hi-ROX Kit (Bioline) according to the manufacturer’s instructions. The transcript level analysis was done by the ΔΔC(T)-Method with Elongation Factor 1-α (EF-1α) as a house keeping gene for normalization. Three biological replicates were performed for each experiment and triplicate measurements were made for each sample in each experiment. No template conditions served as negative controls. The data are presented as relative quantity (RQ) by 2^(-ΔΔC(T)) calculation. qPCR primer sequences are given in Additional file 7: Table S4.

### *Xenopus* experiments

*In vitro* fertilization, embryo culture and culture of explants were carried out as described [50]. Staging was done according to Nieuwkopp [51]. mRNA was produced with the mMessage mMachine SP6 trancription Kit (Ambion) from the HyWnt3, XWnt8, HAS-7, flag-tagged GFP (control mRNA) ORFs in the respective linearized pCS2+ vectors. mRNA was purified with a phenol/chloroform extraction and a subsequent isopropanol precipitation. Injections were done into the marginal zone of both ventral cells of the 4-cell stage. Total amounts of each injected mRNA were: XWnt8 (10 pg), HyWnt3 (10 or 100 pg), HyWnt3/HAS-7 (100 pg each), GFP control (100 pg).

### Structural modeling

Protein-Protein docking experiments were performed in ClusPro2 [52] using the crystal structures of *Xenopus* Wnt8 (pdb-code:4F0A) and promeprin ß (pdb-code: 4GWM). The propetide E25-G66 blocking access to the active site cleft and disordered loop segments in Wnt8 were removed to allow for proper structural analysis. Targeted search matrices were chosen by selecting the zinc-binding site of promeprin ß (H132-H136-H142) and the putative cleavage site of HyWnt3 (K186-D187-P188) as attractive search targets. Based on Lennart-Jones potentials, distance metrices and electrostatic evaluations the best 100 docking hits were subject to gradient energy minimization in the Crystallography and NMR system. 14 The lowest energy structures were further subject to 500 cycles of unrestrained Powell minimization. Harmonic restraints were imposed on the target molecule (2 kcal/mol Å^2^) with increased weight (25 kcal/mol Å^2^). Protein structure and model assessment tools were used to verify the quality of the modeled structure. Additionally, HAS-7 modelling was performed using Modeller [53] implemented in Chimera [54] with astacin complexed to a transition state analogue inhibitor (pdb-code: 1QJI) and zebrafisch hatching enzyme (3LQB), the latter being the most closely HAS-7-related astacin with known structure to date.

## List of abbreviations

ALP: Alsterpaullone
AZK: 1-Azakenpaullone
BMP: bone morphogenetic protein
ChIP: chromatin immunoprecipitation
DFG: Deutsche Forschungsgemeinschaft
Dkk: Dickkopf
DMEM: Dulbecco’s modified Eagle’s medium
DMSO: dimethylsulfoxid
FTMS: Fourier transform mass spectrometry
GFP: Green fluorescent protein
GSK-3: glycogen synthase kinase 3
HAS: *Hydra* astacin
HPLC: high performance liquid chromatography
HL: head lysate
Hm: *Hydra magnipapillata*
HM: Hydra medium
HMP: Hydra metalloproteinase
iCRT: inhibitor of β-catenin-responsive transcription
ISH: *in situ* hybridization
LNA: locked nucleic acid
ORF: open reading frame
PBS: phosphate-buffered saline
PCR: polymerase chain reaction
PDB: protein data bank
RFP: red fluorescent protein
RT: room temperature
SEC: Secretome
ShkT: Stichodactyla toxin
siRNA: short interfering RNA
TCF: T-cell factor
TES: 2-[Tris-(hydroxymethyl)methylamino]-1-ethane sulfonic acid
TSP: Thrombospondin
Wnt: wingless/integrated
UPLC: ultra-high-performance liquid chromatography
WISH: whole mount *in situ* hybridization
X: *Xenopus*.

## Declarations

### Ethics approval and consent to participate

*Xenopus* experiments were approved by the state review board of Baden-Wuerttemberg, Germany (License number: G-13/186) and performed according to the federal and institutional guideline.

### Authors’ contributions

SÖ, BZ, WS and IY designed experiments. BZ, BT, SK, MF, SK and IY performed experiments. JS and WS performed the structural modeling. UW and MS performed the mass spectrometry analysis. MH designed and performed the ChIP analysis. TWH provided the HyWnt3 transgenes. MM and AMC performed the mathematical modeling. SÖ, BZ, MM, WS and TWH wrote the manuscript. All authors read and approved the final manuscript.

## Acknowledgements

We thank Yukio Nakamura for the preparation of the actin::HyWnt3 pBSSA-AR vector, Jutta Tennigkeit for preparation of the recombinant expression constructs, Ann-Kathrin Heilig for performing *in situ* hybridization experiments, Alexander Hirth and Christof Niehrs for sharing *Xenopus* embryos and assisting with mRNA injections. We thank Kane Puglisi for technical assistance in the ChIP analysis.

