## Supplemental Files for "A Wnt-specific astacin proteinase controls head formation in *Hydra*"

**Supplemental file 1: Fig. S1**

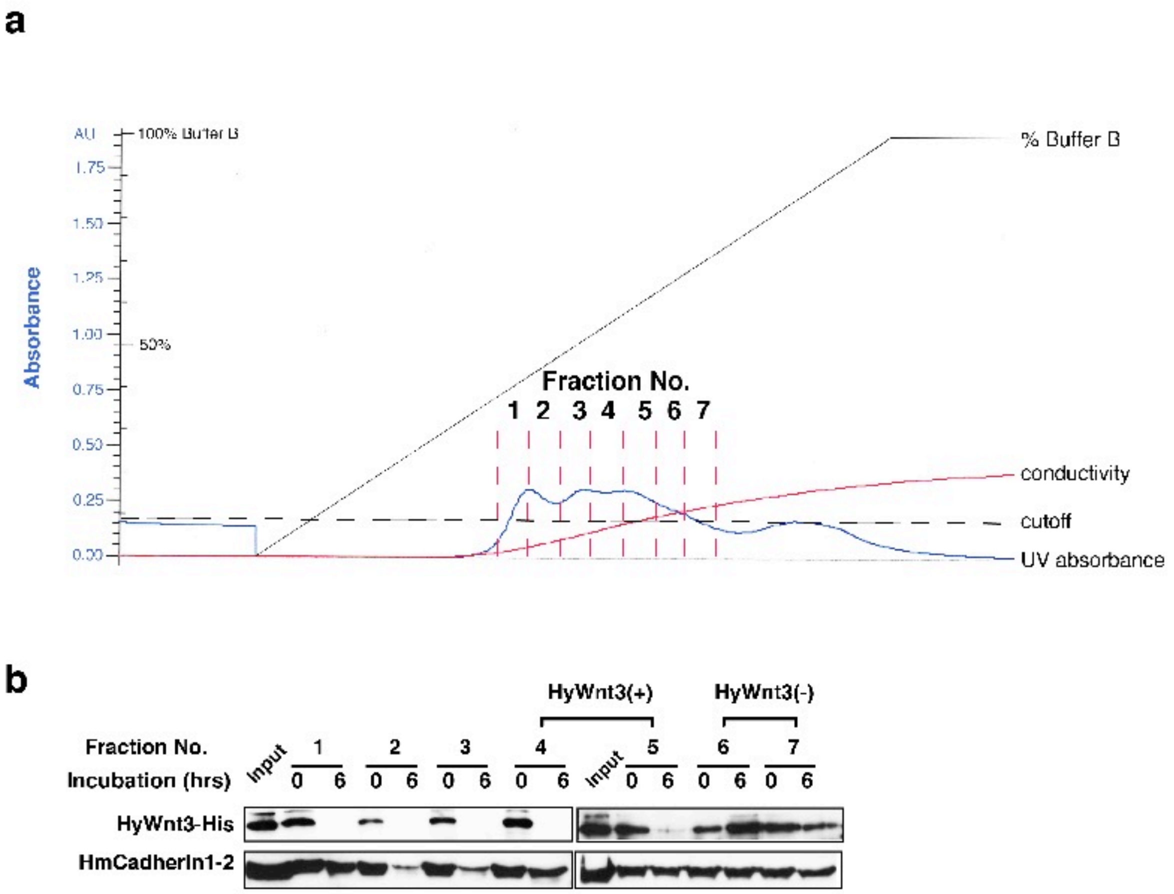


**Fig. S1**. (a) Ion exchange chromatogram of hydra head lysate pool. 7 fractions of 0.5ml exceeding an absorption unit threshold of 0.175 were collected as indicated. The cut-off was chosen to provide a critical total protein concentration (>80μg) for the subsequent proteome analysis. (b) Peak fractions from (a) were re-screened for HyWnt3-His processing activity. A fragment of *Hydra* cadherin extracellular domain comprising the first two N-terminal cadherin repeats (HmCadherin1-2) was used as control substrate to monitor unspecific matrix metalloproteinase activity. Accordingly, fractions 4-5 were pooled and analyzed by mass spectrometry as HyWnt3-His(+) sample, fractions 6-7 as HyWnt3-His(-) sample.

#### Supplemental file 2: Fig. S2

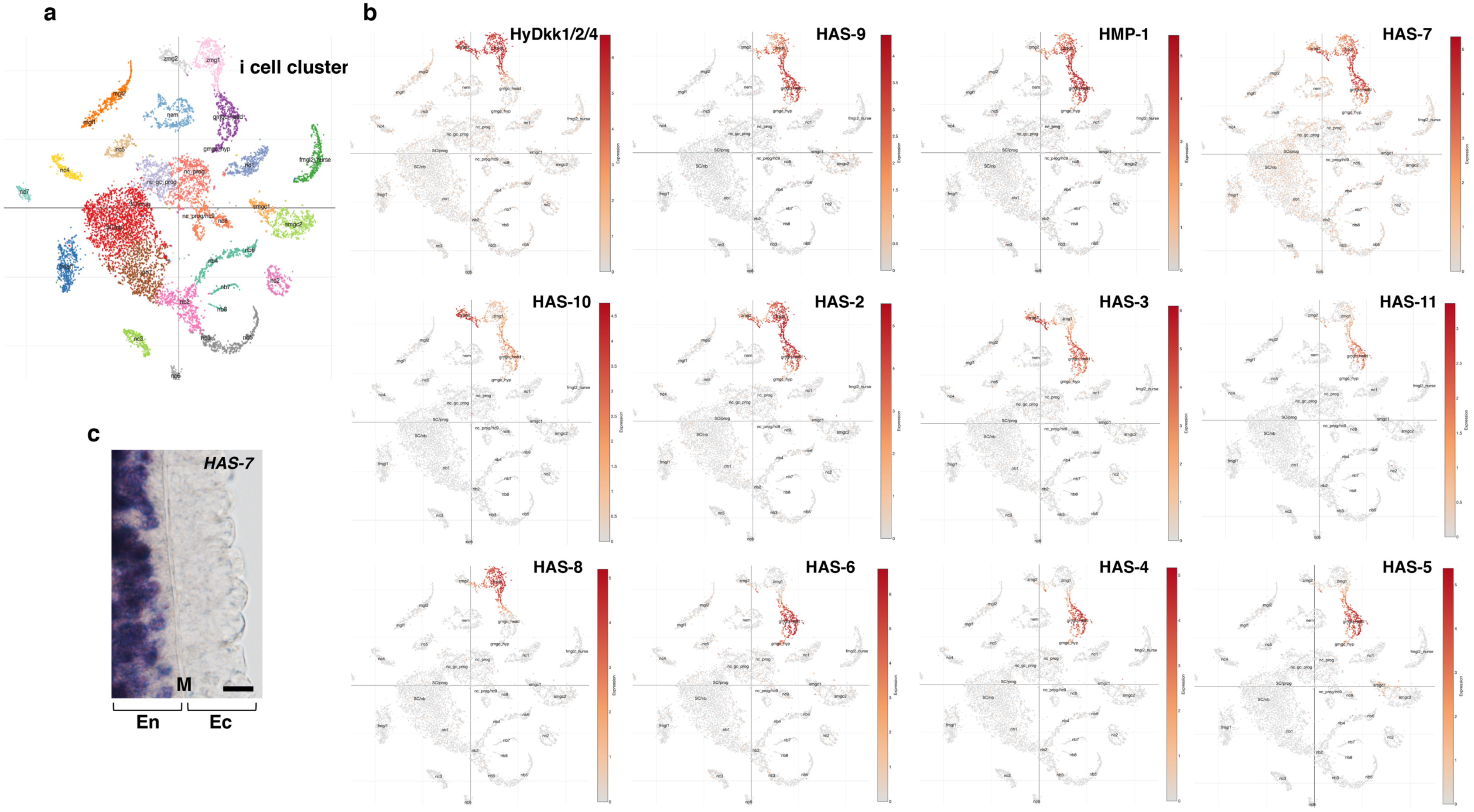

**Fig. S2**. (a) t-SNE representation of interstitial cells with clusters labeled by cell state as presented in [24]. (b) Interstitial cell cluster annotation of *HyDkk1/2/4* and ataxin genes identified in HyWnt3(+) head lysate fraction. The cells in the t-SNE plots were colored based on expression levels for the respective gene using online tools provided at https://singlecell.broadinstitute.org/single_cell/study/SCP260/stem-cell-differentiation-trajectories-in-hydra-resolved-at-single-cell-resolution#study-visualize [24]. The transcript IDs are as follows: *HMP1*: t1098aep, *HAS-1*: t20535aep, *HAS-2*: t18494aep, *HAS-3*: t22149aep, *HAS-4*: t11453aep, *HAS-5*: t596aep, *HAS-6*: t19593aep, *HAS-7*: t16296aep, *HAS-8*: t22154aep, *HAS-9*: t3416aep, *HAS-10*: t10258aep, *HAS-11*: t19316aep. *HyDkk1/2/4*: t8678aep. Cluster label abbreviation key: bat: battery cell, fmgl: female germ-line, gc: gland cell, gmgc: granular mucous gland cell, hyp: hypostome, id: integration doublet, mgl: male germline, nb: nematoblast, nc: neuronal cell, nem: nematocyte, nurse: nurse cells prog: progenitor, SC: stem cell, smgc: spumous mucous gland cell, zmg: zymogen gland cell. Numbers indicate different cell populations within a cluster. (c) Microscopic image showing the epithelial bilayer of the upper gastric region of *Hydra*. *HAS-7* WISH marks gland cells interspersed between the endodermal epithelial cells that are aligned to the central mesoglea (M) separating endo (En)- and ectoderm (Ec). Bar = 20μM.

**Supplemental file 3: Fig. S3**

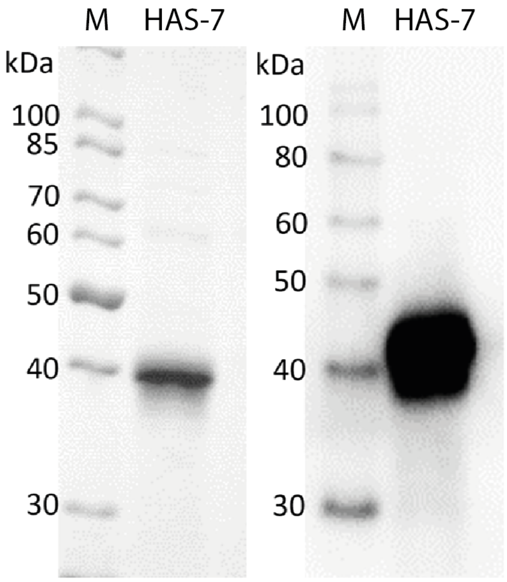


**Fig. S3**. Ni-NTA affinity purified recombinant HAS-7. Separation of by 12% SDS-PAGE was followed by staining with Coomassie brilliant blue (left) or transfer to PVDF and immunodetection (right) using the Penta-His-antibody as described above. For each lane 1.8 *μ*g of recombinant HAS-7 protein eluted with 250 mM imidazole were applied. M, marker proteins as indicated.

#### Supplemental file 4: Table S1

**Table S1a.** Secretome of *Hydra* HL HyWnt3(+) fraction.

| No. | Accession No. | Protein description | Protein Score | Peptide matches | Astacin Protease | Other Protease |
|---|---|---|---|---|---|---|
| 1 | gil828197727 | Fibronectin type III domain-containing protein-like | 1454 | 54 | | |
| 2 | gil828194560 | Contactin-associated protein-like 2 | 967 | 26 | | |
| 3 | gil526117389 | Peroxidase PPOD1-like precursor | 817 | 21 | | |
| 4 | gil449667373 | Peptidyl-prolyl cis-trans isomerase B-like | 793 | 42 | | |
| 5 | gil221114999 | Chitinase-3-like protein 1 | 792 | 20 | | |
| 6 | gil828217704 | Collagen alpha-6(VI) chain-like | 788 | 16 | | |
| 7 | gil828232304 | Blastula protease 10-like / Hydra Astacin 1 (HAS-1) | 668 | 18 | ✂ | |
| 8 | gil15072473 | Peroxidase ppod2 | 575 | 26 | | |
| 9 | gil221113405 | Chymotrypsin-like elastase family member 3B | 573 | 29 | | ✂ |
| 10 | gil449682831 | Chitinase-3-like protein 1 | 509 | 15 | | |
| 11 | gil526117401 | PPOD2 peroxidase-like precursor | 475 | 13 | | |
| 12 | gil828219566 | Chitotriosidase-1-like | 472 | 11 | | |
| 13 | gil828215752 | Zinc metalloproteinase nas-4-like / Hydra Astacin 7 (HAS-7) | 459 | 17 | ✂ | |
| 14 | gil221130733 | Astacin-like metalloprotease toxin 5 / Hydra Astacin 2 (HAS-2) | 415 | 14 | ✂ | |
| 15 | gil828204323 | Hemicentin-2-like isoform X1 | 408 | 12 | | |
| 16 | gil221121571 | Zinc metalloproteinase nas-15-like / Hydra Astacin 9 (HAS-9) | 372 | 14 | ✂ | |
| 17 | gil221129013 | Protein PRY1-like | 341 | 12 | | |
| 18 | gil221119142 | Zinc carboxypeptidase-like | 333 | 11 | | ✂ |
| 19 | gil830260228 | Matrix metalloproteinase-14-like precursor | 318 | 8 | | ✂ |
| 20 | gil828234415 | Carbonic anhydrase 7-like | 313 | 8 | | |
| 21 | gil830260307 | HMP-1 | 311 | 12 | ✂ | |
| 22 | gil449671849 | Protein disulfide-isomerase A3-like | 305 | 7 | | |
| 23 | gil828218801 | Zinc metalloproteinase nas-13-like / Hydra Astacin 11 (HAS-11) | 278 | 11 | ✂ | |
| 24 | gil828208094 | Protein DD3-3-like | 254 | 5 | | |
| 25 | gil828191663 | Neogenin-like | 249 | 5 | | |

| | | | | | | |
|---|---|---|---|---|---|---|
| 26 | gil828194030 | Uncharacterized protein LOC100200589 | 249 | 5 | | |
| 27 | gil828195809 | Astacin-like metalloprotease toxin 5 / Hydra Astacin 3 (HAS-3) | 213 | 5 | ✂ | |
| 28 | gil221126057 | Antistasin-like | 212 | 12 | | |
| 29 | gil828203414 | Protein DD3-3-like | 210 | 5 | | |
| 30 | gil221113429 | Zinc metalloproteinase nas-4-like / Hydra Astacin 8 (HAS-8) | 199 | 7 | ✂ | |
| 31 | gil221111801 | Uncharacterized protein LOC100215485 | 194 | 4 | | |
| 33 | gil526117507 | Kazal-type serine protease inihibitor 2 precursor | 181 | 6 | | |
| 34 | gil449690619 | Low choriolytic enzyme-like / Hydra Astacin 10 (HAS-10) | 173 | 4 | ✂ | |
| 35 | gil828226352 | Glutathione peroxidase-like | 169 | 3 | | |
| 36 | gil221130731 | Protein Span-like / Hydra Astacin 4 (HAS-4) | 161 | 3 | ✂ | |
| 37 | gil449666332 | Zinc metalloproteinase nas-6-like / Hydra Astacin 6 (HAS-6) | 160 | 6 | ✂ | |
| 38 | gil221124062 | Heme-binding protein 1-like | 145 | 5 | | |
| 39 | gil449677685 | Ferritin heavy chain-like | 143 | 6 | | |
| 40 | gil221132488 | Uncharacterized protein LOC100213474 | 140 | 3 | | |
| 41 | gil828215949 | Alpha-L-fucosidase-like isoform X1 | 135 | 3 | | |
| 42 | gil449664802 | Epididymal secretory protein E1-like | 133 | 2 | | |
| 43 | gil221130772 | Carboxypeptidase B-like | 121 | 3 | | ✂ |
| 44 | gil828206799 | Probable G-protein coupled receptor 112 | 113 | 5 | | |
| 45 | gil828224549 | Uncharacterized protein LOC100205745 | 111 | 3 | | |
| 46 | gil449686976 | Protein disulfide-isomerase A6-like | 108 | 3 | | |
| 47 | gil221125481 | Protein Span-like / Hydra Astacin 5 (HAS-5) | 94 | 3 | ✂ | |
| 48 | gil828227729 | Protein DD3-3-like, partial | 86 | 2 | | |
| 49 | gil828225443 | Thrombospondin type-1 domain-containing protein 7A-like | 82 | 2 | | |
| 50 | gil828196752 | Probable G-protein coupled receptor 112 isoform X1 | 81 | 2 | | |
| 51 | gil526117631 | Cysteine-rich BMP regulator 2 precursor | 79 | 2 | | |

Unique protein hits resulting from the Orbitrap mass spectrometry analysis are listed in descending order according to their MASCOT protein score. The table comprises proteins selected for having a leader peptide. The complete list of protein hits for the HL HyWnt3(+) is given in Table S2.

**Table S1b.** Secretome of *Hydra* HL HyWnt3(-) fraction.

| No. | Accession No. | Protein description | Protein Score | Peptide matches | Astacin Protease | Other Protease |
|---|---|---|---|---|---|---|
| 1 | gil828198642 | Uncharacterized protein LOC100198704, partial | 1262 | 32 | | |
| 2 | gil449671849 | Protein disulfide-isomerase A3-like | 1196 | 37 | | |
| 3 | gil828197727 | Fibronectin type III domain-containing protein-like | 777 | 28 | | |
| 4 | gil828201587 | Probable protein disulfide-isomerase A6 | 740 | 23 | | |
| 5 | gil449667373 | Peptidyl-prolyl cis-trans isomerase B-like | 498 | 16 | | |
| 6 | gil221132017 | 78 kda glucose-regulated protein-like | 409 | 10 | | |
| 8 | gil146271914 | Thrombospondin type 1 repeat-containing protein 2 precursor | 402 | 11 | | |
| 9 | gil221113405 | Chymotrypsin-like elastase family member 3B | 395 | 13 | | ✂ |
| 10 | gil449667073 | Acidic mammalian chitinase-like | 390 | 12 | | |
| 11 | gil526117559 | Four-domain proteases inhibitor-like precursor | 386 | 11 | | |
| 12 | gil828217704 | Collagen alpha-6(VI) chain-like | 368 | 7 | | |
| 13 | gil828220687 | Protein disulfide-isomerase A4-like | 365 | 7 | | |
| 14 | gil449686976 | Protein disulfide-isomerase A6-like | 354 | 7 | | |
| 15 | gil828202697 | Peroxiredoxin-4-like | 339 | 10 | | |
| 16 | gil449690552 | Chymotrypsin-like elastase family member 3B | 325 | 11 | | ✂ |
| 17 | gil221114999 | Chitinase-3-like protein 1 | 313 | 8 | | |
| 18 | gil221121571 | Zinc metalloproteinase nas-15-like / Hydra Astacin 9 (HAS-9) | 312 | 7 | ✂ | |
| 19 | gil828199374 | Probable protein disulfide-isomerase A4 | 310 | 7 | | |
| 20 | gil828234415 | Carbonic anhydrase 7-like | 299 | 9 | | |
| 21 | gil221132488 | Uncharacterized protein LOC100213474 | 297 | 8 | | |
| 22 | gil828194030 | Uncharacterized protein LOC100200589 | 294 | 7 | | |
| 23 | gil221118599 | Dolichyl-diphosphooligosaccharide--protein glycosyltransferase subunit 1-like | 293 | 6 | | |
| 24 | gil828225443 | Thrombospondin type-1 domain-containing protein 7A-like | 291 | 7 | | |
| 25 | gil449685905 | Acid ceramidase-like | 277 | 8 | | |
| 26 | gil221119142 | Zinc carboxypeptidase-like | 275 | 8 | | ✂ |
| 27 | gil828218618 | Putative phospholipase B-like 2 | 256 | 4 | | |

| | | | | | | |
|---|---|---|---|---|---|---|
| 28 | gil221129013 | Protein PRY1-like | 249 | 5 | | |
| 29 | gil828224104 | Beta-glucuronidase-like | 218 | 6 | | |
| 30 | gil449666857 | Lysosomal aspartic protease-like | 212 | 6 | | ✂ |
| 31 | gil828215949 | Alpha-L-fucosidase-like isoform X1 | 201 | 5 | | |
| 32 | gil828213548 | Uncharacterized protein LOC105845774 | 193 | 4 | | |
| 33 | gil828197619 | Lysosomal alpha-mannosidase-like | 175 | 4 | | |
| 34 | gil449684402 | Endochitinase 1-like isoform X1 | 162 | 3 | | |
| 35 | gil449678353 | Bandaporin-like (pore forming toxin) | 161 | 4 | | |
| 36 | gil221126057 | Antistasin-like | 154 | 11 | | |
| 37 | gil828191663 | Neogenin-like (receptor) | 145 | 4 | | |
| 38 | gil221090861 | Cathepsin L1-like | 141 | 3 | | ✂ |
| 39 | gil449666332 | Zinc metalloproteinase nas-6-like / Hydra Astacin 6 (HAS-6) | 136 | 4 | ✂ | |
| 40 | gil526117401 | PPOD2 peroxidase-like precursor | 135 | 2 | | |
| 41 | gil449667021 | Zinc metalloproteinase nas-14-like | 135 | 3 | ✂ | |
| 42 | gil221124062 | Heme-binding protein 1-like | 134 | 5 | | |
| 43 | gil526117507 | Kazal-type serine protease inihibitor 2 precursor | 134 | 5 | | |
| 44 | gil828232304 | Blastula protease 10-like / Hydra Astacin 1 (HAS-1) | 129 | 3 | ✂ | |
| 45 | gil15072473 | Peroxidase ppod2 | 119 | 3 | | |
| 46 | gil449670322 | Dipeptidyl peptidase 1-like | 112 | 3 | | ✂ |
| 47 | gil221130733 | Astacin-like metalloprotease toxin 5 (2) | 108 | 2 | ✂ | |
| 48 | gil449670247 | Probable inactive purple acid phosphatase 2 | 104 | 2 | | |
| 49 | gil449665331 | Contactin-associated protein-like 5 | 102 | 3 | | |
| 50 | gil526117489 | Kazal-type serine protease inihibitor 3 precursor | 102 | 5 | | |
| 51 | gil828196768 | Calsequestrin-2-like | 101 | 3 | | |
| 52 | gil449680876 | Endochitinase 4-like | 98 | 2 | | |
| 53 | gil221121832 | Multiple inositol polyphosphate phosphatase 1-like | 96 | 2 | | |
| 54 | gil449687197 | Golgi-associated plant pathogenesis-related protein 1-like | 92 | 3 | | |
| 55 | gil828204323 | Hemicentin-2-like isoform X1 | 92 | 2 | | |
| 56 | gil449682262 | Zinc metalloproteinase nas-13-like / Hydra Astacin 11 (HAS-11) | 87 | 3 | ✂ | |
| 57 | gil221113277 | Uncharacterized protein LOC100214198 | 86 | 4 | | |

| | | | | |
|---|---|---|---|---|
| 58 | gil828212124 | MAM and LDL-receptor class A domain-containing protein 1-like | 79 | 2 |
| 59 | gil449679397 | Uncharacterized protein LOC100197967 | 75 | 3 |
| 60 | gil221124690 | Endoplasmin-like | 71 | 2 |
| 61 | gil828195809 | Astacin-like metalloprotease toxin 5 / Hydra Astacin 3 (HAS-3) | 67 | 2 |
| 62 | gil828231348 | Transmembrane 9 superfamily member 2-like | 67 | 2 |

Unique protein hits resulting from the Orbitrap mass spectrometry analysis are listed in descending order according to their MASCOT protein score. The table comprises proteins selected for having a leader peptide. The complete list of protein hits for the HL HyWnt3(-) is given in Table S2.

### Supplemental file 5: Table S2

**Table S2.** Complete proteome data of HyWnt3(+) and HyWnt3(-) HL fractions.

HyWnt3(+) Fraction

| Hit No. | Accession No. | Protein Description | Score | Mass [Da] | Prot Matches | sign Prot Matches | Prot Sequences | sign Prot Sequences | Coverage [%] |
|---|---|---|---|---|---|---|---|---|---|
| 1 | gi\|47132620 | keratin, type II cytoskeletal 2 epidermal [Homo sapiens] | 2873 | 65678 | 117 | 39 | 41 | 21 | 75,6 |
| 2 | gi\|11935049 | keratin 1 [Homo sapiens] | 2763 | 66198 | 174 | 51 | 45 | 19 | 64,1 |
| 3 | gi\|21961605 | Keratin 10 [Homo sapiens] | 2268 | 59020 | 116 | 32 | 32 | 15 | 51,9 |
| 4 | gi\|795247621 | PREDICTED: keratin, type I cytoskeletal 16 isoform X1 [Mandrillus leucophaeu | 2232 | 92585 | 88 | 17 | 41 | 11 | 49,1 |
| 5 | gi\|332206121 | PREDICTED: keratin, type II cytoskeletal 2 epidermal isoform X1 [Nomascus leu | 2108 | 66822 | 92 | 30 | 31 | 15 | 42,2 |
| 6 | gi\|795536398 | PREDICTED: keratin, type II cytoskeletal 1 [Cercocebus atys] | 2040 | 65879 | 141 | 42 | 35 | 15 | 47,2 |
| 7 | gi\|435476 | cytokeratin 9 [Homo sapiens] | 1984 | 62320 | 95 | 22 | 31 | 12 | 60,4 |
| 8 | gi\|908801 | keratin type II [Homo sapiens] | 1907 | 60448 | 74 | 17 | 35 | 10 | 53,2 |
| 9 | gi\|5031839 | keratin, type II cytoskeletal 6C [Homo sapiens] | 1905 | 60293 | 75 | 17 | 35 | 10 | 53,2 |
| 10 | gi\|1195531 | type I keratin 16 [Homo sapiens] | 1882 | 51548 | 76 | 15 | 33 | 9 | 68,5 |
| 11 | gi\|155969697 | keratin, type II cytoskeletal 6C [Homo sapiens] | 1877 | 60273 | 74 | 17 | 35 | 10 | 53,2 |
| 12 | gi\|635065484 | PREDICTED: keratin, type II cytoskeletal 6A [Chlorocebus sabaeus] | 1848 | 60236 | 74 | 16 | 34 | 9 | 48,4 |
| 13 | gi\|795360291 | PREDICTED: keratin, type II cytoskeletal 6A [Colobus angolensis palliatus] | 1812 | 60225 | 79 | 20 | 33 | 10 | 46,8 |
| 14 | gi\|908803 | keratin type II [Homo sapiens] | 1805 | 60472 | 72 | 16 | 34 | 9 | 53,2 |
| 15 | gi\|395744310 | PREDICTED: keratin, type II cytoskeletal 6B [Pongo abelii] | 1685 | 60398 | 72 | 16 | 31 | 9 | 45,7 |
| 16 | gi\|18999435 | Keratin 5 [Homo sapiens] | 1572 | 62568 | 62 | 16 | 31 | 10 | 41,7 |
| 17 | gi\|828197727 | PREDICTED: fibronectin type III domain-containing protein-like [Hydra vulgari | 1454 | 152931 | 54 | 9 | 26 | 9 | 25,3 |
| 18 | gi\|803121007 | PREDICTED: keratin, type I cytoskeletal 14 isoform X1 [Ovis aries] | 1287 | 49277 | 54 | 9 | 23 | 6 | 48,5 |
| 19 | gi\|676264984 | Keratin, type II cytoskeletal 1 [Fukomys damarensis] | 1202 | 159046 | 63 | 21 | 20 | 8 | 11,4 |
| 20 | gi\|821004446 | PREDICTED: keratin, type II cytoskeletal 1 [Nomascus leucogenys] | 1149 | 32860 | 81 | 26 | 20 | 7 | 48,8 |
| 21 | gi\|927195534 | PREDICTED: keratin, type I cytoskeletal 17 isoform X2 [Sus scrofa] | 1146 | 50320 | 43 | 9 | 20 | 6 | 38,4 |
| 22 | gi\|4557701 | keratin, type I cytoskeletal 17 [Homo sapiens] | 1110 | 48361 | 39 | 7 | 20 | 6 | 41,2 |
| 23 | gi\|828194560 | PREDICTED: contactin-associated protein-like 2 [Hydra vulgaris] | 967 | 138908 | 26 | 4 | 20 | 4 | 21,7 |
| 24 | gi\|558156247 | PREDICTED: keratin, type II cytoskeletal 1 [Myotis lucifugus] | 933 | 63621 | 60 | 20 | 14 | 7 | 17,6 |
| 25 | gi\|221108525 | PREDICTED: malate dehydrogenase, mitochondrial-like [Hydra vulgaris] | 866 | 36471 | 23 | 7 | 16 | 4 | 52,9 |
| 26 | gi\|1351907 | RecName: Full=Serum albumin; AltName: Full=BSA; AltName: Allergen=Bos d 6 | 828 | 71244 | 29 | 7 | 17 | 4 | 32,9 |
| 27 | gi\|526117389 | peroxidase PPOD1-like precursor [Hydra vulgaris] | 817 | 32270 | 21 | 5 | 15 | 4 | 58,6 |
| 28 | gi\|301775745 | PREDICTED: keratin, type II cytoskeletal 2 epidermal isoform X1 [Ailuropoda m | 801 | 65009 | 35 | 13 | 12 | 6 | 16 |
| 29 | gi\|449667373 | PREDICTED: peptidyl-prolyl cis-trans isomerase B-like [Hydra vulgaris] | 793 | 22428 | 42 | 13 | 12 | 5 | 69,2 |
| 30 | gi\|221114999 | PREDICTED: chitinase-3-like protein 1 [Hydra vulgaris] | 792 | 52131 | 20 | 3 | 17 | 3 | 42,6 |
| 31 | gi\|344257799 | Keratin, type II cytoskeletal 1b [Cricetulus griseus] | 790 | 93670 | 53 | 12 | 14 | 5 | 12,8 |
| 32 | gi\|828217704 | PREDICTED: collagen alpha-6(VI) chain-like [Hydra vulgaris] | 788 | 68564 | 16 | 6 | 13 | 5 | 26,7 |
| 33 | gi\|449690552 | PREDICTED: chymotrypsin-like elastase family member 3B [Hydra vulgaris] | 764 | 29089 | 43 | 6 | 14 | 5 | 66,5 |
| 34 | gi\|828208682 | PREDICTED: filamin-A-like [Hydra vulgaris] | 698 | 294642 | 15 | 2 | 15 | 2 | 6,8 |
| 35 | gi\|828232304 | PREDICTED: blastula protease 10-like [Hydra vulgaris] | 668 | 27019 | 18 | 5 | 10 | 4 | 58 |
| 36 | gi\|828203690 | PREDICTED: hemicentin-1-like [Hydra vulgaris] | 667 | 85973 | 15 | 4 | 13 | 4 | 19,6 |
| 37 | gi\|221122379 | PREDICTED: staphylococcal nuclease domain-containing protein 1-like [Hydra | 647 | 100759 | 19 | 4 | 11 | 4 | 15,6 |
| 38 | gi\|828229911 | PREDICTED: 2-amino-3-ketobutyrate coenzyme A ligase, mitochondrial-like [H | 645 | 46963 | 17 | 4 | 12 | 3 | 33,8 |
| 39 | gi\|221102389 | PREDICTED: protein PRY1-like [Hydra vulgaris] | 599 | 20363 | 48 | 5 | 12 | 3 | 74,9 |
| 40 | gi\|828224124 | PREDICTED: catalase-like [Hydra vulgaris] | 578 | 57689 | 18 | 1 | 14 | 1 | 33,3 |
| 41 | gi\|15072473 | peroxidase ppod2 [Hydra vulgaris] | 575 | 33192 | 26 | 1 | 12 | 1 | 38,3 |
| 42 | gi\|221113405 | PREDICTED: chymotrypsin-like elastase family member 3B [Hydra vulgaris] | 573 | 29175 | 29 | 4 | 10 | 3 | 52,1 |
| 43 | gi\|829991204 | PREDICTED: desmoplakin isoform X1 [Microcebus murinus] | 565 | 335112 | 14 | 2 | 14 | 2 | 5,7 |
| 44 | gi\|828218611 | PREDICTED: nucleoside diphosphate kinase A-like [Hydra vulgaris] | 520 | 25885 | 22 | 2 | 11 | 2 | 53,2 |
| 45 | gi\|449682831 | PREDICTED: chitinase-3-like protein 1 [Hydra vulgaris] | 509 | 52203 | 15 | 1 | 11 | 1 | 29,8 |

| | | | | | | | | | |
|---|---|---|---|---|---|---|---|---|---|
| 46 | gi\|198139671 | fructose 1,6-bisphosphate aldolase [Artemia franciscana] | 506 | 25865 | 12 | 7 | 8 | 6 | 43,5 |
| 47 | gi\|697450140 | Keratin, type II cytoskeletal 75 [Charadrius vociferus] | 502 | 63147 | 35 | 12 | 9 | 2 | 12,5 |
| 48 | gi\|526117401 | PPOD2 peroxidase-like precursor [Hydra vulgaris] | 475 | 32661 | 13 | 1 | 11 | 1 | 38,4 |
| 49 | gi\|449669348 | PREDICTED: gamma-glutamyltranspeptidase 1-like [Hydra vulgaris] | 475 | 63187 | 17 | 1 | 12 | 1 | 23,6 |
| 50 | gi\|828219566 | PREDICTED: chitotriosidase-1-like [Hydra vulgaris] | 472 | 27976 | 11 | 2 | 9 | 2 | 38,9 |
| 51 | gi\|221115109 | PREDICTED: profilin-like [Hydra vulgaris] | 467 | 14833 | 20 | 6 | 8 | 3 | 75,5 |
| 52 | gi\|221130743 | PREDICTED: uncharacterized protein LOC100211495 [Hydra vulgaris] | 461 | 13958 | 24 | 6 | 7 | 3 | 52,4 |
| 53 | gi\|828215752 | PREDICTED: zinc metalloproteinase nas-4-like [Hydra vulgaris] | 459 | 38857 | 17 | 1 | 10 | 1 | 33,1 |
| 54 | gi\|260223405 | hypothetical protein Csp_B21540 [Curvibacter putative symbiont of Hydra ma | 435 | 35777 | 7 | 5 | 7 | 5 | 33,2 |
| 55 | gi\|31074631 | keratin 1b [Homo sapiens] | 430 | 62049 | 27 | 8 | 8 | 2 | 12,6 |
| 56 | gi\|449662449 | PREDICTED: polyubiquitin-B [Hydra vulgaris] | 424 | 42564 | 13 | 2 | 8 | 2 | 20,8 |
| 57 | gi\|221130733 | PREDICTED: astacin-like metalloprotease toxin 5 [Hydra vulgaris] | 415 | 27816 | 14 | 4 | 6 | 3 | 33,6 |
| 58 | gi\|828204323 | PREDICTED: hemicentin-2-like isoform X1 [Hydra vulgaris] | 408 | 158275 | 12 | 1 | 9 | 1 | 7,9 |
| 59 | gi\|3318722 | Chain E, Leech-Derived Tryptase InhibitorTRYPSIN COMPLEX | 399 | 24142 | 37 | 3 | 6 | 1 | 26 |
| 60 | gi\|194373749 | unnamed protein product [Homo sapiens] | 390 | 62862 | 14 | 1 | 8 | 1 | 14,2 |
| 61 | gi\|221121571 | PREDICTED: zinc metalloproteinase nas-15-like [Hydra vulgaris] | 372 | 40888 | 14 | 0 | 9 | 0 | 28,5 |
| 62 | gi\|829966121 | PREDICTED: LOW QUALITY PROTEIN: polyubiquitin-B [Microcebus murinus] | 359 | 25839 | 10 | 0 | 8 | 0 | 33,6 |
| 63 | gi\|646697674 | Fructose-bisphosphate aldolase [Zootermopsis nevadensis] | 343 | 39761 | 8 | 5 | 6 | 4 | 15,4 |
| 64 | gi\|221129013 | PREDICTED: protein PRY1-like [Hydra vulgaris] | 341 | 18775 | 12 | 2 | 7 | 2 | 56,2 |
| 65 | gi\|565324183 | nucleoside diphosphate kinase-like [Hydra vulgaris] | 340 | 17048 | 12 | 1 | 7 | 1 | 58,9 |
| 66 | gi\|449667021 | PREDICTED: zinc metalloproteinase nas-14-like [Hydra vulgaris] | 335 | 20331 | 12 | 1 | 7 | 1 | 42,3 |
| 67 | gi\|221119142 | PREDICTED: zinc carboxypeptidase-like [Hydra vulgaris] | 333 | 47718 | 11 | 1 | 9 | 1 | 29 |
| 68 | gi\|830260228 | matrix metalloproteinase-14-like precursor [Hydra vulgaris] | 318 | 55390 | 8 | 2 | 8 | 2 | 16,1 |
| 69 | gi\|449661907 | PREDICTED: low choriolytic enzyme-like [Hydra vulgaris] | 317 | 33953 | 12 | 2 | 6 | 1 | 22,3 |
| 70 | gi\|828195674 | PREDICTED: gamma-glutamyltranspeptidase 1-like [Hydra vulgaris] | 316 | 67827 | 7 | 1 | 7 | 1 | 14,3 |
| 71 | gi\|828234415 | PREDICTED: carbonic anhydrase 7-like [Hydra vulgaris] | 313 | 36918 | 8 | 0 | 8 | 0 | 29,1 |
| 72 | gi\|830260307 | zinc metalloproteinase nas-15-like precursor [Hydra vulgaris] | 311 | 33203 | 12 | 1 | 6 | 1 | 30,5 |
| 73 | gi\|449671849 | PREDICTED: protein disulfide-isomerase A3-like [Hydra vulgaris] | 305 | 55425 | 7 | 0 | 7 | 0 | 15,7 |
| 74 | gi\|242025190 | Fructose-bisphosphate aldolase, putative [Pediculus humanus corporis] | 298 | 40383 | 6 | 4 | 5 | 3 | 11,5 |
| 75 | gi\|725586596 | PREDICTED: polyubiquitin-like [Saimiri boliviensis boliviensis] | 297 | 18868 | 8 | 0 | 6 | 0 | 41,2 |
| 76 | gi\|828209655 | PREDICTED: thymocyte nuclear protein 1-like [Hydra vulgaris] | 278 | 29264 | 9 | 0 | 8 | 0 | 30,8 |
| 77 | gi\|449682262 | PREDICTED: zinc metalloproteinase nas-13-like [Hydra vulgaris] | 278 | 58069 | 11 | 1 | 7 | 1 | 18,3 |
| 78 | gi\|159163108 | Chain A, Solution Structure Of The Designed Hydrophobic Core Mutant Of Ubi | 277 | 8580 | 8 | 0 | 6 | 0 | 64,5 |
| 79 | gi\|449678439 | PREDICTED: peptidyl-prolyl cis-trans isomerase 5-like [Hydra vulgaris] | 266 | 25229 | 9 | 1 | 6 | 1 | 34,1 |
| 80 | gi\|499801699 | membrane protein [Polaromonas sp. JS666] | 259 | 41227 | 4 | 3 | 4 | 3 | 15,8 |
| 81 | gi\|221131289 | PREDICTED: methylmalonyl-CoA epimerase, mitochondrial-like [Hydra vulgari | 259 | 18258 | 9 | 1 | 5 | 1 | 33,9 |
| 82 | gi\|828224553 | PREDICTED: delta-1-pyrroline-5-carboxylate dehydrogenase, mitochondrial-li | 258 | 28510 | 6 | 1 | 6 | 1 | 29,2 |
| 83 | gi\|697423976 | Keratin, type II cytoskeletal 4, partial [Tinamus guttatus] | 256 | 59243 | 12 | 0 | 5 | 0 | 7 |
| 84 | gi\|828208094 | PREDICTED: protein DD3-3-like [Hydra vulgaris] | 254 | 69972 | 5 | 1 | 5 | 1 | 10,7 |
| 85 | gi\|33304714 | ubiquitin/actin fusion protein 2 [Bigelowiella natans] | 253 | 51325 | 8 | 0 | 6 | 0 | 14,4 |
| 86 | gi\|828191663 | PREDICTED: neogenin-like [Hydra vulgaris] | 249 | 209395 | 5 | 1 | 5 | 1 | 3,5 |
| 87 | gi\|828194030 | PREDICTED: uncharacterized protein LOC100200589 [Hydra vulgaris] | 249 | 86226 | 5 | 0 | 5 | 0 | 8,2 |
| 88 | gi\|676246284 | Keratin, type I cytoskeletal 15 [Balearica regulorum gibbericeps] | 248 | 48549 | 11 | 0 | 6 | 0 | 10,8 |
| 89 | gi\|828214768 | PREDICTED: delta-1-pyrroline-5-carboxylate dehydrogenase, mitochondrial-li | 247 | 40600 | 7 | 1 | 6 | 1 | 20,3 |
| 90 | gi\|449675219 | PREDICTED: uncharacterized protein LOC101234497 [Hydra vulgaris] | 240 | 27697 | 6 | 1 | 6 | 1 | 28,9 |
| 91 | gi\|33346945 | ubiquitin/actin fusion protein [Gymnochlora stellata] | 236 | 49580 | 7 | 0 | 5 | 0 | 13,3 |
| 92 | gi\|30506 | desmoglein type 1 [Homo sapiens] | 230 | 114670 | 7 | 0 | 6 | 0 | 7,3 |
| 93 | gi\|16751921 | dermcidin isoform 1 preproprotein [Homo sapiens] | 229 | 11391 | 8 | 1 | 4 | 1 | 35,5 |

| | | | | | | | | | |
|---|---|---|---|---|---|---|---|---|---|
| 94 | gi\|762885 | Plakoglobin [Homo sapiens] | 225 | 82381 | 5 | 0 | 5 | 0 | 8,7 |
| 95 | gi\|828195809 | PREDICTED: astacin-like metalloprotease toxin 5 [Hydra vulgaris] | 213 | 28949 | 5 | 0 | 5 | 0 | 19,8 |
| 96 | gi\|221126057 | PREDICTED: antistasin-like [Hydra vulgaris] | 212 | 26072 | 12 | 1 | 5 | 1 | 27,4 |
| 97 | gi\|828217561 | PREDICTED: zinc metalloproteinase nas-15-like, partial [Hydra vulgaris] | 211 | 31952 | 4 | 1 | 4 | 1 | 16,5 |
| 98 | gi\|828203414 | PREDICTED: protein DD3-3-like [Hydra vulgaris] | 210 | 76301 | 5 | 0 | 5 | 0 | 13 |
| 99 | gi\|28336 | mutant beta-actin (beta'-actin) [Homo sapiens] | 208 | 42128 | 6 | 0 | 6 | 0 | 18,4 |
| 100 | gi\|830260001 | tight junction protein ZO-2-like [Hydra vulgaris] | 204 | 191163 | 3 | 2 | 3 | 2 | 2,8 |
| 101 | gi\|221113429 | PREDICTED: zinc metalloproteinase nas-4-like [Hydra vulgaris] | 199 | 41939 | 7 | 0 | 5 | 0 | 14,2 |
| 102 | gi\|221111801 | PREDICTED: uncharacterized protein LOC100215485 [Hydra vulgaris] | 194 | 19588 | 4 | 1 | 3 | 1 | 26,7 |
| 103 | gi\|828221194 | PREDICTED: fibronectin type III domain-containing protein-like [Hydra vulgari | 193 | 143336 | 5 | 0 | 5 | 0 | 5,2 |
| 104 | gi\|828198152 | PREDICTED: glycogenin-1-like [Hydra vulgaris] | 192 | 39767 | 4 | 1 | 4 | 1 | 14,1 |
| 105 | gi\|740383561 | membrane protein [Xenophilus azovorans] | 191 | 41199 | 3 | 2 | 3 | 2 | 12,1 |
| 106 | gi\|586632450 | Purine-binding protein precursor [Hydrogenophaga sp. T4] | 190 | 31703 | 3 | 2 | 3 | 2 | 15,5 |
| 107 | gi\|84402 | glutathione transferase (EC 2.5.1.18) - fluke (Schistosoma japonicum) (fragmer | 190 | 25834 | 5 | 1 | 4 | 1 | 20,1 |
| 108 | gi\|449689369 | PREDICTED: protein NipSnap homolog 1-like [Hydra vulgaris] | 188 | 32344 | 5 | 0 | 5 | 0 | 21,5 |
| 109 | gi\|21755908 | unnamed protein product [Homo sapiens] | 183 | 57583 | 4 | 0 | 4 | 0 | 7,9 |
| 110 | gi\|526117507 | kazal-type serine protease inihibitor 2 precursor [Hydra vulgaris] | 181 | 19400 | 6 | 0 | 6 | 0 | 33,9 |
| 111 | gi\|828203917 | PREDICTED: aminomethyltransferase, mitochondrial-like [Hydra vulgaris] | 178 | 44995 | 4 | 0 | 4 | 0 | 11 |
| 112 | gi\|449690619 | PREDICTED: low choriolytic enzyme-like [Hydra vulgaris] | 173 | 38696 | 4 | 0 | 4 | 0 | 15,2 |
| 113 | gi\|828226352 | PREDICTED: glutathione peroxidase-like [Hydra vulgaris] | 169 | 26056 | 3 | 1 | 3 | 1 | 20,9 |
| 114 | gi\|828222174 | PREDICTED: 3-hydroxyacyl-CoA dehydrogenase type-2-like [Hydra vulgaris] | 168 | 26807 | 3 | 1 | 3 | 1 | 13,4 |
| 115 | gi\|828206951 | PREDICTED: uncharacterized protein LOC100208668 isoform X1 [Hydra vulga | 166 | 253566 | 5 | 0 | 5 | 0 | 2,9 |
| 116 | gi\|221130731 | PREDICTED: protein SpAN-like [Hydra vulgaris] | 161 | 35171 | 3 | 1 | 3 | 1 | 10,3 |
| 117 | gi\|449666332 | PREDICTED: zinc metalloproteinase nas-6-like [Hydra vulgaris] | 160 | 33291 | 6 | 0 | 4 | 0 | 19,1 |
| 118 | gi\|221124062 | PREDICTED: heme-binding protein 1-like [Hydra vulgaris] | 145 | 29921 | 5 | 0 | 4 | 0 | 14,8 |
| 119 | gi\|449677685 | PREDICTED: ferritin heavy chain-like [Hydra vulgaris] | 143 | 27387 | 6 | 0 | 5 | 0 | 25,9 |
| 120 | gi\|221132488 | PREDICTED: uncharacterized protein LOC100213474 [Hydra vulgaris] | 140 | 27971 | 3 | 1 | 3 | 1 | 14,7 |
| 121 | gi\|449678564 | PREDICTED: lysosome-associated membrane glycoprotein 1-like [Hydra vulgari | 137 | 21879 | 5 | 0 | 3 | 0 | 15,7 |
| 122 | gi\|828215949 | PREDICTED: alpha-L-fucosidase-like isoform X1 [Hydra vulgaris] | 135 | 56266 | 3 | 0 | 3 | 0 | 5,4 |
| 123 | gi\|449664802 | PREDICTED: epididymal secretory protein E1-like [Hydra vulgaris] | 133 | 16839 | 2 | 1 | 2 | 1 | 18,7 |
| 124 | gi\|828216609 | PREDICTED: salivary glue protein Sgs-3-like, partial [Hydra vulgaris] | 129 | 34483 | 2 | 1 | 2 | 1 | 11,2 |
| 125 | gi\|221132768 | PREDICTED: RNA polymerase II transcriptional coactivator-like [Hydra vulgaris | 129 | 13128 | 3 | 0 | 3 | 0 | 27 |
| 126 | gi\|395826450 | PREDICTED: keratin, type I cytoskeletal 28 [Otolemur garnettii] | 129 | 51136 | 13 | 0 | 3 | 0 | 5,4 |
| 127 | gi\|449683097 | PREDICTED: peptidyl-prolyl cis-trans isomerase-like [Hydra vulgaris] | 125 | 17530 | 3 | 0 | 3 | 0 | 24,2 |
| 128 | gi\|828222429 | PREDICTED: AP-2 complex subunit alpha-2-like [Hydra vulgaris] | 122 | 103472 | 4 | 0 | 4 | 0 | 4,2 |
| 129 | gi\|526117746 | flp protein [Hydra vulgaris] | 121 | 14995 | 4 | 0 | 3 | 0 | 26,2 |
| 130 | gi\|221130772 | PREDICTED: carboxypeptidase B-like [Hydra vulgaris] | 121 | 51584 | 3 | 0 | 3 | 0 | 5,8 |
| 131 | gi\|46621276 | CEP152 protein, partial [Homo sapiens] | 118 | 68324 | 2 | 1 | 2 | 1 | 7,4 |
| 132 | gi\|58005 | aprotinin [synthetic construct] | 116 | 7011 | 4 | 1 | 2 | 1 | 47,5 |
| 133 | gi\|62122917 | filaggrin-2 [Homo sapiens] | 114 | 249296 | 3 | 1 | 2 | 1 | 1 |
| 134 | gi\|828206799 | PREDICTED: probable G-protein coupled receptor 112 [Hydra vulgaris] | 113 | 98355 | 5 | 0 | 3 | 0 | 3,8 |
| 135 | gi\|828224549 | PREDICTED: uncharacterized protein LOC100205745 [Hydra vulgaris] | 111 | 70074 | 3 | 0 | 3 | 0 | 5,5 |
| 136 | gi\|449686976 | PREDICTED: protein disulfide-isomerase A6-like [Hydra vulgaris] | 108 | 48142 | 3 | 0 | 2 | 0 | 8,2 |
| 137 | gi\|221121838 | PREDICTED: endothelin-converting enzyme 1-like [Hydra vulgaris] | 105 | 88293 | 3 | 0 | 2 | 0 | 2,7 |
| 138 | gi\|828209711 | PREDICTED: protein DD3-3-like [Hydra vulgaris] | 105 | 62180 | 2 | 0 | 2 | 0 | 3,6 |
| 139 | gi\|828235608 | PREDICTED: cytosol aminopeptidase-like [Hydra vulgaris] | 104 | 56366 | 3 | 0 | 3 | 0 | 7,5 |
| 140 | gi\|828197350 | PREDICTED: contactin-2-like [Hydra vulgaris] | 100 | 93069 | 2 | 1 | 2 | 1 | 2,2 |
| 141 | gi\|828202743 | PREDICTED: uncharacterized protein LOC100202739 isoform X1 [Hydra vulga | 98 | 869663 | 3 | 0 | 3 | 0 | 0,4 |

| | | | | | | | | | |
|---|---|---|---|---|---|---|---|---|---|
| 142 | gi\|223130 | fibrinogen betaB 1-118 | 97 | 12891 | 5 | 2 | 1 | 1 | 11,9 |
| 143 | gi\|449690428 | PREDICTED: nascent polypeptide-associated complex subunit alpha-like [Hydr | 96 | 24795 | 2 | 0 | 2 | 0 | 11,8 |
| 144 | gi\|449687420 | PREDICTED: uncharacterized protein LOC100205832 isoform X1 [Hydra vulga | 96 | 24618 | 2 | 0 | 2 | 0 | 8,4 |
| 145 | gi\|449674503 | PREDICTED: uncharacterized protein LOC101239382 [Hydra vulgaris] | 95 | 18169 | 2 | 1 | 2 | 1 | 17,4 |
| 146 | gi\|221125481 | PREDICTED: protein SpAN-like [Hydra vulgaris] | 94 | 33407 | 3 | 0 | 2 | 0 | 10 |
| 147 | gi\|28557150 | hornerin [Homo sapiens] | 92 | 48797 | 3 | 0 | 2 | 0 | 8,1 |
| 148 | gi\|828218801 | PREDICTED: zinc metalloproteinase nas-13-like [Hydra vulgaris] | 91 | 57813 | 3 | 0 | 2 | 0 | 4,8 |
| 149 | gi\|449680259 | PREDICTED: glutathione peroxidase 2-like [Hydra vulgaris] | 89 | 23459 | 5 | 0 | 3 | 0 | 15,3 |
| 150 | gi\|170582740 | cyclophilin-type peptidyl-prolyl cis-trans isomerase-15, Bmcyp-5 [Brugia mala | 88 | 22456 | 2 | 0 | 2 | 0 | 10,8 |
| 151 | gi\|493257619 | azurin [Achromobacter piechaudii] | 87 | 16082 | 1 | 1 | 1 | 1 | 10,7 |
| 152 | gi\|828227729 | PREDICTED: protein DD3-3-like, partial [Hydra vulgaris] | 86 | 54925 | 2 | 0 | 2 | 0 | 4 |
| 153 | gi\|449668124 | PREDICTED: CUGBP Elav-like family member 2 isoform X3 [Hydra vulgaris] | 86 | 55769 | 1 | 1 | 1 | 1 | 3,1 |
| 154 | gi\|828200742 | PREDICTED: ADP-ribosyl cyclase/cyclic ADP-ribose hydrolase-like [Hydra vulga | 86 | 35219 | 1 | 1 | 1 | 1 | 3,6 |
| 155 | gi\|828225443 | PREDICTED: thrombospondin type-1 domain-containing protein 7A-like [Hydr | 82 | 41599 | 2 | 0 | 2 | 0 | 6,5 |
| 156 | gi\|113531039 | chitinase 2 [Hydractinia echinata] | 81 | 48377 | 1 | 1 | 1 | 1 | 2,8 |
| 157 | gi\|828196752 | PREDICTED: probable G-protein coupled receptor 112 isoform X1 [Hydra vulga | 81 | 130717 | 2 | 0 | 2 | 0 | 1,9 |
| 158 | gi\|4757756 | annexin A2 isoform 2 [Homo sapiens] | 80 | 38808 | 2 | 0 | 2 | 0 | 6,2 |
| 159 | gi\|828223895 | PREDICTED: myoferlin [Hydra vulgaris] | 80 | 231950 | 2 | 0 | 2 | 0 | 1,4 |
| 160 | gi\|526117631 | cysteine rich BMP regulator 2 precursor [Hydra vulgaris] | 79 | 132361 | 2 | 0 | 2 | 0 | 2,1 |
| 161 | gi\|31645 | glyceraldehyde-3-phosphate dehydrogenase [Homo sapiens] | 79 | 36202 | 2 | 0 | 2 | 0 | 8,7 |
| 162 | gi\|828189807 | PREDICTED: cysteine and glycine-rich protein 1-like [Hydra vulgaris] | 76 | 11681 | 2 | 0 | 2 | 0 | 10,2 |
| 163 | gi\|736817112 | DNA polymerase III subunit beta [[Eubacterium] nodatum] | 74 | 41468 | 3 | 0 | 2 | 0 | 2,2 |
| 164 | gi\|847168677 | PREDICTED: LOW QUALITY PROTEIN: keratin-3, type I cytoskeletal 51 kDa-like [ | 73 | 52538 | 3 | 0 | 2 | 0 | 3,2 |
| 165 | gi\|449683115 | PREDICTED: astacin-like, partial [Hydra vulgaris] | 73 | 15801 | 4 | 0 | 2 | 0 | 11,6 |
| 166 | gi\|221131483 | PREDICTED: uncharacterized protein LOC100199100 [Hydra vulgaris] | 72 | 20190 | 2 | 0 | 2 | 0 | 10,9 |
| 167 | gi\|449661942 | PREDICTED: arginase-1-like [Hydra vulgaris] | 71 | 36923 | 1 | 1 | 1 | 1 | 2,7 |
| 168 | gi\|3891470 | Chain A, Crystal Structure Of Human Galectin-7 In Complex With Galactosami | 70 | 14992 | 1 | 1 | 1 | 1 | 8,1 |
| 169 | gi\|449673266 | PREDICTED: alkyl/aryl-sulfatase BDS1-like [Hydra vulgaris] | 70 | 67246 | 2 | 0 | 2 | 0 | 4,8 |
| 170 | gi\|302673269 | hypothetical protein SCHCODRAFT_238605 [Schizophyllum commune H4-8] | 70 | 151449 | 3 | 0 | 2 | 0 | 0,7 |
| 171 | gi\|27806789 | transthyretin precursor [Bos taurus] | 69 | 15831 | 2 | 0 | 2 | 0 | 15,6 |
| 172 | gi\|780839549 | hypothetical protein VR70_05230 [Rhodospirillaceae bacterium BRH_c57] | 68 | 122394 | 2 | 1 | 1 | 1 | 1 |
| 173 | gi\|828209114 | PREDICTED: uncharacterized protein LOC101235124 isoform X1 [Hydra vulga | 68 | 54646 | 1 | 1 | 1 | 1 | 3,8 |
| 174 | gi\|47227198 | unnamed protein product [Tetraodon nigroviridis] | 67 | 29312 | 2 | 0 | 2 | 0 | 7,3 |
| 175 | gi\|4204211 | actin-binding protein ABP-280, partial [Hydra vulgaris] | 64 | 24207 | 1 | 0 | 1 | 0 | 8,2 |
| 176 | gi\|449661938 | PREDICTED: single-stranded DNA-binding protein, mitochondrial-like [Hydra v | 64 | 18793 | 1 | 0 | 1 | 0 | 8,4 |

HyWnt3(-) Fraction

| Hit No. | Accession No. | Protein Description | Score | Mass [Da] | Prot Matches | sign Prot Matches | Prot Sequences | sign Prot Sequences | Coverage [%] |
|---|---|---|---|---|---|---|---|---|---|
| 1 | gi\|828208682 | PREDICTED: filamin-A-like [Hydra vulgaris] | 3228 | 294642 | 84 | 19 | 59 | 17 | 25,7 |
| 2 | gi\|11935049 | keratin 1 [Homo sapiens] | 1555 | 66198 | 46 | 16 | 26 | 12 | 35,7 |
| 3 | gi\|375314779 | keratin 1 [Homo sapiens] | 1536 | 66197 | 44 | 14 | 26 | 12 | 35,7 |
| 4 | gi\|28317 | unnamed protein product [Homo sapiens] | 1282 | 59720 | 32 | 8 | 23 | 7 | 42,3 |
| 5 | gi\|828198642 | PREDICTED: uncharacterized protein LOC100198704, partial [Hydra vulgaris] | 1262 | 162069 | 32 | 7 | 24 | 6 | 19,9 |
| 6 | gi\|435476 | cytokeratin 9 [Homo sapiens] | 1225 | 62320 | 31 | 9 | 21 | 7 | 34,2 |
| 7 | gi\|449671849 | PREDICTED: protein disulfide-isomerase A3-like [Hydra vulgaris] | 1196 | 55425 | 37 | 13 | 20 | 9 | 44,3 |
| 8 | gi\|181402 | epidermal cytokeratin 2 [Homo sapiens] | 1119 | 66110 | 27 | 8 | 21 | 7 | 39,7 |
| 9 | gi\|2392071 | Chain A, Crystal Structure Of The Annexin Xii Hexamer | 1082 | 35070 | 34 | 7 | 20 | 5 | 57,5 |
| 10 | gi\|828195098 | PREDICTED: 60 kDa heat shock protein, mitochondrial-like [Hydra vulgaris] | 1036 | 61302 | 21 | 6 | 19 | 6 | 40,2 |
| 11 | gi\|828224124 | PREDICTED: catalase-like [Hydra vulgaris] | 923 | 57689 | 23 | 5 | 19 | 4 | 37,6 |
| 12 | gi\|221108525 | PREDICTED: malate dehydrogenase, mitochondrial-like [Hydra vulgaris] | 896 | 36471 | 22 | 5 | 16 | 5 | 51,8 |
| 13 | gi\|828203840 | PREDICTED: betaine--homocysteine S-methyltransferase 1-like [Hydra vulgaris] | 803 | 44885 | 32 | 8 | 13 | 7 | 28,7 |
| 14 | gi\|449672141 | PREDICTED: glutamate dehydrogenase, mitochondrial-like [Hydra vulgaris] | 796 | 59949 | 21 | 7 | 15 | 6 | 34,1 |
| 15 | gi\|828197727 | PREDICTED: fibronectin type III domain-containing protein-like [Hydra vulgari | 777 | 152931 | 28 | 4 | 16 | 3 | 14,7 |
| 16 | gi\|828201587 | PREDICTED: probable protein disulfide-isomerase A6 [Hydra vulgaris] | 740 | 27489 | 23 | 5 | 14 | 4 | 53,9 |
| 17 | gi\|221115947 | PREDICTED: fructose-bisphosphate aldolase A-like [Hydra vulgaris] | 721 | 39494 | 15 | 7 | 11 | 7 | 37,1 |
| 18 | gi\|221122379 | PREDICTED: staphylococcal nuclease domain-containing protein 1-like [Hydra | 693 | 100759 | 18 | 6 | 13 | 5 | 16,3 |
| 19 | gi\|449679909 | PREDICTED: transketolase-like protein 2 [Hydra vulgaris] | 682 | 68714 | 15 | 4 | 13 | 4 | 31,4 |
| 20 | gi\|828194560 | PREDICTED: contactin-associated protein-like 2 [Hydra vulgaris] | 668 | 138908 | 18 | 3 | 15 | 3 | 14 |
| 21 | gi\|38640805 | cathepsin L-associated protein [Artemia franciscana] | 650 | 34918 | 19 | 6 | 12 | 4 | 49,4 |
| 22 | gi\|565324183 | nucleoside diphosphate kinase-like [Hydra vulgaris] | 642 | 17048 | 31 | 4 | 13 | 3 | 84,1 |
| 23 | gi\|828230018 | PREDICTED: alcohol dehydrogenase [NADP(+)]-like [Hydra vulgaris] | 630 | 35493 | 16 | 4 | 11 | 4 | 56,5 |
| 24 | gi\|1703135 | RecName: Full=Actin, cytoskeletal 3A; AltName: Full=Actin, cytoskeletal IIIA; Fl | 551 | 42162 | 14 | 3 | 12 | 2 | 36,2 |
| 25 | gi\|828223684 | PREDICTED: transaldolase-like [Hydra vulgaris] | 528 | 36670 | 13 | 2 | 11 | 2 | 29,2 |
| 26 | gi\|9739163 | keratin 5 [Homo sapiens] | 527 | 62651 | 16 | 2 | 13 | 2 | 21,4 |
| 27 | gi\|221115109 | PREDICTED: profilin-like [Hydra vulgaris] | 520 | 14833 | 25 | 7 | 8 | 3 | 75,5 |
| 28 | gi\|449667373 | PREDICTED: peptidyl-prolyl cis-trans isomerase B-like [Hydra vulgaris] | 498 | 22428 | 16 | 3 | 9 | 3 | 51,7 |
| 29 | gi\|198139671 | fructose 1,6-bisphosphate aldolase [Artemia franciscana] | 493 | 25865 | 10 | 6 | 7 | 5 | 29,3 |
| 30 | gi\|449667063 | PREDICTED: ADP-ribose pyrophosphatase, mitochondrial-like isoform X2 [Hyd | 491 | 36625 | 13 | 3 | 9 | 3 | 34,7 |
| 31 | gi\|828229950 | PREDICTED: coadhesin-like, partial [Hydra vulgaris] | 487 | 64136 | 12 | 3 | 10 | 3 | 23,3 |
| 32 | gi\|828235386 | PREDICTED: serine/threonine-protein phosphatase 6 regulatory ankyrin repeat | 478 | 105630 | 12 | 2 | 9 | 2 | 10,1 |
| 33 | gi\|1351907 | RecName: Full=Serum albumin; AltName: Full=BSA; AltName: Allergen=Bos d 6 | 466 | 71244 | 10 | 3 | 9 | 3 | 16,3 |
| 34 | gi\|33346945 | ubiquitin/actin fusion protein [Gymnochlora stellata] | 430 | 49580 | 15 | 2 | 9 | 1 | 23,5 |
| 35 | gi\|828223535 | PREDICTED: golgin subfamily B member 1-like [Hydra vulgaris] | 424 | 545146 | 12 | 0 | 12 | 0 | 2,5 |
| 36 | gi\|221112786 | PREDICTED: Na(+)/H(+) exchange regulatory cofactor NHE-RF1-like [Hydra vulga | 414 | 37612 | 12 | 2 | 7 | 2 | 22,1 |
| 37 | gi\|828234048 | PREDICTED: lupus La protein homolog A-like [Hydra vulgaris] | 414 | 44610 | 10 | 3 | 7 | 2 | 22,9 |
| 38 | gi\|221103278 | PREDICTED: fumarate hydratase, mitochondrial-like [Hydra vulgaris] | 411 | 54261 | 11 | 2 | 8 | 2 | 20,5 |
| 39 | gi\|221129526 | PREDICTED: radixin-like [Hydra vulgaris] | 411 | 66644 | 15 | 1 | 9 | 1 | 14,2 |
| 40 | gi\|221132017 | PREDICTED: 78 kDa glucose-regulated protein-like [Hydra vulgaris] | 409 | 74120 | 10 | 1 | 8 | 1 | 13,1 |
| 41 | gi\|146271914 | thrombospondin type 1 repeat-containing protein 2 precursor [Hydra vulgaris | 402 | 102458 | 11 | 1 | 10 | 1 | 11,8 |
| 42 | gi\|828210115 | PREDICTED: glutathione S-transferase Mu 1-like [Hydra vulgaris] | 398 | 13022 | 11 | 3 | 7 | 2 | 57,1 |
| 43 | gi\|221113405 | PREDICTED: chymotrypsin-like elastase family member 3B [Hydra vulgaris] | 395 | 29175 | 13 | 2 | 8 | 2 | 40,3 |
| 44 | gi\|449667073 | PREDICTED: acidic mammalian chitinase-like [Hydra vulgaris] | 390 | 48553 | 12 | 1 | 8 | 1 | 25,1 |
| 45 | gi\|828209655 | PREDICTED: thymocyte nuclear protein 1-like [Hydra vulgaris] | 390 | 29264 | 19 | 1 | 10 | 1 | 37,2 |

| | | | | | | | | | |
|---|---|---|---|---|---|---|---|---|---|
| 46 | gi\|828192702 | PREDICTED: WD repeat-containing protein 1-A-like isoform X1 [Hydra vulgaris] | 387 | 68092 | 11 | 1 | 10 | 1 | 17,2 |
| 47 | gi\|526117559 | four-domain proteases inhibitor-like precursor [Hydra vulgaris] | 386 | 19425 | 11 | 3 | 8 | 2 | 54,2 |
| 48 | gi\|449662397 | PREDICTED: fumarylacetoacetase-like [Hydra vulgaris] | 385 | 46438 | 10 | 1 | 9 | 1 | 24,5 |
| 49 | gi\|449689369 | PREDICTED: protein NipSnap homolog 1-like [Hydra vulgaris] | 383 | 32344 | 11 | 0 | 9 | 0 | 33,3 |
| 50 | gi\|4204211 | actin-binding protein ABP-280, partial [Hydra vulgaris] | 375 | 24207 | 17 | 4 | 8 | 1 | 35,9 |
| 51 | gi\|291406077 | PREDICTED: keratin, type I cytoskeletal 13 [Oryctolagus cuniculus] | 375 | 44693 | 11 | 0 | 8 | 0 | 16,2 |
| 52 | gi\|449692316 | PREDICTED: aldose reductase-like [Hydra vulgaris] | 374 | 13333 | 11 | 3 | 7 | 2 | 76,1 |
| 53 | gi\|221091687 | PREDICTED: triosephosphate isomerase-like [Hydra vulgaris] | 370 | 27190 | 8 | 3 | 7 | 3 | 37,3 |
| 54 | gi\|221130743 | PREDICTED: uncharacterized protein LOC100211495 [Hydra vulgaris] | 370 | 13958 | 13 | 4 | 7 | 3 | 52,4 |
| 55 | gi\|828217704 | PREDICTED: collagen alpha-6(VI) chain-like [Hydra vulgaris] | 368 | 68564 | 7 | 2 | 6 | 2 | 12,6 |
| 56 | gi\|2724046 | beta-actin [Mustela putorius furo] | 367 | 36099 | 9 | 2 | 8 | 1 | 34,1 |
| 57 | gi\|3318722 | Chain E, Leech-Derived Tryptase InhibitorTRYPSIN COMPLEX | 365 | 24142 | 18 | 2 | 6 | 2 | 26 |
| 58 | gi\|828202745 | PREDICTED: uncharacterized protein LOC100202739 isoform X2 [Hydra vulga | 365 | 801426 | 7 | 2 | 7 | 2 | 1,2 |
| 59 | gi\|828220687 | PREDICTED: protein disulfide-isomerase A4-like [Hydra vulgaris] | 365 | 72268 | 7 | 3 | 7 | 3 | 12,9 |
| 60 | gi\|926716364 | PREDICTED: keratin, type I cytoskeletal 10 [Capra hircus] | 362 | 52798 | 10 | 1 | 7 | 1 | 13,5 |
| 61 | gi\|449692014 | PREDICTED: glutathione S-transferase Mu 3-like [Hydra vulgaris] | 357 | 17181 | 13 | 3 | 7 | 2 | 53,4 |
| 62 | gi\|449686976 | PREDICTED: protein disulfide-isomerase A6-like [Hydra vulgaris] | 354 | 48142 | 7 | 2 | 7 | 2 | 19,9 |
| 63 | gi\|221115097 | PREDICTED: glutathione S-transferase-like [Hydra vulgaris] | 343 | 24049 | 13 | 4 | 6 | 2 | 24,3 |
| 64 | gi\|828202697 | PREDICTED: peroxiredoxin-4-like [Hydra vulgaris] | 339 | 27499 | 10 | 1 | 8 | 1 | 30,7 |
| 65 | gi\|449662449 | PREDICTED: polyubiquitin-B [Hydra vulgaris] | 335 | 42564 | 15 | 2 | 8 | 2 | 17,6 |
| 66 | gi\|828199466 | PREDICTED: adenylyl cyclase-associated protein-like [Hydra vulgaris] | 334 | 53995 | 9 | 3 | 6 | 2 | 16,9 |
| 67 | gi\|828205217 | PREDICTED: pirin-like protein [Hydra vulgaris] | 333 | 25924 | 9 | 0 | 8 | 0 | 37,9 |
| 68 | gi\|828192654 | PREDICTED: myosin heavy chain, embryonic smooth muscle isoform-like [Hyd | 330 | 68182 | 6 | 1 | 6 | 1 | 11,3 |
| 69 | gi\|449690552 | PREDICTED: chymotrypsin-like elastase family member 3B [Hydra vulgaris] | 325 | 29089 | 11 | 2 | 7 | 2 | 31,9 |
| 70 | gi\|697450140 | Keratin, type II cytoskeletal 75 [Charadrius vociferus] | 325 | 63147 | 11 | 4 | 6 | 2 | 10,2 |
| 71 | gi\|221109840 | PREDICTED: glucose-6-phosphate isomerase-like, partial [Hydra vulgaris] | 320 | 15636 | 7 | 1 | 7 | 1 | 53,4 |
| 72 | gi\|828218611 | PREDICTED: nucleoside diphosphate kinase A-like [Hydra vulgaris] | 319 | 25885 | 11 | 1 | 7 | 1 | 39,6 |
| 73 | gi\|828229911 | PREDICTED: 2-amino-3-ketobutyrate coenzyme A ligase, mitochondrial-like [H | 316 | 46963 | 9 | 2 | 6 | 1 | 16,8 |
| 74 | gi\|221114999 | PREDICTED: chitinase-3-like protein 1 [Hydra vulgaris] | 313 | 52131 | 8 | 2 | 7 | 2 | 21,3 |
| 75 | gi\|646697674 | Fructose-bisphosphate aldolase [Zootermopsis nevadensis] | 312 | 39761 | 9 | 4 | 5 | 3 | 12,4 |
| 76 | gi\|221121571 | PREDICTED: zinc metalloproteinase nas-15-like [Hydra vulgaris] | 312 | 40888 | 7 | 1 | 6 | 1 | 19,6 |
| 77 | gi\|194772468 | GF20391 [Drosophila ananassae] | 311 | 21231 | 8 | 2 | 6 | 1 | 33,3 |
| 78 | gi\|828199374 | PREDICTED: probable protein disulfide-isomerase A4 [Hydra vulgaris] | 310 | 127080 | 7 | 1 | 7 | 1 | 7,7 |
| 79 | gi\|386848 | keratin [Homo sapiens] | 310 | 51916 | 11 | 1 | 7 | 1 | 13,6 |
| 80 | gi\|567757496 | guanine nucleotide-binding protein subunit beta-like protein [Hydra vulgaris] | 307 | 35804 | 9 | 0 | 7 | 0 | 20,2 |
| 81 | gi\|828234415 | PREDICTED: carbonic anhydrase 7-like [Hydra vulgaris] | 299 | 36918 | 9 | 1 | 7 | 1 | 22,3 |
| 82 | gi\|221132488 | PREDICTED: uncharacterized protein LOC100213474 [Hydra vulgaris] | 297 | 27971 | 8 | 1 | 7 | 1 | 34,9 |
| 83 | gi\|828205187 | PREDICTED: disks large homolog 1-like [Hydra vulgaris] | 297 | 87452 | 8 | 1 | 8 | 1 | 12,9 |
| 84 | gi\|514683975 | heat shock protein 60 [Salpingoeca rosetta] | 296 | 61289 | 6 | 2 | 6 | 2 | 8 |
| 85 | gi\|828222429 | PREDICTED: AP-2 complex subunit alpha-2-like [Hydra vulgaris] | 296 | 103472 | 7 | 1 | 7 | 1 | 8,4 |
| 86 | gi\|828194030 | PREDICTED: uncharacterized protein LOC100200589 [Hydra vulgaris] | 294 | 86226 | 7 | 1 | 7 | 1 | 10,5 |
| 87 | gi\|221118599 | PREDICTED: dolichyl-diphosphooligosaccharide--protein glycosyltransferase s | 293 | 68116 | 6 | 1 | 6 | 1 | 11,2 |
| 88 | gi\|828225443 | PREDICTED: thrombospondin type-1 domain-containing protein 7A-like [Hydr | 291 | 41599 | 7 | 1 | 6 | 1 | 18,7 |
| 89 | gi\|828228866 | PREDICTED: branched-chain-amino-acid aminotransferase, cytosolic-like [Hyd | 288 | 45736 | 9 | 1 | 8 | 1 | 24,1 |
| 90 | gi\|828201601 | PREDICTED: probable methylmalonate-semialdehyde dehydrogenase [acylatin | 277 | 77319 | 6 | 0 | 6 | 0 | 9,1 |
| 91 | gi\|449685905 | PREDICTED: acid ceramidase-like [Hydra vulgaris] | 277 | 42798 | 8 | 1 | 6 | 1 | 20,1 |
| 92 | gi\|829966121 | PREDICTED: LOW QUALITY PROTEIN: polyubiquitin-B [Microcebus murinus] | 275 | 25839 | 14 | 0 | 7 | 0 | 27,5 |
| 93 | gi\|221119142 | PREDICTED: zinc carboxypeptidase-like [Hydra vulgaris] | 275 | 47718 | 8 | 2 | 5 | 2 | 13,5 |

| | | | | | | | | | |
|---|---|---|---|---|---|---|---|---|---|
| 94 | gi\|828213638 | PREDICTED: heterogeneous nuclear ribonucleoprotein A0-like [Hydra vulgaris] | 272 | 33657 | 6 | 1 | 6 | 1 | 22,1 |
| 95 | gi\|828210482 | PREDICTED: branched-chain-amino-acid aminotransferase-like [Hydra vulgaris | 272 | 39139 | 9 | 1 | 7 | 1 | 25,4 |
| 96 | gi\|449673266 | PREDICTED: alkyl/aryl-sulfatase BDS1-like [Hydra vulgaris] | 270 | 67246 | 7 | 0 | 6 | 0 | 15,7 |
| 97 | gi\|449662275 | PREDICTED: thioredoxin-like [Hydra vulgaris] | 267 | 11812 | 9 | 1 | 5 | 1 | 43,8 |
| 98 | gi\|449676410 | PREDICTED: cdc42-interacting protein 4 homolog [Hydra vulgaris] | 264 | 57919 | 7 | 1 | 6 | 1 | 13,8 |
| 99 | gi\|828206617 | PREDICTED: kinectin-like [Hydra vulgaris] | 264 | 113168 | 7 | 0 | 7 | 0 | 7,4 |
| 100 | gi\|565324171 | high mobility group-T protein-like [Hydra vulgaris] | 261 | 20500 | 7 | 1 | 6 | 1 | 29,9 |
| 101 | gi\|221102389 | PREDICTED: protein PRY1-like [Hydra vulgaris] | 260 | 20363 | 8 | 1 | 5 | 1 | 32,8 |
| 102 | gi\|221111162 | PREDICTED: peroxiredoxin-2-like [Hydra vulgaris] | 258 | 22270 | 8 | 1 | 5 | 1 | 20,8 |
| 103 | gi\|221125749 | PREDICTED: putative acetyltransferase DDB_G0275913 [Hydra vulgaris] | 257 | 24335 | 9 | 1 | 6 | 1 | 33,9 |
| 104 | gi\|828218618 | PREDICTED: putative phospholipase B-like 2 [Hydra vulgaris] | 256 | 63709 | 4 | 2 | 4 | 2 | 8,9 |
| 105 | gi\|828214833 | PREDICTED: major vault protein-like [Hydra vulgaris] | 255 | 96304 | 7 | 1 | 6 | 1 | 8,5 |
| 106 | gi\|3046400 | actin 1 [Schmidtea polychroa] | 251 | 8257 | 6 | 1 | 5 | 1 | 53,8 |
| 107 | gi\|221129013 | PREDICTED: protein PRY1-like [Hydra vulgaris] | 249 | 18775 | 5 | 1 | 5 | 1 | 32,5 |
| 108 | gi\|828198329 | PREDICTED: uncharacterized protein LOC100198634 [Hydra vulgaris] | 247 | 166979 | 6 | 1 | 6 | 1 | 4 |
| 109 | gi\|221116483 | PREDICTED: translation elongation factor 2-like [Hydra vulgaris] | 244 | 95554 | 7 | 0 | 7 | 0 | 8,3 |
| 110 | gi\|221131022 | PREDICTED: glucosamine-6-phosphate isomerase 1-like [Hydra vulgaris] | 243 | 30402 | 4 | 1 | 4 | 1 | 22,1 |
| 111 | gi\|828234147 | PREDICTED: cystatin-A-like [Hydra vulgaris] | 242 | 11384 | 6 | 2 | 4 | 2 | 52,5 |
| 112 | gi\|544430751 | PREDICTED: uncharacterized protein LOC102118097 [Macaca fascicularis] | 238 | 61529 | 7 | 0 | 6 | 0 | 10,4 |
| 113 | gi\|449678439 | PREDICTED: peptidyl-prolyl cis-trans isomerase 5-like [Hydra vulgaris] | 235 | 25229 | 5 | 2 | 5 | 2 | 24,9 |
| 114 | gi\|315439538 | vitellogenin-superoxide dismutase fusion protein [Artemia parthenogenetica] | 235 | 249350 | 6 | 0 | 6 | 0 | 3,4 |
| 115 | gi\|830260001 | tight junction protein ZO-2-like [Hydra vulgaris] | 232 | 191163 | 6 | 2 | 4 | 1 | 2,9 |
| 116 | gi\|1330252 | translation elongation factor 1 alpha [Hydra vulgaris] | 231 | 51200 | 5 | 0 | 5 | 0 | 10,5 |
| 117 | gi\|449669260 | PREDICTED: eukaryotic translation initiation factor 3 subunit B-like [Hydra vul | 231 | 80964 | 6 | 1 | 4 | 1 | 6,9 |
| 118 | gi\|449670492 | PREDICTED: elongation factor 1-gamma-like [Hydra vulgaris] | 228 | 49912 | 6 | 0 | 5 | 0 | 11,5 |
| 119 | gi\|828201918 | PREDICTED: glyceraldehyde-3-phosphate dehydrogenase [Hydra vulgaris] | 226 | 36793 | 8 | 1 | 5 | 1 | 18,2 |
| 120 | gi\|449665286 | PREDICTED: nucleoredoxin-like protein 2 [Hydra vulgaris] | 224 | 16201 | 4 | 1 | 4 | 1 | 35,9 |
| 121 | gi\|828230255 | PREDICTED: glucose-6-phosphate isomerase-like, partial [Hydra vulgaris] | 220 | 39603 | 5 | 0 | 5 | 0 | 22,6 |
| 122 | gi\|828224104 | PREDICTED: beta-glucuronidase-like [Hydra vulgaris] | 218 | 73149 | 6 | 0 | 6 | 0 | 11 |
| 123 | gi\|828213796 | PREDICTED: probable aminopeptidase NPEPL1 [Hydra vulgaris] | 217 | 54416 | 5 | 0 | 5 | 0 | 11,2 |
| 124 | gi\|449666857 | PREDICTED: lysosomal aspartic protease-like [Hydra vulgaris] | 212 | 42777 | 6 | 1 | 4 | 1 | 12,7 |
| 125 | gi\|828218104 | PREDICTED: myosin-10-like [Hydra vulgaris] | 202 | 225409 | 5 | 0 | 5 | 0 | 2,7 |
| 126 | gi\|221121754 | PREDICTED: very long-chain specific acyl-CoA dehydrogenase, mitochondrial-l | 202 | 68420 | 6 | 1 | 6 | 1 | 10,2 |
| 127 | gi\|828215949 | PREDICTED: alpha-L-fucosidase-like isoform X1 [Hydra vulgaris] | 201 | 56266 | 5 | 1 | 5 | 1 | 9,4 |
| 128 | gi\|828228888 | PREDICTED: stress-induced-phosphoprotein 1-like [Hydra vulgaris] | 197 | 61796 | 4 | 0 | 4 | 0 | 8,2 |
| 129 | gi\|828189807 | PREDICTED: cysteine and glycine-rich protein 1-like [Hydra vulgaris] | 196 | 11681 | 5 | 1 | 3 | 1 | 25,9 |
| 130 | gi\|449669354 | PREDICTED: oxygen-dependent coproporphyrinogen-III oxidase-like [Hydra vu | 194 | 39728 | 5 | 1 | 5 | 1 | 14,4 |
| 131 | gi\|828213548 | PREDICTED: uncharacterized protein LOC105845774 [Hydra vulgaris] | 193 | 19785 | 4 | 0 | 4 | 0 | 27,7 |
| 132 | gi\|221125639 | PREDICTED: alanine--glyoxylate aminotransferase 2, mitochondrial-like [Hydra | 192 | 55680 | 6 | 0 | 6 | 0 | 12,6 |
| 133 | gi\|221132111 | PREDICTED: hypoxia up-regulated protein 1-like [Hydra vulgaris] | 191 | 112944 | 4 | 0 | 4 | 0 | 4,2 |
| 134 | gi\|449684469 | PREDICTED: stress-70 protein, mitochondrial-like [Hydra vulgaris] | 190 | 72978 | 6 | 1 | 4 | 1 | 7,5 |
| 135 | gi\|449669580 | PREDICTED: peptidyl-prolyl cis-trans isomerase FKBP9-like [Hydra vulgaris] | 189 | 30984 | 7 | 0 | 6 | 0 | 23,1 |
| 136 | gi\|828221194 | PREDICTED: fibronectin type III domain-containing protein-like [Hydra vulgari | 188 | 143336 | 4 | 1 | 4 | 1 | 4 |
| 137 | gi\|74483475 | elongation factor 1 alpha [Ceratinia neso] | 187 | 45127 | 4 | 0 | 4 | 0 | 9,7 |
| 138 | gi\|221114449 | PREDICTED: nuclear polyadenylated RNA-binding protein 4-like [Hydra vulgari | 185 | 43864 | 6 | 1 | 4 | 1 | 10,3 |
| 139 | gi\|221111160 | PREDICTED: peroxiredoxin-1-like [Hydra vulgaris] | 184 | 26936 | 7 | 1 | 4 | 1 | 15,5 |
| 140 | gi\|828220681 | PREDICTED: macrophage migration inhibitory factor-like [Hydra vulgaris] | 182 | 13896 | 4 | 2 | 3 | 2 | 18,7 |
| 141 | gi\|321460290 | cytosolic malate dehydrogenase [Daphnia pulex] | 181 | 35988 | 4 | 0 | 4 | 0 | 15,3 |

| | | | | | | | | | |
|---|---|---|---|---|---|---|---|---|---|
| 142 | gi\|209156284 | Heat shock 70 kDa protein [Salmo salar] | 175 | 71189 | 6 | 0 | 5 | 0 | 6,5 |
| 143 | gi\|828197619 | PREDICTED: lysosomal alpha-mannosidase-like [Hydra vulgaris] | 175 | 49198 | 4 | 0 | 4 | 0 | 10,2 |
| 144 | gi\|449678000 | PREDICTED: S-methyl-5'-thioadenosine phosphorylase-like [Hydra vulgaris] | 173 | 30317 | 2 | 2 | 2 | 2 | 11,4 |
| 145 | gi\|221132295 | PREDICTED: glutathione S-transferase-like [Hydra vulgaris] | 166 | 23892 | 6 | 0 | 4 | 0 | 12 |
| 146 | gi\|828192764 | PREDICTED: aminopeptidase N-like [Hydra vulgaris] | 166 | 104641 | 4 | 0 | 4 | 0 | 4,4 |
| 147 | gi\|148229965 | heat shock 70kDa protein 2 [Xenopus laevis] | 166 | 69803 | 4 | 0 | 4 | 0 | 5,2 |
| 148 | gi\|221126616 | PREDICTED: THO complex subunit 4-like [Hydra vulgaris] | 166 | 26671 | 7 | 0 | 5 | 0 | 17,6 |
| 149 | gi\|754342395 | heat shock protein 70 [Capsaspora owczarzaki ATCC 30864] | 164 | 72508 | 6 | 0 | 5 | 0 | 6,9 |
| 150 | gi\|291234001 | PREDICTED: late histone H2B.L3-like isoform X1 [Saccoglossus kowalevskii] | 162 | 13838 | 4 | 2 | 3 | 1 | 28 |
| 151 | gi\|449684402 | PREDICTED: endochitinase 1-like isoform X1 [Hydra vulgaris] | 162 | 52676 | 3 | 1 | 3 | 1 | 8,5 |
| 152 | gi\|449678353 | PREDICTED: bandaporin-like [Hydra vulgaris] | 161 | 22016 | 4 | 1 | 3 | 1 | 19,2 |
| 153 | gi\|449683097 | PREDICTED: peptidyl-prolyl cis-trans isomerase-like [Hydra vulgaris] | 158 | 17530 | 3 | 0 | 3 | 0 | 20,5 |
| 154 | gi\|449687222 | PREDICTED: allograft inflammatory factor 1-like [Hydra vulgaris] | 155 | 17334 | 3 | 1 | 3 | 1 | 29,3 |
| 155 | gi\|221126057 | PREDICTED: antistasin-like [Hydra vulgaris] | 154 | 26072 | 11 | 2 | 3 | 1 | 15,1 |
| 156 | gi\|831771570 | hypothetical protein SAMD00019534_067750 [Acytostelium subglobosum L | 150 | 71282 | 5 | 0 | 4 | 0 | 6,1 |
| 157 | gi\|449661942 | PREDICTED: arginase-1-like [Hydra vulgaris] | 150 | 36923 | 4 | 1 | 3 | 1 | 10,3 |
| 158 | gi\|828213619 | PREDICTED: gelsolin-like protein 1 [Hydra vulgaris] | 147 | 40776 | 5 | 0 | 4 | 0 | 14,1 |
| 159 | gi\|828220786 | PREDICTED: rho GDP-dissociation inhibitor 1-like [Hydra vulgaris] | 147 | 22744 | 5 | 0 | 4 | 0 | 18,2 |
| 160 | gi\|828191663 | PREDICTED: neogenin-like [Hydra vulgaris] | 145 | 209395 | 4 | 0 | 4 | 0 | 2,8 |
| 161 | gi\|478520725 | PREDICTED: keratin, type I cytoskeletal 26 [Ceratotherium simum simum] | 144 | 52420 | 6 | 1 | 3 | 1 | 5,6 |
| 162 | gi\|449679776 | PREDICTED: failed axon connections homolog [Hydra vulgaris] | 143 | 51908 | 3 | 1 | 3 | 1 | 7,7 |
| 163 | gi\|83595133 | manganese superoxide dismutase [Hydra vulgaris] | 142 | 24334 | 4 | 0 | 3 | 0 | 12,8 |
| 164 | gi\|13111486 | elongation factor-2, partial [Artemia salina] | 141 | 71657 | 4 | 1 | 3 | 1 | 6,2 |
| 165 | gi\|221090861 | PREDICTED: cathepsin L1-like [Hydra vulgaris] | 141 | 36531 | 3 | 1 | 3 | 1 | 9,6 |
| 166 | gi\|526117746 | flp protein [Hydra vulgaris] | 141 | 14995 | 3 | 0 | 3 | 0 | 26,2 |
| 167 | gi\|701307518 | PREDICTED: LOW QUALITY PROTEIN: keratin, type II cytoskeletal cochleal-like [ | 139 | 54856 | 5 | 0 | 4 | 0 | 6,8 |
| 168 | gi\|221120289 | PREDICTED: uncharacterized protein LOC100199298 isoform X1 [Hydra vulga | 138 | 18479 | 3 | 0 | 3 | 0 | 17,6 |
| 169 | gi\|449666332 | PREDICTED: zinc metalloproteinase nas-6-like [Hydra vulgaris] | 136 | 33291 | 4 | 0 | 4 | 0 | 16 |
| 170 | gi\|526117401 | PPOD2 peroxidase-like precursor [Hydra vulgaris] | 135 | 32661 | 2 | 1 | 2 | 1 | 8 |
| 171 | gi\|449667021 | PREDICTED: zinc metalloproteinase nas-14-like [Hydra vulgaris] | 135 | 20331 | 3 | 0 | 3 | 0 | 16,6 |
| 172 | gi\|76560210 | elongation factor 1 alpha [Colletes halophilus] | 135 | 8539 | 3 | 0 | 3 | 0 | 25 |
| 173 | gi\|221091838 | PREDICTED: calcyphosin-like protein [Hydra vulgaris] | 134 | 21273 | 3 | 1 | 3 | 1 | 14,7 |
| 174 | gi\|221124062 | PREDICTED: heme-binding protein 1-like [Hydra vulgaris] | 134 | 29921 | 5 | 0 | 4 | 0 | 14,8 |
| 175 | gi\|828204847 | PREDICTED: urocanate hydratase-like [Hydra vulgaris] | 134 | 75467 | 3 | 1 | 3 | 1 | 4,9 |
| 176 | gi\|526117507 | kazal-type serine protease inihibitor 2 precursor [Hydra vulgaris] | 134 | 19400 | 5 | 0 | 4 | 0 | 23,2 |
| 177 | gi\|221109761 | PREDICTED: uncharacterized protein LOC100200582 [Hydra vulgaris] | 131 | 21912 | 3 | 0 | 3 | 0 | 18,5 |
| 178 | gi\|110433182 | heat shock protein [Bursaphelenchus xylophilus] | 130 | 70411 | 4 | 0 | 4 | 0 | 6,2 |
| 179 | gi\|114389 | RecName: Full=Sodium/potassium-transporting ATPase subunit beta; AltName | 129 | 36274 | 4 | 0 | 3 | 0 | 8,6 |
| 180 | gi\|828232304 | PREDICTED: blastula protease 10-like [Hydra vulgaris] | 129 | 27019 | 3 | 0 | 3 | 0 | 12,6 |
| 181 | gi\|221123276 | PREDICTED: omega-amidase NIT2-like [Hydra vulgaris] | 129 | 30745 | 4 | 0 | 4 | 0 | 16,2 |
| 182 | gi\|828195143 | PREDICTED: pre-mRNA-processing factor 19-like [Hydra vulgaris] | 127 | 55883 | 3 | 0 | 3 | 0 | 6,3 |
| 183 | gi\|828194138 | PREDICTED: dystonin-like, partial [Hydra vulgaris] | 123 | 381036 | 3 | 0 | 3 | 0 | 0,8 |
| 184 | gi\|828224045 | PREDICTED: uncharacterized protein LOC100212316 [Hydra vulgaris] | 122 | 91458 | 4 | 0 | 4 | 0 | 4,3 |
| 185 | gi\|449686817 | PREDICTED: 40S ribosomal protein S12-like [Hydra vulgaris] | 121 | 16472 | 2 | 1 | 2 | 1 | 14,3 |
| 186 | gi\|449679956 | PREDICTED: gamma-aminobutyric acid receptor-associated protein-like 2 [Hyc | 120 | 13699 | 4 | 0 | 4 | 0 | 26,3 |
| 187 | gi\|828227420 | PREDICTED: LOW QUALITY PROTEIN: asparagine--tRNA ligase, cytoplasmic-like | 120 | 64826 | 3 | 0 | 2 | 0 | 4,8 |
| 188 | gi\|15072473 | peroxidase ppod2 [Hydra vulgaris] | 119 | 33192 | 3 | 1 | 2 | 1 | 8,5 |
| 189 | gi\|742871900 | hypothetical protein [Halocynthiibacter sp. PAMC 20958] | 116 | 50662 | 2 | 0 | 2 | 0 | 2 |

| | | | | | | | | | |
|---|---|---|---|---|---|---|---|---|---|
| 190 | gi\|559183818 | Heat shock protein 70, partial [Giardia intestinalis] | 116 | 48333 | 3 | 0 | 3 | 0 | 4,6 |
| 191 | gi\|449671399 | PREDICTED: actin-related protein 2/3 complex subunit 2-like isoform X2 [Hydr | 115 | 34480 | 2 | 1 | 2 | 1 | 9,5 |
| 192 | gi\|828201578 | PREDICTED: purine nucleoside phosphorylase-like, partial [Hydra vulgaris] | 115 | 28573 | 2 | 1 | 2 | 1 | 9,9 |
| 193 | gi\|828227874 | PREDICTED: uncharacterized protein LOC105848518, partial [Hydra vulgaris] | 115 | 14191 | 4 | 0 | 4 | 0 | 29,7 |
| 194 | gi\|828224366 | PREDICTED: xaa-Pro dipeptidase-like [Hydra vulgaris] | 114 | 55873 | 3 | 0 | 3 | 0 | 5,3 |
| 195 | gi\|828225026 | PREDICTED: peptidase M20 domain-containing protein 2-like [Hydra vulgaris] | 113 | 58925 | 3 | 0 | 3 | 0 | 4,9 |
| 196 | gi\|449689149 | PREDICTED: uncharacterized protein LOC100208770, partial [Hydra vulgaris] | 113 | 23100 | 3 | 0 | 3 | 0 | 15,7 |
| 197 | gi\|828195701 | PREDICTED: kynurenine--oxoglutarate transaminase-like [Hydra vulgaris] | 112 | 21895 | 1 | 1 | 1 | 1 | 10,7 |
| 198 | gi\|449670322 | PREDICTED: dipeptidyl peptidase 1-like [Hydra vulgaris] | 112 | 51835 | 3 | 0 | 3 | 0 | 7 |
| 199 | gi\|221111809 | PREDICTED: carbonic anhydrase 2-like [Hydra vulgaris] | 110 | 34574 | 3 | 0 | 3 | 0 | 7,7 |
| 200 | gi\|221113581 | PREDICTED: protein DEK-like [Hydra vulgaris] | 109 | 41907 | 3 | 0 | 3 | 0 | 7,3 |
| 201 | gi\|221130733 | PREDICTED: astacin-like metalloprotease toxin 5 [Hydra vulgaris] | 108 | 27816 | 2 | 1 | 2 | 1 | 9,7 |
| 202 | gi\|828197209 | PREDICTED: annexin A4-like [Hydra vulgaris] | 106 | 66184 | 3 | 1 | 3 | 1 | 4,4 |
| 203 | gi\|828211213 | PREDICTED: heterogeneous nuclear ribonucleoprotein A/B-like [Hydra vulgaris | 106 | 19419 | 2 | 1 | 2 | 1 | 14,8 |
| 204 | gi\|828197199 | PREDICTED: uncharacterized protein LOC101236102 [Hydra vulgaris] | 106 | 66420 | 4 | 0 | 3 | 0 | 5,9 |
| 205 | gi\|510863409 | peptidyl-prolyl cis-trans isomerase, cyclophilin-type [Ancylostoma ceylanicum | 105 | 16883 | 8 | 0 | 3 | 0 | 9,7 |
| 206 | gi\|221121520 | PREDICTED: agmatinase, mitochondrial-like [Hydra vulgaris] | 105 | 34037 | 2 | 1 | 2 | 1 | 9,1 |
| 207 | gi\|449670247 | PREDICTED: probable inactive purple acid phosphatase 2 [Hydra vulgaris] | 104 | 67196 | 2 | 1 | 2 | 1 | 3,4 |
| 208 | gi\|828218463 | PREDICTED: uncharacterized protein LOC101241534 [Hydra vulgaris] | 103 | 142054 | 3 | 0 | 3 | 0 | 2,5 |
| 209 | gi\|449678564 | PREDICTED: lysosome-associated membrane glycoprotein 1-like [Hydra vulgari | 103 | 21879 | 2 | 0 | 2 | 0 | 10,7 |
| 210 | gi\|221131162 | PREDICTED: ATP synthase subunit alpha, mitochondrial [Hydra vulgaris] | 103 | 59027 | 3 | 0 | 3 | 0 | 6,2 |
| 211 | gi\|449665331 | PREDICTED: contactin-associated protein-like 5 [Hydra vulgaris] | 102 | 136592 | 3 | 0 | 3 | 0 | 3,2 |
| 212 | gi\|46909251 | ATP synthase beta subunit, partial [Obelia sp. KJP-2004] | 102 | 46198 | 2 | 0 | 2 | 0 | 7,3 |
| 213 | gi\|828195674 | PREDICTED: gamma-glutamyltranspeptidase 1-like [Hydra vulgaris] | 102 | 67827 | 3 | 0 | 3 | 0 | 7,5 |
| 214 | gi\|526117489 | kazal-type serine protease inihibitor 3 precursor [Hydra vulgaris] | 102 | 19438 | 5 | 0 | 3 | 0 | 19 |
| 215 | gi\|221131289 | PREDICTED: methylmalonyl-CoA epimerase, mitochondrial-like [Hydra vulgari | 101 | 18258 | 2 | 0 | 2 | 0 | 18,8 |
| 216 | gi\|828196768 | PREDICTED: calsequestrin-2-like [Hydra vulgaris] | 101 | 46695 | 3 | 0 | 3 | 0 | 9,2 |
| 217 | gi\|221120850 | PREDICTED: dihydropteridine reductase-like [Hydra vulgaris] | 100 | 25220 | 3 | 1 | 2 | 1 | 11,9 |
| 218 | gi\|526117377 | caspase 7 [Hydra vulgaris] | 100 | 47366 | 3 | 0 | 3 | 0 | 7,4 |
| 219 | gi\|449664868 | PREDICTED: threonine--tRNA ligase, cytoplasmic-like isoform X1 [Hydra vulgar | 100 | 83661 | 3 | 0 | 3 | 0 | 3,6 |
| 220 | gi\|449676841 | PREDICTED: putative acyl-coenzyme A oxidase 3.2, peroxisomal [Hydra vulgari | 100 | 72300 | 3 | 0 | 3 | 0 | 5,1 |
| 221 | gi\|828190890 | PREDICTED: 60S acidic ribosomal protein P2-like [Hydra vulgaris] | 99 | 11782 | 3 | 1 | 2 | 1 | 21,6 |
| 222 | gi\|449689337 | PREDICTED: uncharacterized protein LOC100212684, partial [Hydra vulgaris] | 99 | 24278 | 1 | 1 | 1 | 1 | 8,2 |
| 223 | gi\|221122769 | PREDICTED: uncharacterized protein LOC100209607 [Hydra vulgaris] | 98 | 25274 | 3 | 1 | 2 | 1 | 13,8 |
| 224 | gi\|538775593 | piwi-like protein HyWI [Hydra vulgaris] | 98 | 101873 | 3 | 0 | 3 | 0 | 2,8 |
| 225 | gi\|449680876 | PREDICTED: endochitinase 4-like [Hydra vulgaris] | 98 | 52510 | 2 | 0 | 2 | 0 | 5,8 |
| 226 | gi\|156402513 | predicted protein [Nematostella vectensis] | 97 | 21970 | 2 | 0 | 2 | 0 | 9,5 |
| 227 | gi\|221121832 | PREDICTED: multiple inositol polyphosphate phosphatase 1-like [Hydra vulgar | 96 | 50712 | 2 | 0 | 2 | 0 | 6 |
| 228 | gi\|221126625 | PREDICTED: polypyrimidine tract-binding protein 1-like [Hydra vulgaris] | 96 | 56616 | 1 | 1 | 1 | 1 | 3,2 |
| 229 | gi\|449680926 | PREDICTED: nuclear transport factor 2-like [Hydra vulgaris] | 95 | 13926 | 2 | 0 | 2 | 0 | 17,6 |
| 230 | gi\|440200331 | triosephosphate isomerase, partial [Odites leucostola] | 94 | 16181 | 2 | 0 | 2 | 0 | 13,5 |
| 231 | gi\|17137630 | cytoplasmic dynein light chain 2, isoform A [Drosophila melanogaster] | 94 | 10465 | 2 | 0 | 2 | 0 | 24,7 |
| 232 | gi\|449687420 | PREDICTED: uncharacterized protein LOC100205832 isoform X1 [Hydra vulga | 93 | 24618 | 4 | 0 | 2 | 0 | 8,4 |
| 233 | gi\|221123418 | PREDICTED: eukaryotic translation initiation factor 4H-like [Hydra vulgaris] | 93 | 30810 | 1 | 1 | 1 | 1 | 5,8 |
| 234 | gi\|828214768 | PREDICTED: delta-1-pyrroline-5-carboxylate dehydrogenase, mitochondrial-li | 93 | 40600 | 3 | 0 | 3 | 0 | 10,6 |
| 235 | gi\|449680259 | PREDICTED: glutathione peroxidase 2-like [Hydra vulgaris] | 92 | 23459 | 3 | 0 | 3 | 0 | 13,8 |
| 236 | gi\|449687197 | PREDICTED: golgi-associated plant pathogenesis-related protein 1-like [Hydra v | 92 | 29150 | 3 | 0 | 2 | 0 | 8,6 |
| 237 | gi\|828204323 | PREDICTED: hemicentin-2-like isoform X1 [Hydra vulgaris] | 92 | 158275 | 2 | 0 | 2 | 0 | 1,6 |

| | | | | | | | | | |
|---|---|---|---|---|---|---|---|---|---|
| 238 | gi\|449684745 | PREDICTED: glutathione S-transferase-like [Hydra vulgaris] | 90 | 24182 | 2 | 0 | 2 | 0 | 11,4 |
| 239 | gi\|497899026 | MULTISPECIES: hypothetical protein [Pseudomonas] | 90 | 25538 | 2 | 0 | 2 | 0 | 8 |
| 240 | gi\|449666254 | PREDICTED: glutamic acid-rich protein-like [Hydra vulgaris] | 90 | 44127 | 1 | 1 | 1 | 1 | 3,9 |
| 241 | gi\|449691823 | PREDICTED: dipeptidyl peptidase 3-like, partial [Hydra vulgaris] | 89 | 33965 | 2 | 0 | 2 | 0 | 7,7 |
| 242 | gi\|221132768 | PREDICTED: RNA polymerase II transcriptional coactivator-like [Hydra vulgaris | 89 | 13128 | 3 | 0 | 3 | 0 | 15,7 |
| 243 | gi\|221114177 | PREDICTED: alpha-crystallin A chain-like [Hydra vulgaris] | 88 | 26553 | 2 | 0 | 2 | 0 | 9,4 |
| 244 | gi\|828192317 | PREDICTED: peptidyl-prolyl cis-trans isomerase H-like [Hydra vulgaris] | 87 | 17635 | 2 | 0 | 2 | 0 | 14,5 |
| 245 | gi\|449692187 | PREDICTED: purine nucleoside phosphorylase-like, partial [Hydra vulgaris] | 87 | 16840 | 2 | 0 | 2 | 0 | 12,8 |
| 246 | gi\|501292293 | myosin heavy chain [Riptortus pedestris] | 87 | 167897 | 2 | 0 | 2 | 0 | 1,4 |
| 247 | gi\|493710903 | hypothetical protein [Providencia alcalifaciens] | 87 | 14905 | 2 | 0 | 2 | 0 | 10,8 |
| 248 | gi\|551617245 | hypothetical protein EMIHUDRAFT_423268 [Emiliania huxleyi CCMP1516] | 87 | 56358 | 2 | 0 | 2 | 0 | 3,2 |
| 249 | gi\|449682262 | PREDICTED: zinc metalloproteinase nas-13-like [Hydra vulgaris] | 87 | 58069 | 3 | 0 | 3 | 0 | 6 |
| 250 | gi\|221113277 | PREDICTED: uncharacterized protein LOC100214198 [Hydra vulgaris] | 86 | 30392 | 4 | 0 | 2 | 0 | 7,8 |
| 251 | gi\|828220298 | PREDICTED: ena/VASP-like protein [Hydra vulgaris] | 86 | 39847 | 3 | 0 | 2 | 0 | 3,7 |
| 252 | gi\|828206951 | PREDICTED: uncharacterized protein LOC100208668 isoform X1 [Hydra vulga | 85 | 253566 | 3 | 0 | 2 | 0 | 2 |
| 253 | gi\|449689073 | PREDICTED: ornithine aminotransferase, mitochondrial-like [Hydra vulgaris] | 85 | 48365 | 2 | 0 | 2 | 0 | 7 |
| 254 | gi\|766939774 | PREDICTED: triosephosphate isomerase [Ceratosolen solmsi marchali] | 85 | 27658 | 3 | 0 | 3 | 0 | 7,7 |
| 255 | gi\|828222781 | PREDICTED: hypoxanthine-guanine phosphoribosyltransferase-like [Hydra vul | 85 | 24651 | 2 | 0 | 2 | 0 | 9,7 |
| 256 | gi\|221125651 | PREDICTED: small nuclear ribonucleoprotein Sm D1-like [Hydra vulgaris] | 85 | 13631 | 2 | 0 | 2 | 0 | 24,2 |
| 257 | gi\|487975987 | MULTISPECIES: SnoaL-like domain protein [Acinetobacter calcoaceticus/baum | 85 | 20833 | 2 | 0 | 2 | 0 | 5,1 |
| 258 | gi\|685831842 | Annexin family and Annexin repeat-containing protein [Strongyloides ratti] | 84 | 36712 | 2 | 2 | 1 | 1 | 3,4 |
| 259 | gi\|828206370 | PREDICTED: epidermal growth factor receptor substrate 15-like 1 [Hydra vulga | 84 | 89358 | 2 | 0 | 2 | 0 | 2,6 |
| 260 | gi\|768420965 | PREDICTED: peptidyl-prolyl cis-trans isomerase B [Plutella xylostella] | 83 | 22092 | 3 | 0 | 2 | 0 | 9,9 |
| 261 | gi\|32532 | unnamed protein product [Homo sapiens] | 83 | 26932 | 1 | 1 | 1 | 1 | 5,3 |
| 262 | gi\|28194281 | ubiquitin extension protein [Heterodera glycines] | 82 | 11762 | 6 | 0 | 2 | 0 | 21 |
| 263 | gi\|221103804 | PREDICTED: probable glutathione S-transferase 7 [Hydra vulgaris] | 82 | 24521 | 1 | 1 | 1 | 1 | 7 |
| 264 | gi\|828217423 | PREDICTED: glutamate decarboxylase 2-like isoform X1 [Hydra vulgaris] | 82 | 61072 | 1 | 1 | 1 | 1 | 2,8 |
| 265 | gi\|449666836 | PREDICTED: malectin-A-like [Hydra vulgaris] | 81 | 32778 | 2 | 0 | 2 | 0 | 8,5 |
| 266 | gi\|828211197 | PREDICTED: sorting nexin-32-like [Hydra vulgaris] | 81 | 48466 | 2 | 0 | 2 | 0 | 5,4 |
| 267 | gi\|459367722 | hypothetical protein G210_3086 [Candida maltosa Xu316] | 80 | 74058 | 2 | 0 | 2 | 0 | 1,2 |
| 268 | gi\|899147016 | peroxiredoxin 3 [Esox lucius] | 80 | 27673 | 3 | 0 | 2 | 0 | 7,6 |
| 269 | gi\|828207749 | PREDICTED: N-acyl-phosphatidylethanolamine-hydrolyzing phospholipase D-l | 80 | 52007 | 2 | 0 | 2 | 0 | 3,8 |
| 270 | gi\|828212124 | PREDICTED: MAM and LDL-receptor class A domain-containing protein 1-like [ | 79 | 770950 | 2 | 0 | 2 | 0 | 0,3 |
| 271 | gi\|828192461 | PREDICTED: aspartyl aminopeptidase-like [Hydra vulgaris] | 79 | 51537 | 2 | 0 | 2 | 0 | 4,4 |
| 272 | gi\|999604 | Chain A, Crystallographic Studies On A Family Of Cellular Lipophilic Transport | 79 | 14923 | 2 | 1 | 1 | 1 | 9,2 |
| 273 | gi\|294882533 | Metal homeostasis factor ATX1, putative [Perkinsus marinus ATCC 50983] | 78 | 7429 | 1 | 1 | 1 | 1 | 17,6 |
| 274 | gi\|449679798 | PREDICTED: U1 small nuclear ribonucleoprotein A-like [Hydra vulgaris] | 78 | 25255 | 3 | 2 | 1 | 1 | 5,9 |
| 275 | gi\|890692673 | GntR family transcriptional regulator [Photobacterium swingsii] | 78 | 25836 | 1 | 1 | 1 | 1 | 4 |
| 276 | gi\|828205429 | PREDICTED: uncharacterized protein LOC100204510 [Hydra vulgaris] | 77 | 134052 | 2 | 0 | 2 | 0 | 2,3 |
| 277 | gi\|221126681 | PREDICTED: prefoldin subunit 5-like [Hydra vulgaris] | 77 | 18675 | 1 | 1 | 1 | 1 | 7,7 |
| 278 | gi\|449663959 | PREDICTED: acylpyruvase FAHD1, mitochondrial-like [Hydra vulgaris] | 76 | 23816 | 2 | 0 | 2 | 0 | 14,9 |
| 279 | gi\|443702382 | hypothetical protein CAPTEDRAFT_177200 [Capitella teleta] | 76 | 23316 | 1 | 1 | 1 | 1 | 6 |
| 280 | gi\|449662629 | PREDICTED: iodotyrosine dehalogenase 1-like [Hydra vulgaris] | 76 | 31350 | 2 | 0 | 2 | 0 | 10,7 |
| 281 | gi\|828197526 | PREDICTED: malate dehydrogenase, cytoplasmic-like [Hydra vulgaris] | 76 | 37041 | 3 | 0 | 3 | 0 | 9,6 |
| 282 | gi\|221123857 | PREDICTED: lactoylglutathione lyase-like [Hydra vulgaris] | 75 | 20110 | 1 | 1 | 1 | 1 | 11 |
| 283 | gi\|221131112 | PREDICTED: prefoldin subunit 6-like [Hydra vulgaris] | 75 | 15339 | 2 | 0 | 2 | 0 | 21,5 |
| 284 | gi\|553309896 | polysaccharide pyruvyl transferase CsaB [Peptoniphilus sp. BV3C26] | 75 | 41687 | 3 | 0 | 2 | 0 | 3,8 |
| 285 | gi\|221130032 | PREDICTED: isochorismatase domain-containing protein 2, mitochondrial-like | 75 | 22754 | 1 | 1 | 1 | 1 | 6,5 |

| | | | | | | | | | |
|---|---|---|---|---|---|---|---|---|---|
| 286 | gi\|449679397 | PREDICTED: uncharacterized protein LOC100197967 [Hydra vulgaris] | 75 | 40545 | 3 | 0 | 2 | 0 | 4,2 |
| 287 | gi\|449671578 | PREDICTED: S-crystallin 4-like [Hydra vulgaris] | 74 | 23979 | 2 | 0 | 2 | 0 | 7,7 |
| 288 | gi\|20145612 | putative actin, partial [Hydractinia echinata] | 74 | 8589 | 3 | 0 | 3 | 0 | 49,4 |
| 289 | gi\|828225023 | PREDICTED: peptidase M20 domain-containing protein 2-like, partial [Hydra v | 74 | 34646 | 3 | 0 | 3 | 0 | 7,5 |
| 290 | gi\|156336944 | hypothetical protein NEMVEDRAFT_v1g150407 [Nematostella vectensis] | 74 | 7062 | 2 | 0 | 2 | 0 | 29,7 |
| 291 | gi\|165979176 | Cu-Zn superoxide dismutase, partial [Rhizophagus proliferus] | 73 | 12709 | 1 | 1 | 1 | 1 | 8,3 |
| 292 | gi\|221124690 | PREDICTED: endoplasmin-like [Hydra vulgaris] | 71 | 94322 | 2 | 0 | 2 | 0 | 2,4 |
| 293 | gi\|828235608 | PREDICTED: cytosol aminopeptidase-like [Hydra vulgaris] | 70 | 56366 | 2 | 0 | 2 | 0 | 4,1 |
| 294 | gi\|330796823 | replication factor C subunit [Dictyostelium purpureum] | 70 | 55856 | 2 | 0 | 2 | 0 | 3 |
| 295 | gi\|828203837 | PREDICTED: dimethylglycine dehydrogenase, mitochondrial-like [Hydra vulgar | 70 | 96756 | 2 | 0 | 2 | 0 | 2,6 |
| 296 | gi\|221102513 | PREDICTED: S-adenosylmethionine synthase isoform type-1-like [Hydra vulgari | 69 | 42626 | 1 | 1 | 1 | 1 | 5,7 |
| 297 | gi\|828194686 | PREDICTED: alpha-mannosidase 2-like [Hydra vulgaris] | 69 | 130659 | 2 | 0 | 2 | 0 | 1,8 |
| 298 | gi\|5596622 | isovaleryl-CoA-dehydrogenase precursor [Arabidopsis thaliana] | 69 | 45371 | 1 | 1 | 1 | 1 | 2,9 |
| 299 | gi\|674263667 | glutathione s transferase mu [Echinococcus multilocularis] | 69 | 25815 | 3 | 0 | 2 | 0 | 4,1 |
| 300 | gi\|828202993 | PREDICTED: prelamin-A/C-like [Hydra vulgaris] | 68 | 68122 | 3 | 0 | 2 | 0 | 3,1 |
| 301 | gi\|490068496 | MULTISPECIES: hypothetical protein [Streptomyces] | 68 | 17530 | 2 | 1 | 1 | 1 | 7,6 |
| 302 | gi\|637068998 | replication initiation protein [Streptococcus suis] | 68 | 48271 | 2 | 0 | 2 | 0 | 3,2 |
| 303 | gi\|828195809 | PREDICTED: astacin-like metalloprotease toxin 5 [Hydra vulgaris] | 67 | 28949 | 2 | 0 | 2 | 0 | 7,1 |
| 304 | gi\|221131677 | PREDICTED: cytochrome b-c1 complex subunit Rieske, mitochondrial-like [Hy | 67 | 29671 | 3 | 0 | 2 | 0 | 4,1 |
| 305 | gi\|828226794 | PREDICTED: superoxide dismutase [Cu-Zn]-like [Hydra vulgaris] | 67 | 21118 | 1 | 1 | 1 | 1 | 6,3 |
| 306 | gi\|221102622 | PREDICTED: lactadherin-like [Hydra vulgaris] | 67 | 21699 | 2 | 0 | 2 | 0 | 10,2 |
| 307 | gi\|83595137 | mitochondrial phospholipid hydroperoxide glutathione peroxidase [Hydra vu | 67 | 21685 | 3 | 0 | 2 | 0 | 14,2 |
| 308 | gi\|493622316 | REX family transcriptional regulator [Pseudoflavonifractor capillosus] | 67 | 23378 | 1 | 1 | 1 | 1 | 3,7 |
| 309 | gi\|449683356 | PREDICTED: proliferation-associated protein 2G4-like [Hydra vulgaris] | 67 | 45036 | 2 | 0 | 2 | 0 | 5,5 |
| 310 | gi\|828231348 | PREDICTED: LOW QUALITY PROTEIN: transmembrane 9 superfamily member 2- | 67 | 76636 | 2 | 0 | 2 | 0 | 2,9 |
| 311 | gi\|88942082 | superoxide dismutase [Azumapecten farreri] | 66 | 15764 | 2 | 0 | 1 | 0 | 9,8 |
| 312 | gi\|828199555 | PREDICTED: microtubule-actin cross-linking factor 1-like, partial [Hydra vulgar | 66 | 777032 | 1 | 0 | 1 | 0 | 0,2 |
| 313 | gi\|221108650 | PREDICTED: serpin B6-like [Hydra vulgaris] | 65 | 43560 | 2 | 0 | 2 | 0 | 8,7 |
| 314 | gi\|449676978 | PREDICTED: heterogeneous nuclear ribonucleoprotein Q-like [Hydra vulgaris] | 65 | 68606 | 3 | 0 | 2 | 0 | 3,4 |
| 315 | gi\|828204809 | PREDICTED: branched-chain-amino-acid aminotransferase, cytosolic-like [Hyd | 65 | 38516 | 2 | 0 | 2 | 0 | 5,3 |
| 316 | gi\|828209870 | PREDICTED: hydroxyacid-oxoacid transhydrogenase, mitochondrial-like [Hydr | 65 | 51491 | 2 | 0 | 2 | 0 | 4,2 |

**Supplemental file 6: Table S3**

**Table S3.** LNA and RNA probe sequences used for WISH.

| Gene name | Accession Number | LNA or in situ probe sequence |
|---|---|---|
| HAS-1 | XP_012565441.1 | Full length antisense mRNA (1-717) |
| HAS-2 | XP_002162822.1 | ATCACGGTCAGGACGGCATTGT |
| HAS-3 | XP_002166229.3 | TAGTGACATATCTATCTCTGT |
| HAS-4 | XP_002162738.1 | ATTGTTCAGGTGTCAATTGTA |
| HAS-5 | XP_002164800.1 | TCAGACAAGTGTAGGTGTGATA |
| HAS-6 | XP_002157397.2 | TCTAAGGCAAGTGTAAGTGTGA |
| HAS-7 | XP_012560086.1 | Full length antisense mRNA (1-1021) |
| HAS-8 | XP_002153855.1 | TATGACGTAAGGTACAACAGCA |
| HAS-9 | XP_002161766.1 | ACGGCAAGATCTGCGGCAAGAT |
| HAS-10 | XP_002159980.2 | TACTGTACCAAGTCGCAAGCAA |
| HAS-11 | XP_012561076.1 | ACATGACTTGCAGCATAGCTGA |
| HMP-1 | NP_001296695.1 | Full length antisense mRNA (1-858) |

**Supplemental file 7: Table S4**

**Table S4.** siRNA and qPCR primer sequences.

| Gene name | siRNA Sequence (anti-sense) |
|---|---|
| HAS-7 | GUCUCCUUCAAACAGAUUGUUUU (siRNA1)<br>AAUGUUAUUCCAUAUAAUAUUUU (siRNA2)<br>UGAUUUGCAAUAACCUGUAUUUU (siRNA3) |
| HMP1 | UCACUGCAGAUAUGUAUGCUUUU (siRNA1)<br>UCCAGUGACACCGCUACACUUUU (siRNA2) |
| HAS-1 | ACUAUUGGAGUAUAGAGAUCUUU (siRNA1)<br>AUCGUAUGGAACAACAUUGUUUU (siRNA2) |
| HyDKK1/2/4 | GCAACGAAUGCAGCUACAACUUU (siRNA1)<br>UUUCGCAGUCUGCAUCCUUAUUU (siRNA2) |
| GFP | AAUUGGCCAUGGAACAGGUAGUUUU |
| Scrambled GFP | AAACCGGUGUGAAUCGAUGAGUUUU |
| HyWnt3 | AAATGGAGTTTCTATACAAAGUU |

| | qPCR Primer Sequence |
|---|---|
| EF1alpha | Fw: TATTGATAGACCTTTTCGACTTTGC<br>Rev: CTGTACAGAGCCACTTTCAACTTTT |
| HAS-7 | Fw: GGATGTGAAATCAAATGGTTATGCT<br>Rev: TGATGAACTCATTCTTCGAAGATCG |

#### Supplemental file 8: Supporting Information S1. Mathematical models and simulations

##### 1. Introduction of the mathematical model and its biological justification

The Gierer-Meinhard model of development of organs in *Hydra* influenced research on pattern formation in development [1, 2]. It also stirred controversy, since the nature of the hypothetical inhibitor has not been clarified as yet. The desired dynamics may alternatively result from a repressor in the intracellular signaling [3, 4] in *Hydra*, or a negative feedback loop stemming from mechano-chemical interactions [5]. In the current study, we propose to leave aside the controversy, and focus on new experimental findings specifically concerning Wnt3-HAS-7 interaction. We propose a mathematical model describing HAS-7 interactions with a coupled pattern formation system controlling the Wnt3 organizing center, the beta-Catenin/TCF body-scale patterning, and formation of tentacles. Each of the three subsystems is regulated by signaling feedbacks involving molecular components such as Wnt3 or beta-Catenin/TCF and their inhibitors. As noticed, the precise network of molecular interactions underlying the observed de novo formation of Wnt3 and beta-Catenin/TCF expression patterns has not been identified yet. Hence, the model includes an activator-inhibitor feedback loop, which fits the effects of the missing component.

##### Submodel of Wnt3 and beta-Catenin/TCF signaling

The core of the model accounts for the dynamics of Wnt3 and beta-Catenin/TCF signaling that are coupled through the canonical Wnt signaling pathway [6, 7]. Although it is then frequently assumed that beta-Catenin/TCF and Wnt3 molecules act in the confines of the same pattern formation system to coordinate body axis and head formation, e.g., [1, 8-10], we distinct between them in the model and describe dynamics of Wnt3 and dynamics of beta-Catenin using two model variables. The latter is motivated by experimental observations showing that Wnt3 and beta-Catenin/TCF act on distinct different spatial scales and may accomplish different developmental tasks. In contrast to beta-Catenin/TCF expression observed in diffusive patterns on the scale of the body axis, Wnt3 expression always appears as tiny spots with sharp boundaries in regions where beta-Catenin levels are sufficiently high (e.g. [7, 11-13]). In conclusion, Wnt3 expression centers define the head organizer region, a tiny structure at the tip of the hypostome, while the beta-Catenin patterns match the large-scale body axis gradient [7, 14, 15]. Distinction in the Wnt3 and the beta-Catenin/TCF pattern formation can e.g. be observed in an experiment following AZK treatment. A body-wide uniformly distributed increase of beta-Catenin/TCF does not change the size of Wnt3 spots but leads to an increase in the number of spots distributed over the whole body [10-12]. This observation contradicts the hypothesis proposed in Ref. [2] that Wnt3 expression patterns follow directly beta-Catenin patterns and are generated at the spots with beta-Catenin/TCF expression exceeding some threshold. In case of strong, uniform beta-Catenin upregulation such a threshold-mechanism would lead to larger Wnt3 spots, or, in an extreme case, to a body-wide constant Wnt3 expression. The latter observation indicates that the Wnt3 patterning undergoes independent nonlinear regulation that depending on the beta-Catenin concentration can result in a single spot (gradient-like) or multiple-spot (periodic or chaotic) structure. To model it, we assume that the organizer (Wnt3 spots) formation is driven by a self-contained pattern formation system acting on a small spatial scale, activated by high beta-Catenin levels. The model assumes a hypothetical Wnt3 inhibitor as it is the simplest mechanism generating such patterns. The recently discovered Sp5 transcription factor may play an important role in the transcriptional inhibition process of Wnt3 [10]. However, accounting for details of the molecular mechanism including Sp5 would require an additional model that is beyond the scope of this work. Additionally, following the previous models [2], we assume that the large-scale pattern formation system defining the body axis is shaped by beta-Catenin/TCF signaling with Dkk molecules possibly involved in the inhibition process. Dkk1/2/4-C has been shown to be positively regulated by the head-related positional value and to negatively regulate the canonical Wnt3 signaling [16, 17], which corresponds to generic features of the long-range inhibitor in the context of the activator-inhibitor model [2, 18]. Hence, we describe the beta-Catenin/TCF pattern formation applying a two-component activator-inhibitor model as a reduction of the signaling loop most probably involving more components [2, 10].

**Submodel for tentacle formation**

The model describing head and body axis formation is coupled to an additional subsystem describing tentacle formation. The latter receives information from the beta-Catenin/TCF signaling but does not feed back to the coupled Wnt3-beta-Catenin/TCF subsystem [2, 11, 19, 20]. Including the tentacle formation process in the model provides data for additional model verification, e.g., accounting for AZK treatment. The coupling is modeled using a source density concept. More specifically, we assume that the body axis gradient is recorded by a variable with a half-life distinctly longer than beta-Catenin/TCF. This assumption is based on the works of Gierer and Meinhardt and stay in agreement with several experimental observations [1, 2]. However, the molecular nature of the 'source density' is still unknown. In the model, we assume a simple feedforward from the beta-Catenin system [1, 2]. In turn, the source density activates the tentacle pattern formation system whereas the latter is inhibited by head-specific molecules (Wnt3). The assumed interactions among the body-axis system, the tentacles, and the source density allow for explaining a broad range of experimental observations (in detail presented and discussed e.g. in Ref. [2, 10]).

**Submodel of Wnt3-HAS-7 interactions**

The model is completed by including the HAS-7 function as discovered in this work. In particular, we assume that HAS-7 is positively regulated by beta-Catenin. The indirect regulation is modeled as a transcriptional HAS-7 activation downstream of the organizer by head-specific molecules (some candidates of the latter are presented in Ref. [13]). The assumption is motivated by (1) our HAS-7 promoter analysis, (2) the HAS-7 expression patterns after AZK treatment presented in this study, and (3) the natural mechanistic assumption that HAS-7 should be activated as soon as a head is established in order to suppress the formation of additional heads. Furthermore, we assume that HAS-7 degrades Wnt3 ligands as demonstrated in this work, and that Wnt3 negatively regulates HAS-7. The latter is motivated by (1) our observation that HAS-7 transcript is absent from the upper hypostome, and (2) the natural mechanistic assumption that HAS-7 suppresses organizer formation in the surroundings but not of the existing organizer itself. It is not known if the local negative regulation of HAS-7 by the hypostome is governed directly by Wnt3 or by other molecules downstream of Wnt3. Nevertheless, accounting for such modification of the regulatory mechanism does not influence the results of the model. Hence, for the sake of simplicity, we assume a local negative regulation of HAS-7 by Wnt3.

#### Evolving geometry of the model

To model a realistic geometry of the Hydra tissue bilayer, we adopt the mechano-chemical modeling approach proposed in Ref. [21] for a membrane deformation and extended to describe small deformations of a thin tissue in Ref. [5]. We assume that chemical reaction-diffusion equations are defined on a curved 2-dimensional surface that is embedded in a 3-dimensional space. The geometry of the curved surface evolves in time following a gradient-flow of a Helfrich-type energy reflecting the property that bending of the tissue away from a preferred local curvature is energetically unfavorable. In contrast to the fully coupled mechano-chemical model of [5], in the current model we do not consider any feedback from mechanical properties of the tissue to the gene expression processes. Consequently, the mechanical part of the model does control the pattern formation mechanism that is purely based on molecular interactions but provides a realistic description of the tissue described as a radial-symmetric ellipsoid that undergoes small deformations due to the gene expression patterns.

#### 2. Model equations and simulations

##### General structure of the mathematical model

The above-mentioned processes are translated to a mathematical model given in terms of a system of partial differential equations (PDEs). In particular, we apply a continuous modeling approach that is justified by the large number of cells ($\geq 10^4$) in the system [25]. The cell bilayer forming a hollow tissue ellipsoid is at any time $t$ approximated by a closed 2D surface $\Gamma(t)$, embedded in a 3D space. The evolution of $\Gamma(t)$ is given by a diffeomorphic time-dependent representation $\dot{X}$, parameterized over the unit sphere $S^2 \subset \mathrm{R}^3$. Thus, $\Gamma(t)$ is the image of $\dot{X}(\cdot, t)$ with $\dot{X}(\dot{s}, t)$: $S^2 \times [0, T] \rightarrow \mathbb{R}^3$ for a $T \in \mathbb{R}_{>0}$. Local concentrations of different gene products $a$ at time $t$ are given by continuous functions $\Phi_a$ on the deforming tissue surface $\Gamma(t)$, defined as gene product concentrations per cell volume, $\Phi_a(t)$: $\Gamma(t) \rightarrow \mathbb{R}_{\geq 0}$. In order to achieve a consistent formulation with chemical processes being defined on $S^2$ rather than on $\Gamma$, we redefine $\Phi_a$ identifying material points $\dot{X}(\dot{s}, t)$ on $\Gamma(t)$ with $\dot{s} \in S^2$. The latter is ensured since $\dot{X}$ is smooth and bijective. Thus, for each $\dot{s} \in S^2$, $t \in [0, T]$, we define the function $\phi_a : S^2 \times [0, T] \rightarrow \mathbb{R}_{\geq 0}$ by $\phi_a(\dot{s}, t) = \Phi_a(\dot{X}(\dot{s}, t))$.

##### Model of molecular interactions

As motivated above, the model accounts for spatio-temporal dynamics of nine different chemical components (Table 1). In particular, model variables *β_cat* and $\beta\_cat_{ant}$ denote the activator and the inhibitor associated with the body-axis patterning system involving nuclear beta-Catenin/TCF. Furthermore, *Wnt*3 and $Wnt3_{ant}$ describe the small-scale Wnt3 organizer pattern formation system, *Head* denotes a fast diffusing head-specific gene product downstream of the organizer (such as one of the multiple head-specific Wnts [22]), *HAS* describes HAS-7 dynamics, *SD* represents source density, and *Tent* and $Tent_{ant}$ model the tentacle system. The resulting reaction-diffusion system defined on the domain corresponding to the tissue is given in Eq. (1)-(9). The process of spatial movement of the molecules is described by a surface Laplace (Laplace-Beltrami-) operator $\Delta^{\Gamma}(\cdot)$. Eq. (1)-(2) are an extension of the classical Gierer-Meinhardt model [1, 2]. In particular, we model production of beta-Catenin/TCF as a function of *Wnt*3. This assumption is derived from canonical Wnt signaling [6] and reflects also the experimental observation that transplantation of the organizer induces a secondary body axis in Hydra (e.g., [11, 23]). Eq. (3)-(4) describe the Wnt3-related small-scale pattern formation. Additional to the classical interactions model [1, 2], we take into account the activation of the Wnt3 by beta-Catenin system, which is reflecting experimental observations [12]. We model it using a function that depends on $\beta\_cat_{ant}$. However, the latter dependence can be replaced by a function depending on $\beta\_cat$, since both variables express a similar spatial pattern. Thus, using $\beta\_cat_{ant}$ instead of $\beta\_cat$ is an arbitrary choice here. Furthermore, the parameters $b_3$ and $b_4$ control above which $\beta\_cat_{ant}$ threshold Wnt3 patterning is locally activated. Since in siHAS-7 knockdown experiments, in < 50 % of polyps a secondary axis is induced if not treated with AZK, whereas after additional AZK treatment, approx. 90 % of the polyps develop a secondary axis, $b_3$ and $b_4$ have been adjusted such that only in simulations of siHAS-7 + AZK experiments a secondary axis develops. Eq. (5) is a simple reaction-diffusion equation with *Head*-production depending linearly on *Wnt*3 and *HAS* production (c.f. Eq. (6)) depending linearly on *Head* and reduced by *Wnt*3. Finally, Eq. (7)-(8) describing the tentacle system Eq. (9) for the source density are adopted from Ref. [1, 2].

Table 1. Model variables and their biological meaning.

| **Variable name** | **Explanation** |
|---|---|
| *β_cat* | Nuclear beta-Catenin/TCF |
| *β_cat*$_{ant}$ | *β_cat* antagonist, probably involving Dickkopf1/2/4-C [2, 16] HyWnt3 |
| *Wnt*3 | HyWnt3 |
| *Wnt*3$_{ant}$ | HyWnt3 antagonist (probably Sp5 [10] and HmTSP [24] involved) |
| *Head* | Head-related factors downstream of HyWnt3 (e.g., multiple Wnts [13]) |
| *HAS* | HAS-7 |
| *SD* | Source density (long-term storage of the head forming potential) |
| *Tent* | Tentacle activator (probably HyAlx [19], Wnt8 [22], BMB5-8b [25] involved) |
| *Tent*$_{ant}$ | Tentacle activator antagonist (unknown) |

$$
\begin{aligned}
d_t\beta\_cat &= a_1\Delta^\Gamma\beta\_cat + b_1\cdot(1.0+c_1\cdot Wnt3)\cdot SD\cdot\frac{0.05+\beta_{cat}^2}{\beta\_cat_{ant}} - d_1\cdot\beta\_cat && (1)\\
d_t\,\beta\_cat_{ant} &= a_2\Delta^\Gamma\beta\_cat_{ant} + b_2\cdot(1.0+c_2\cdot Wnt3)\cdot SD\cdot\beta_{cat}^2 - d_2\cdot\beta\_cat_{ant} && (2)\\
d_tWnt3 &= a_3\Delta^\Gamma Wnt3 + b_3\cdot\beta\_cat_{ant}\cdot\frac{0.005+Wnt3^2}{Wnt3_{ant}\cdot(1.0+c_3\cdot Wnt3^2)}\\
&\quad -d_3\cdot(1.0+e_3\cdot HAS)\cdot Wnt3 && (3)\\
d_tWnt3_{ant} &= a_4\Delta^\Gamma Wnt3_{ant} + b_4\cdot\beta\_cat_{ant}\cdot\frac{0.005+Wnt3^2}{1.0+c_4\cdot Wnt3^2} + 0.035\\
&\quad -d_4\cdot Wnt3_{ant} && (4)\\
d_tHead &= a_5\Delta^\Gamma Head + b_5Wnt3 - d_5Head && (5)\\
d_tHAS &= a_6\Delta^\Gamma HAS + \frac{b_6Head}{1.0+c_6\cdot Wnt3} - d_6HAS && (6)\\
d_tTent &= a_7\Delta^\Gamma Tent + \frac{b_7\cdot SD\cdot(0.005+Tent^2)}{Tent_{ant}\cdot(1+c_7\cdot Tent^2)\cdot(1+e_7\cdot Head_{ant})} - d_7\cdot Tent && (7)\\
d_tTent_{ant} &= a_8\Delta^\Gamma Tent_{ant} + \frac{b_8\cdot SD\cdot(0.005+Tent^2)}{(1+c_8\cdot Tent^2)\cdot(1+e_8\cdot Head_{ant})} + 0.014 - d_8\cdot Tent_{ant} && (8)\\
d_t\,SD &= a_9\Delta^\Gamma SD + b_9\cdot\beta\_cat + 0.00003 - d_9SD. && (9)
\end{aligned}
$$

Since calibration of the model using current experimental data is not possible and the focus of this research is in qualitative system behavior (such as model ability to reproduce specific patterns of gene expression), we performed numerical simulations of the model to obtain insights into dependence of the model results on specific choices of parameters. Our study suggests that most of the parameters involved in Eq. (1)-(9) do not influence critically the qualitative HAS-related simulation results as presented within the main manuscript. In particular, most of them control specific properties of one of the three interplaying *de novo* pattern formation systems, such as spatial scaling of the pattern (size of expression domain), spacing between the maxima of the pattern, or a condition for *de novo* patterning. Following these simulation results, we fix most of parameters to values taken from Ref. [1, 2].

This allows focusing on parameters accounting for the novel aspects of the model such as (A) interactions between beta-Catenin and Wnt3 that govern pattern formation on different spatial scales, and (B) feedback loop between HAS-7 and Wnt3. In general, changing the corresponding parameters, we observe robust model dynamics (qualitatively the same pattern formation). The only discrepancy is observed in simulations of AZP + siHAS-7 animals, where the number of ectopic axes in model simulations depends on the strength of HAS-7 dependent degradation of Wnt3. This observation suggests that there might be an additional mechanism (possibly involving other members of the HAS family) ensuring an experimentally observed robustness with respect to the number of organizers.

##### Model of tissue mechanics

The chemical equations Eq. (1)-(9) are augmented by a set of equations representing the deforming tissue surface. In particular, we treat the tissue as purely elastic, and elastic tissue deformations are based on minimization of the Helfrich free energy [26], which is given by

$$F_{bend} = \kappa \int (H - H_0 (\beta\_cat, Tent))^2 \, dS$$

Here, $H$ is the mean curvature, $\kappa$ the bending rigidity, and $H_0$ the spontaneous curvature [21]. In particular, $H_0$ represents the locally preferred tissue curvature, which again may depend on local morphogen concentrations. Namely here, we assume $H_0(\beta\text{-}cat_{ant}, Tent) = 0.4 \cdot \beta\_cat_{ant} + 5. \cdot Tent$, based on the observation that local tissue evaginations can be observed during both – budding and tentacle formation [22]. Local area-conserving evolution of the deforming *Hydra* tissue is finally given by the $L^2$-gradient flow of the total energy. For further details, we refer to Ref. [21].

##### Numerical implementation

The mathematical model is simulated using the finite element library Gascoigne [27]. It is based on approximation of the fourth order PDEs in a mixed formulation. For spatial discretization, we apply linear finite elements; for time discretization a semi-implicit Euler scheme. For further details of the computation scheme we refer to [21].

**Parameters and initial conditions**

For simulations of the unperturbed system, we apply the following parameters following Ref. model [1, 2]):

$a_1 = 9 \times 10^{-5}$, $b_1 = 3 \times 10^{-3}$, $c_1 = 3 \times 10^{-2}$, $d_1 = 3 \times 10^{-3}$,

$a_2 = 11 \times 10^{-2}$, $b_2 = 3 \times 10^{-3}$, $c_2 = 3 \times 10^{-2}$, $d_2 = 4 \times 10^{-3}$,

$a_3 = 6 \times 10^{-5}$, $b_3 = 7 \times 10^{-3}$, $c_3 = 3 \times 10^{-3}$, $d_3 = 12 \times 10^{-2}$, $e_3 = 1 \times 10^{2}$,

$a_4 = 24 \times 10^{-3}$, $b_4 = 1 \times 10^{-2}$, $c_4 = 3 \times 10^{-3}$, $d_4 = 18 \times 10^{-2}$,

$a_5 = 25 \times 10^{-3}$, $b_5 = 1 \times 10^{0}$, $d_5 = 1 \times 10^{-2}$,

$a_6 = 25 \times 10^{-3}$, $b_6 = 1 \times 10^{-1}$, $c_6 = 1 \times 10^{1}$, $d_6 = 5 \times 10^{-2}$,

$a_7 = 25 \times 10^{-5}$, $b_7 = 2 \times 10^{-3}$, $c_7 = 12 \times 10^{-2}$, $e_7 = 3 \times 10^{-2}$, $d_7 = 2 \times 10^{-2}$,

$a_8 = 27 \times 10^{-3}$, $b_8 = 3 \times 10^{-3}$, $c_8 = 12 \times 10^{-2}$, $e_8 = 3 \times 10^{-2}$, $d_8 = 3 \times 10^{-2}$,

$a_9 = 11 \times 1^{-5}$, $b_9 = 3 \times 10^{-5}$, $d_9 = 3 \times 10^{-5}$.

To approximate the geometry of the Hydra tissue, initial conditions for $X_1$, $X_2$ and $X_3$ are parametrized over a closed 2D unit-sphere $S^2$ embedded in 3D space with $X_1(t = 0) \equiv X_2(t = 0) \equiv 0$ and $X_3(t = 0) = 4 \cdot s_3$, thus, leading to a stretch into the direction of $s_3$ (given that $s_1$, $s_2$, $s_3$ are Eulerian coordinates of the $S^2$-surface). For biological molecules, we use a stochastic initial distribution based on the standard random generator provided by C++. The source density is modeled using an initial gradient given by $SD(t = 0) = 4.0 \cdot (exp(s_3)/exp(1))$. Thus, in all simulations, only the geometric and chemical body axis gradient are initially prescribed.

In simulation of the AZK treatment, we modify the initial conditions for the source density adding an offset by $SD(t = 0) = 2.0 + 4.0 \cdot (exp(s_3)/exp(1))$. To simulate HAS knockdown, we set $b_6 = 0$. For Dkk knockdown, we model a reduction of Dkk activity by increasing $d_2$ by the two-fold. In the model, a complete deactivation of $beta\_cat_{ant}$ prevents creation of any pattern, since the body-scale system is described by just two components that are a minimal set required for pattern formation. Finally, Wnt3 overexpression is simulated by adding the constant $c = 0.1$ to the production.

##### Supplemental Information S1 References
